# A non-autonomous protein quality control mechanism targeting tau aggregate propagation

**DOI:** 10.1101/2024.07.26.605305

**Authors:** Anika Bluemke, Birte Hagemeier, Kamilla Ripkens, Nina Schulze, Michal Strzala, Michelle Koci, Farnusch Kaschani, Markus Kaiser, Michael Erkelenz, Sebastian Schluecker, Melisa Merdanovic, Simon Poepsel, Doris Hellerschmied, Steve Burston, Michael Ehrmann

## Abstract

Tauopathies such as Alzheimer’s disease, frontotemporal dementia with Parkinsonism, and other neurodegenerative disorders are characterized by the spread of tau pathology from an initial brain region to neuroanatomically connected areas. At the molecular level, spreading involves aggregation of tau in a donor cell, externalization of transmissible fragments of amyloid fibrils, internalization by an acceptor cell, followed by seeded aggregation of endogenous tau. However, the protein quality control mechanisms that counteract tau aggregation, and in particular its spreading process, are not well understood. In this context, a co-migrating factor performing location-independent interference of fibril formation and transmission would be an appropriate conceptual solution. Here, we show that the cell-to-cell transfer of the widely conserved serine protease HTRA1 impedes tau pathology by targeting multiple steps within the spreading process. Our results suggest a defense mechanism against the intercellular spread of pathogenic protein conformations.

## INTRODUCTION

Amyloid fibrils are the hallmark of prominent neurodegenerative diseases. They are characterized by tightly packed β-sheets, in which regions of identical amino acid sequences are stacked on top of each other (1). Alzheimer’s disease (AD) is characterized by the presence of amyloid fibrils composed of extracellular Aβ peptides and of intracellular tau protein (microtubule associated protein tau, MAPT). Tau dissociates from microtubules upon hyperphosphorylation before forming fibrillar structures (2, 3). The core of amyloid tau fibrils is formed by residues 306–378 (in the numbering of the 441-residue human tau isoform), corresponding to the microtubule-binding repeats R3 and R4, and some amino acids following R4, while the rest of the protein forms the so-called fuzzy coat (4). Disease progression is caused by the spread of fibrils from one brain region to connected areas, accompanied by cognitive impairment. The individual steps of propagation are initiation of aggregation in a donor neuron, externalization of a transmissible tau species, intercellular transfer, internalization in a recipient cell, followed by seeding with endogenous tau (5). Seeding is a very efficient process not only when carried out with purified proteins but also in cells where a doubling time of self-replication of small tau aggregates can be 5 h in HEK293 cells or one day in primary neurons (6).

From a protein quality control perspective, tau aggregates can be considered a moving target, requiring either dedicated factors in different cellular compartments or factors that co-migrate with the transmissible species. The secreted serine protease HTRA1 is a member of the high temperature requirement A (HtrA) family implicated in protein quality control (7). HTRA1 senses protein folding stress by binding to exposed C-termini of misfolded, fragmented and mislocalized proteins via its C-terminal PDZ domain. Despite being a secreted protein, HTRA1 is also found in the cytosol (8, 9). HTRA1 has been implicated in severe pathologies such as age-related macular degeneration, arthritis, cancer and familial ischemic cerebral small vessel disease (7). In the context of neurodegenerative diseases, HTRA1 degrades tau fibrils by applying a unique mechanism in which fibril dissociation precedes proteolysis, suggesting its role in tau pathology (10). In human AD brain extracts, levels of neurofibrillary tangles and HTRA1 are negatively correlated (11). Consistently, a proteomic study that investigated human brain samples from AD patients found a strong association of HTRA1 with detergent-insoluble tau (12). In addition, colocalization of HTRA1 and neurofibrillary tangles was demonstrated (13). Here, we show that the cell-to-cell transfer of the widely conserved serine protease HTRA1 inhibits tau spreading by targeting multiple steps within this process. We also provide quantitative insights at high temporal and spatial resolution into how HTRA1 degrades tau fibrils.

## RESULTS

### **HTRA1 reduces tau seeding** *in vitro*

Since HTRA1 is able to digest soluble and fibrillar tau, the latter by combining fibril dissociation and proteolysis (10, 11), we hypothesized that this secreted protein quality control factor might also be involved in the defense against intercellular spreading of aggregated tau. To initially address this question, and to obtain direct biochemical evidence confirming that HTRA1 is indeed acting on seeds, we performed experiments with purified proteins. Seeded aggregation of tau was modeled *in vitro* by co-incubating soluble tau with seeds. Proteolytically inactive HTRA1, where the catalytic Ser328 residue is exchanged by Ala (HTRA1^S328A^) was pre-incubated with tau seeds to induce their disassembly, while proteolytically active HTRA1 was added directly to soluble tau and seeds. Subsequently, a sedimentation assay was performed to distinguish between soluble tau in the supernatant and aggregated tau in the pellet fraction (Fig. 1A). The total protein load indicated identical amounts of tau. Soluble tau alone formed fibrils, but upon addition of seeds, 1.8-fold more tau was found in the pellet fraction (Fig. 1A). To assess the amount of fibrillar tau, the pellet fraction was subjected to negative-stain transmission electron microscopy and the total fibril length was calculated by adding the lengths of individual fibrils, as previously established (10). Upon addition of tau seeds, the total fibril length was increased 1.8-fold (Fig. 1B), in line with seeded aggregation. HTRA1^S328A^ caused a concentration-dependent shift of tau from the pellet to the supernatant fraction. At the highest concentration of HTRA1^S328A^, 88% of the tau protein was found in the supernatant, while HTRA1^S328A^ was predominantly soluble. At the highest concentration of HTRA1^S328A^, the total fibril length was reduced to less than 50% as compared to the control. Tau polymerization was also monitored by measuring ThT fluorescence (SI Appendix, Fig. S1A and B). Treatment with HTRA1^S328A^ resulted in consistently low ThT fluorescence, indicating efficient clearance of seeds associated with a reduced nucleation of tau aggregates. In its proteolytically active form, HTRA1 digested tau. Proteolytic products did not pellet upon ultracentrifugation, indicating that HTRA1-mediated degradation of tau did not increase its aggregation propensity. Consistently, increasing HTRA1 amounts reduced total fibril length by 86% and decreased ThT fluorescence to background levels (SI Appendix, Fig. S1B).

**Fig. 1.**
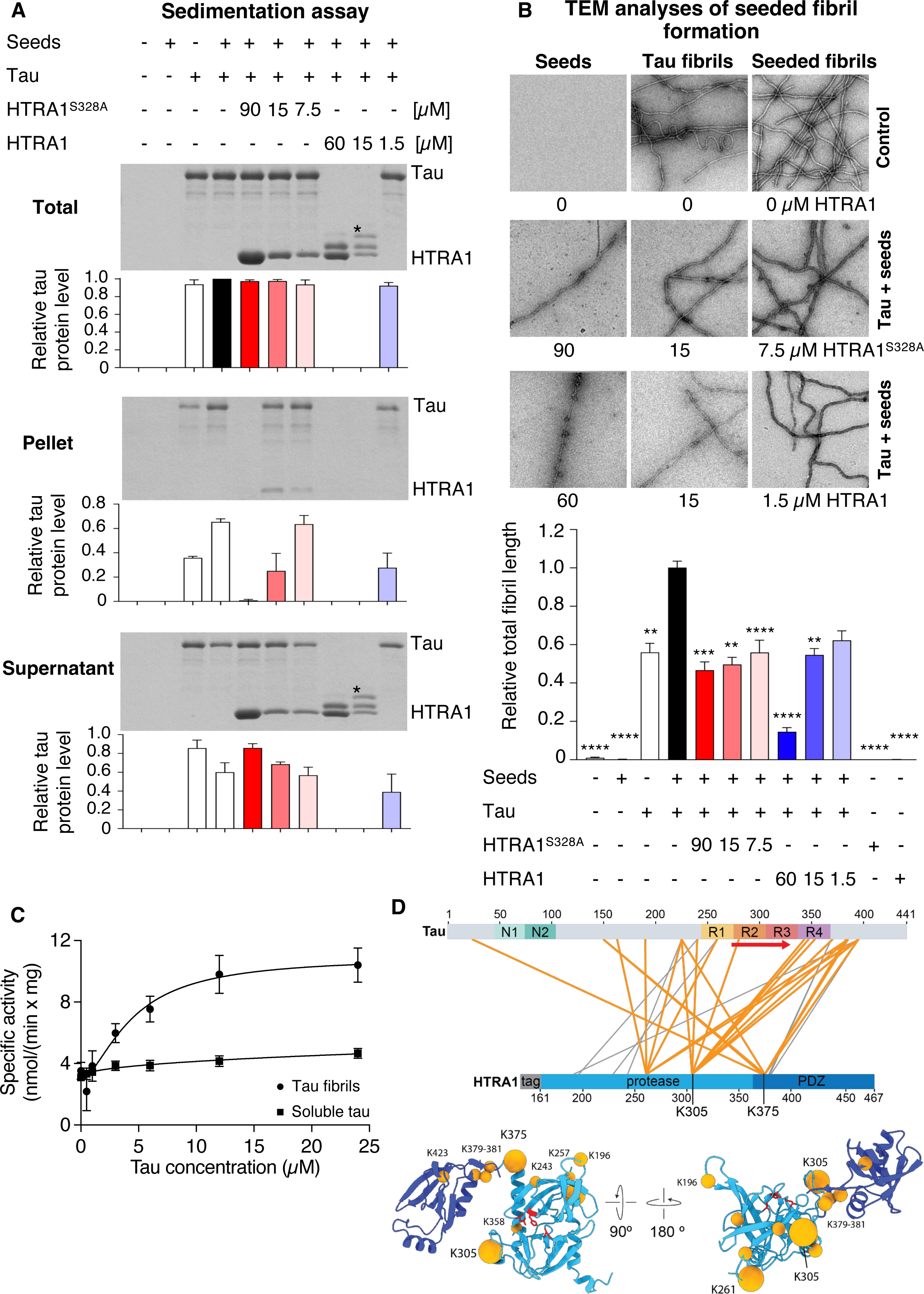
HTRA1 reduces seeded tau aggregation independent of its proteolytic activity. Soluble tau (30 µM) was incubated with tau seeds (0.6 µM) in aggregation buffer for 3 days at 37°C. Proteolytically active HTRA1 (3, 15, 60 µM) or inactive HTRA1^S328A^ (7.5, 15, 90 µM) was applied to study the solubilization and degradation of tau fibrils. For solubilization, seeds were preincubated with HTRA1^S328A^, while HTRA1 was added directly to soluble tau and seeds. (A) Sedimentation assay. Samples were subjected to ultracentrifugation. Total, pellet and supernatant fractions were subjected to SDS-PAGE and Coomassie staining. Bands marked with an asterisk mark proteolytic products of tau. Tau protein levels were quantified by densitometry. Values are normalized to the total amount of tau co-incubated with seeds (black bar at total tau data). Error bars, s.e.m.; n = 3. (B) Samples from (A) were further analyzed by transmission electron microscopy (TEM). Quantification of the total fibril length per image is based on at least 25 images (7.89 µm x 7.89 µm) per condition. Co-incubation of tau and seeds (black bar) was used for the normalization of values and the analysis of significant differences. Error bars, s.e.m.; n = 2. ** P < 0.01, *** P < 0.001, **** P < 0.0001, Kruskal-Wallis test with Dunn’s post-test. Representative TEM images. Scale bar: 250 nm. (C) Activation of HTRA1 by tau fibrils. The specific activity of purified HTRA1 in the presence of soluble (■) and fibrillar (•) Tau was determined using the synthetic substrate VFNTLPMMGKASPV-pNA. HTRA1 was mixed with pNA substrate and the tau concentrations indicated. Proteolysis of the pNA substrate was continuously measured at 405 nm and used to calculate the specific enzyme activity. n=3. (D) Cross-linking mass spectrometry analysis of HTRA1 S328A binding to tau fibrils. Schematic representation of intermolecular cross-links between tau and HTRA1. Cross-links involving the most frequently cross-linked residues of HTRA1 (K261, K305 and K375) are highlighted in orange. N1, N2: N-terminal inserts, R1-4: microtubule-binding repeat regions of tau. Red arrow: β-sheet region of tau fibrils (PDB ID: 6QIH). HTRA1 monomer model including the protease (light blue) and PDZ domains (dark blue). Orange spheres mark the HTRA1 residues with intermolecular cross-links to tau. Most frequently cross-linked residues are highlighted by larger spheres. Catalytic residues of the active site are shown in red and in ball-and-stick representation. Model based on an alphafold2 prediction. Note that the PDZ domain displays flexibility relative to the protease domain and is not in a fixed position.

### Activation of HTRA1 by tau fibrils and identification of binding sites

HTRA1 is regulated by the mechanism of ligand-induced activation. Such ligands can be peptides or folded proteins (14, 15). To test whether tau fibrils activate HTRA1, protease assays of recombinant HTRA1 were performed using the artificial peptidic substrate VFNTLPMMGKASPV-pNA (16) in the presence of soluble or fibrillar tau. HTRA1 was up to >2.5-fold more active in the presence of tau fibrils but not soluble tau (Fig. 1C). These data suggest that tau fibrils interact more tightly with the active site of HTRA1 than soluble tau and that the activation by fibrils supports their efficient degradation.

To investigate how HTRA1 engages with tau fibrils, we performed cross-linking mass spectrometry using the inactive HTRA1^S328A^ mutant, tau fibril seeds, and the cross- linking reagent PhoX (3,5-Bis(((2,5-dioxopyrrolidin-1- yl)oxy)carbonyl)phenyl)phosphonic acid (17). Mass spectrometry identified three amino acids within HTRA1 that extensively cross-linked to tau: K305 in the sensor loop L3, K375 located at the border of the protease and PDZ domains, and K261 located in a loop opposite the active site. Further cross-links clustered at the connecting region between protease and PDZ domains, including the N-terminal part of the PDZ domain as well as lysines at the peripheral face of the HTRA1 trimer (Fig. 1D). Functionally, K305 located in the sensor loop L3 is of interest since it is a key element of the activation domain of HTRA1, which allows for the ligand-induced activation of the proteolytic activity by triggering a disorder-to-order transition of the active site (16). Cross-links near the PDZ domain and on the peripheral surface of the protease domain indicate how HTRA1S328A initially interacts with tau fibrils. Interestingly, the amino acids of tau found to cross-link to these regions of HTRA1 are located primarily in the C-terminal portion, flanking the β-sheet-forming fibrillar core of tau (Fig. 1D). No cross- links were detected closer to the active site of HTRA1, which could be explained by the tight binding of tau near the active site potentially excluding the cross-linking reagent from access to reactive side chains. Lys residues, which preferentially react with PhoX, are unlikely cleavage sites because HTRA1 has a preference for small hydrophobic residues (16). However, tau residues K353, K385 and K395, which cross- linked with the L3 loop of HTRA1, are in close proximity to residues L376, I392 and L408, which are cleaved early and very efficiently (see below for details).

### High-resolution analysis of proteolytic processing of soluble tau and tau fibrils

To better understand the dynamics of tau fibril proteolysis by HTRA1, we used time-resolved mass spectrometry (MS) in combination with bioinformatic analyses to identify proteolytic products. The sequences of the identified proteolytic products are aligned along the primary amino acid sequence of the substrate, and the relative frequency of cuts at each cleavage site is calculated (Fig. 2A, SI Appendix, Fig. S2, Supplemental data 1, 3, 4).

**Fig. 2.**
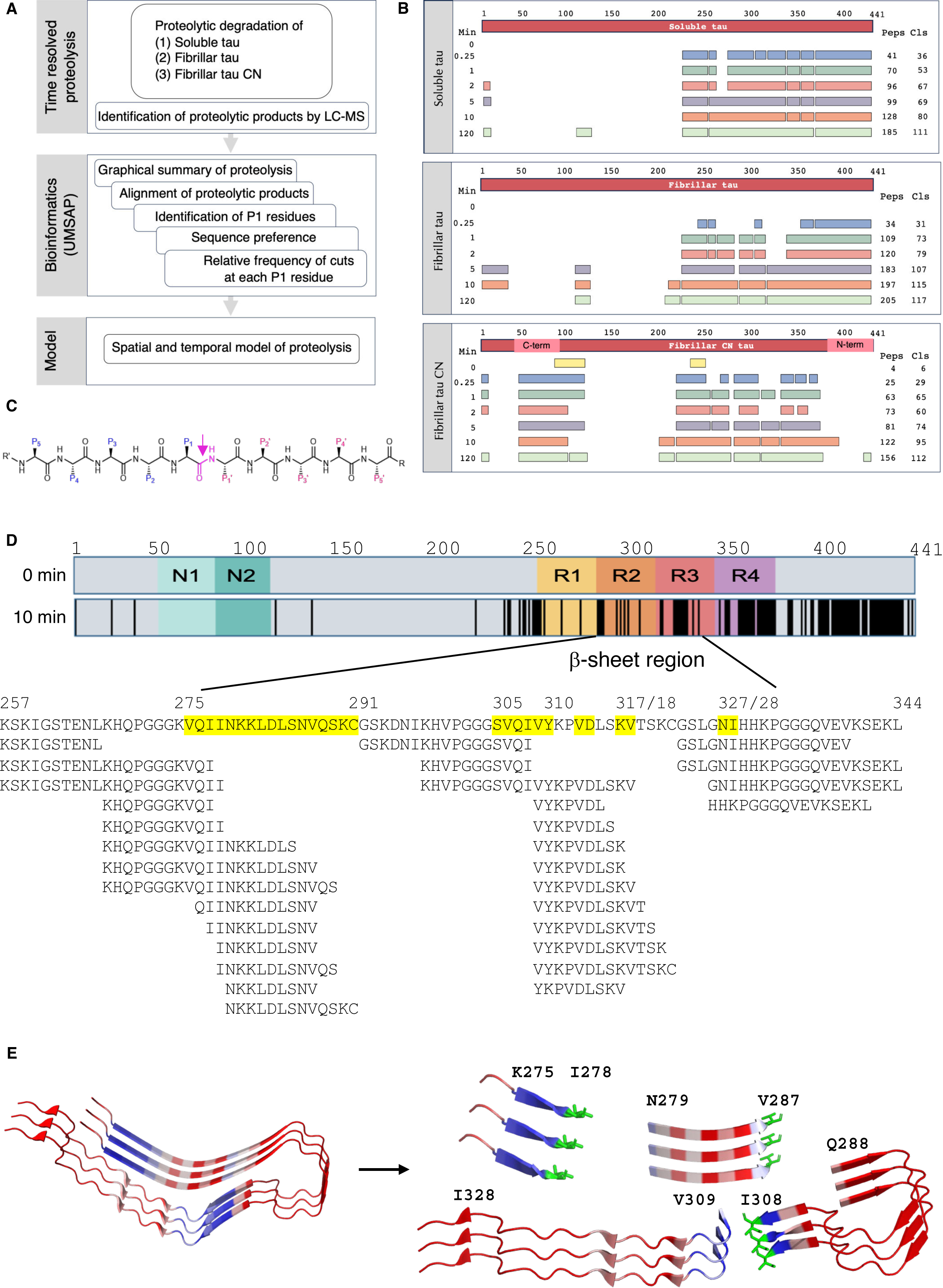
**Proteolytic degradation of soluble and fibrillar tau by HTRA1.** (A) Workflow, see text for details. (B) Time-resolved proteolysis of soluble and fibrillar forms of tau by HTRA1. Top bar: linear representation of the entire substrate protein. For each of the indicated time points (Min), peptide sequences that align without gaps are grouped into fragments, shown as bars. The number of identified peptides (Peps) and the total number of cleavage sites (Cls) are shown at the right. Identified peptides aligned to the primary amino acid sequence of tau are shown in Supplemental data 1 (soluble tau), 3 (fibrillar tau) and 4 (fibrillar tau CN, where parts of the N- and C-termini have been swapped). (C) Model peptide and standard nomenclature. The residue of the scissile bond (magenta) is termed P1, which is often the major determinant of substrate specificity. Residues upstream to P1 are termed P2, P3 etc. Residues located downstream to P1 are termed P1’, P2’, P3’ etc.(47). The proteolytic cleavage site is marked by an arrow. (D) Top bar, graphic representation of 2N4R tau. The conserved N-domains and repeat regions are highlighted. Lower bar, vertical black lines indicate P1 residues that are cleaved after 10 min of incubation with HTRA1. The β-sheet region of fibrils is highlighted, numbers indicate the amino acids present as β-sheets. Amino acid sequence alignment covering residues K257-L344, β-sheet forming residues are highlighted in yellow. Peptides below the primary amino acid sequence correspond to proteolytic products identified by MS. (E) Early cleavage events in fibrillar tau. Following cuts at major proteolytic sites, the fibrillar core (left, PDB ID: 6QJH) is initially processed into 4 fragments (right). Key P1 residues I278, V287 and I308 (green) are shown as sticks. See text for details.

Before studying the proteolysis of tau fibrils, we examined the degradation of soluble tau (18). Soluble tau was incubated with HTRA1 and samples were taken at seven time points within 120 min. LC-MS data were analyzed using UMSAP software (19) to identify proteolytic products and the number of cleavage sites at each time point (Fig. 2B). These data identified 36 sites after 15 sec and 111 sites after 120 min of incubation. Surprisingly, the identified peptides did not span the entire sequence even after 120 min, as the only four cleavage sites that were detected in the N-terminal half (residues 1-225) were identified only once at late time points (Fig. 2B, SI Appendix, Fig. S2A, Supplemental data 1). The downstream region from residues 226-274 contained 11 cleavage sites, six of which were cleaved only once after 120 min of incubation. In contrast, the remaining C-terminal region contained the seven most frequently detected cleavage sites and was thus most efficiently cleaved (SI Appendix, Supplemental text, Fig. S2A, Tables S1, S2).

### Proteolysis of fibrillar tau by HTRA1

For tau fibrils, time-resolved MS revealed 31 cleavage sites after 15 sec and 117 sites after 120 min of incubation (Fig. 2B). Already after 1 min of incubation, the entire β- sheet-forming portion of tau was processed into 12 peptides (SI Appendix, Supplemental data 3), after 10 min into 28 peptides (Fig. 2D), and after 120 min into 31 peptides (SI Appendix, Supplemental data 3).

The P1 sites with the highest relative number of cuts at the earliest timepoints are candidate sites where the proteolysis of fibrils is initiated. These sites are L408 and A392, followed by L376 and L344 located downstream to the fibrillar core. Upstream of the fibrillar core, HTRA1 cleaves after V256 and subsequently after L266, which is located 9 residues upstream of the fibril core. Together, these cuts produce a fragment of K267-L344, containing the entire fibril core. Subsequently, K267-L344 is fragmented by cuts after I308 (in β-sheet 2) and I278 and V287 (in β-sheet 1) into 4 peptides containing 9, 9, 20 and 35 residues respectively, before these peptides are successively degraded into small peptides of a minimum size between 7 and 17 residues (Fig. 2D and E, SI Appendix, Fig. S2B, Supplemental data 3). Likewise, the region downstream of the fibrillar core is degraded into small peptides by cuts at 74 sites (SI Appendix, Fig. S2B, Supplemental data 3).

As in the case of soluble tau, the identified peptides did not cover the entire sequence (Fig. 2B, SI Appendix, Fig. S2B, Supplemental data 3), supporting the notion that the N-terminus of tau up until residue 225 is a poor substrate of HTRA1. In addition, we observed further common features for soluble and fibrillar substrates. The appearance of many products sharing only one identical cleavage site suggests that these products are the result of one high-affinity and several low-affinity binding sites (SI Appendix, Supplemental data 1, 3, 4). This model is supported by the observed tendency for longer proteolytic fragments, again sharing an identical cleavage site, to appear preferentially at later time points, suggesting that these additional cleavage sites are of even lower affinity. A comparison of the similarities and differences of cleavage efficiency at particular sites points to areas that are differently accessible in soluble and fibrillar tau (SI Appendix, Supplemental text, Tables S1 and S2). Different cleavage efficiencies are mainly detected for residues in the fibril core region that are cleaved more frequently in soluble compared to fibrillar tau (SI Appendix, Table S3), as expected due to the tight packing of the fibril core.

### Proteolysis of tau derivatives with swapped N- and C-termini

To address the curiosity of the unexpected protease resistance of the N-terminus and the pronounced protease sensitivity of the C-terminus of tau, we swapped the C-terminal residues 391-441 with the N-terminal residues 42-98 to generate the tau CN construct. Tau CN fibrils were incubated with HTRA1 and the resulting cleavage products were analyzed as described above. 29 cleavage sites were detected after 15 sec and 112 sites after 120 min of incubation (Fig. 2B), which is comparable to the digests of wt tau fibrils (Fig. 2B). Notably, the cleavage pattern of the swapped C-terminal and N-terminal parts of tau CN did only slightly or not change compared to wt tau fibrils (SI Appendix, Fig. S2C, Supplemental data 4). These data support the notion that the N-terminal part of tau, because of its amino acid sequence rather than position within the polypeptide, is a poor substrate for HTRA1. In addition, swapping the N- and C-terminal fragments had only minor effects on the cleavage pattern of the β-sheet region (SI Appendix, Fig. S2C, Supplemental data 4). Also, the calculation of the amino acid distribution at the five positions preceding or succeeding the scissile bond did not show major changes in the three substrates analyzed (SI Appendix, Fig. S2D). However, the relative numbers of cuts at each cleavage site and timepoint were in general lower for tau CN fibrils compared to the digests of soluble and wt fibrillar tau, suggesting that tau CN fibrils were digested more slowly. Therefore, future work is needed to determine the molecular basis of the protease resistance of the N-terminus of tau and if and how the native C-terminus of tau is a determinant of efficient fibril degradation.

### HTRA1 reduces the formation of tau aggregates in cells

To investigate how HTRA1 impacts tau aggregation in cells, a cell-based assay was established to quantify the seeding of intracellular tau aggregation and to study the effect of HTRA1. The assay is based on HEK293 cells stably overexpressing two tau variants consisting of its microtubule-binding domain tagged with CFP or YFP (biosensor cells) (20). Treatment of biosensor cells with tau seeds triggers the aggregation of the tau-CFP and tau-YFP, resulting in a fluorescence resonance energy transfer (FRET) signal that can be quantified by flow cytometry. The sensitivity of this assay was determined by titrating tau seeds to biosensor cells with or without lipotransfection (SI Appendix, Fig. S3A). The FRET signal increased in a dose-dependent manner and was up to >100-fold higher when tau seeds were lipotransfected compared to uptake without lipotransfection (20). The assay was highly sensitive with detection limits in the pM range, with respect to the tau seed concentration. To ensure that HTRA1 does not cleave the fluorescent tags off the hybrid proteins, and thus interfere with FRET flow cytometry analysis, the intactness of the majority of the tau hybrid proteins was demonstrated by Western blotting using extracts of cells overexpressing HTRA1 as well as the controls i.e. the empty vector and proteolytically inactive HTRA1^S328A^ (SI Appendix, Fig. S3B). To investigate the effect of HTRA1 on seeded tau aggregation, recombinant HTRA1 was applied in its proteolytically active or inactive forms, as the latter dissociates tau fibrils (10). When recombinant HTRA1 or HTRA1^S328A^ and subsequently tau seeds were added to the medium of biosensor cells, a dose-dependent lower FRET signal was detected for both HTRA1 forms, suggesting seed removal by proteolysis in the case of the active protease and dissociation in the case of HTRA1^S328A^ (Fig. 3A). Since the mean fluorescence intensity of YFP and CFP was not significantly different between the tested conditions, the HTRA1-mediated reduction of the FRET signal was not due to the degradation of the fluorescent tags themselves (SI Appendix, Fig. S3C). Both HTRA1 variants reduced the FRET signal by about 40%. The slightly greater effect of HTRA1^S328A^ might be best explained by its higher stability, as reflected by higher intracellular levels compared to active HTRA1 (Fig. 3B).

**Fig. 3.**
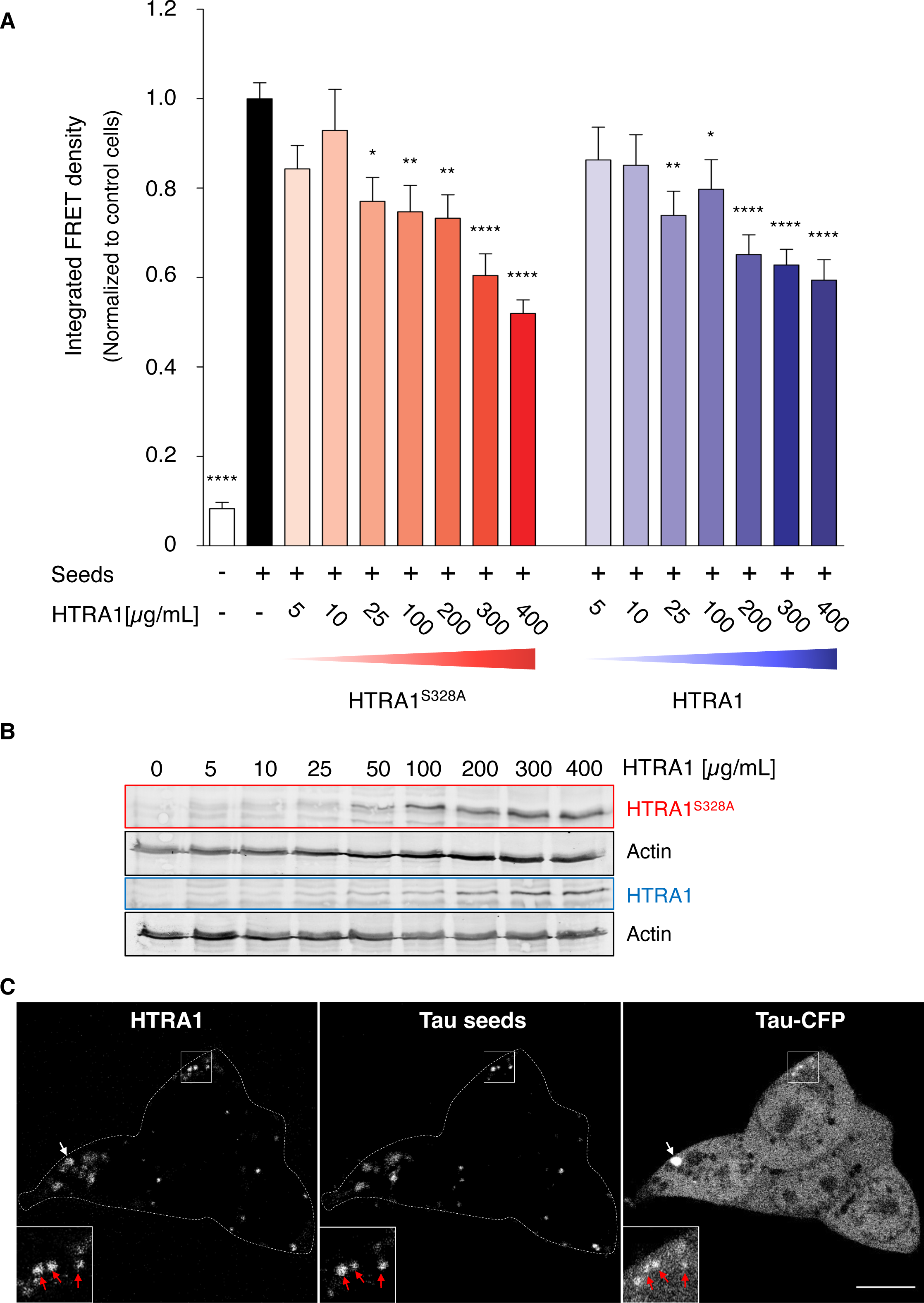
**HTRA1 reduces tau seeding in cells and co-localizes with aggregated tau upon internalization**. (A) HEK293 (biosensor) cells overexpressing tau-CFP and tau- YFP were treated with recombinant HTRA1, HTRA1^S328A^ or the corresponding buffer at the concentrations indicated. After 24 h, 100 nM recombinant tau seeds were added to the medium. After 20 h, tau aggregation was measured by flow cytometry determining the integrated FRET density (percentage of FRET positive cells multiplied by their mean fluorescence intensity). Biosensor cells treated with seeds only (black bar) were used for the normalization of values and the analysis of significant differences. Error bars, s.e.m.; n = 3. * P < 0.05, ** P < 0.01, **** P < 0.0001, one-way ANOVA with Dunnett’s posttest. (B) Biosensor cells were treated with recombinant HTRA1 and HTRA1^S328A^ at the concentrations indicated or the corresponding buffer control for 24 h. Whole cell extracts were subjected to SDS-PAGE and immunoblotted against HTRA1. Actin levels were used as loading controls. (C) Confocal microscopy analysis of biosensor cells treated with 25 µg/ml DyLight 633- labeled HTRA1^S328A^ followed by 300 nM Alexa Fluor 568-labeled seeds. Control images excluded interchannel bleed through (SI Appendix, Fig. S2). Scale bar: 10 µm. Arrows: Co-localization of HTRA1 with seeds and/or spots with a condensed CFP signal.

As a secreted protease, HTRA1 may interact with extracellular tau seeds. However, Western blotting showed the concentration-dependent internalization of HTRA1 and HTRA1^S328A^ upon treatment of biosensor cells (Fig. 3B), a phenomenon previously reported for other cell lines (10). Therefore, confocal microscopy was performed to localize internalized HTRA1 (Fig. 3C). Biosensor cells were treated with HTRA1^S328A^ and tau seeds labeled with DyLight 633 and Alexa Fluor 568, respectively. Control images verified the absence of inter-channel bleed through (SI Appendix, Fig. S3D). Internalized HTRA1^S328A^ was located in punctate agglomerates as previously reported (21) and colocalized with tau seeds. In addition, HTRA1^S328A^ and tau seeds were found at sites with a condensed CFP signal, indicating its localization to sites of seeded tau aggregation (Fig. 3C). To investigate whether HTRA1 targets preformed tau aggregates, biosensor cells were pre-treated with tau seeds, followed by the addition of HTRA1 or HTRA1^S328A^ to the medium of cells. Subsequent flow cytometry analysis revealed a dose-dependent reduction in FRET signal, indicating that both, proteolytically active and inactive HTRA1, interfered with intracellular tau aggregation in response to seeding (SI Appendix, Fig. S3E). The median fluorescence intensity of YFP and CFP was almost identical for the conditions tested (maximal reduction by 1% in the presence of HTRA1), suggesting that degradation of the fluorescent tags is not the cause of the decreased FRET signal (SI Appendix, Fig. S3F).

### HTRA1 reduces tau propagation

The pathogenic spreading of tau involves its aggregation in a donor cell and the concomitant externalization of transmittable tau species that promote the aggregation of endogenous tau in acceptor cells (22). As HTRA1 was shown to reduce tau aggregates and to counteract the seeded aggregation of tau in cells, we investigated its effect on the entire spreading process. We thus established a tau spreading assay using an indirect co-culture system consisting of two cell populations separated by a semi-permeable membrane that allows media exchange while preventing cell migration (Fig. 4A). Tau spreading was modeled by co-culturing donor cells harboring un-tagged tau aggregates with biosensor cells as acceptors. The underlying concept was that donor cells release a pathogenic tau conformation that enters acceptor cells to promote aggregate formation. The seeding of tau aggregates can be quantified by FRET flow cytometry. This setup also allows monitoring the effects of HTRA1 on the process of intercellular seeding. HEK293 cells, induced to co-express tau and HTRA1 or HTRA1^S328A^ were used as donor cells. Control donor cells co-expressed tau and GFP. To trigger tau aggregation, donor cells were incubated with recombinant tau seeds for 2 days. To exclude the possibility that extracellularly located recombinant seeds induce tau aggregation in acceptor cells, donor cells were treated with the protease trypsin prior to co-culturing. The efficient degradation of recombinant tau seeds by trypsin was demonstrated by an *in vitro* degradation assay mimicking cell culture conditions (SI Appendix, Fig. S4). Seeds were digested by trypsin within the first minute of incubation, while they were stable in PBS. After trypsinization, donor cells were replated and cocultured with acceptor cells. After 4 days of co-culture, tau aggregation in acceptor cells was quantified by FRET (Fig. 4B). Acceptor cells co-cultured with untreated donor cells exhibited a low FRET signal. The absence of protein overproduction in donor cells was verified by Western blotting (Fig. 4B). Induction of tau expression in donor cells without the addition of seeds caused an approximately 2-fold increase of the FRET signal in acceptor cells, indicating that tau overproduction causes a low level of aggregation (23), while treatment with seeds in the absence of tau overexpression increased the FRET signal by approximately 5-fold in acceptor cells. In addition, the combination of seed treatment and tau overexpression increased the FRET signal in acceptor cells by >10-fold. This suggests the formation of aggregates in donor cells and the release of seeding-competent tau generated from these aggregates. When donor cells expressed HTRA1^S328A^, the FRET signal in seeded acceptor cells overexpressing tau was reduced to approximately 50%. Co-expression of HTRA1^S328A^ did not alter the tau levels in donor cells as shown by Western blotting. HTRA1^S328A^ may have dissociated seeds and interacted with tau, thereby reducing its aggregation propensity. When donor cells expressed proteolytically active HTRA1, the FRET signal in acceptor cells was reduced to a similar level. Western blotting of cell extracts showed the degradation of tau in donor cells as well as auto-proteolysis of cytosolic HTRA1, which is a common phenomenon (24). Auto-proteolysis of internalized HTRA1 can be explained by the reducing environment of the cytosol causing instability of its N-terminal domain, which is stabilized extracellularly by eight disulfide bonds (25). These data suggest that tau aggregation in donor cells causes tau aggregation in acceptor cells and that HTRA1 reduces the efficiency of this process.

**Fig. 4.**
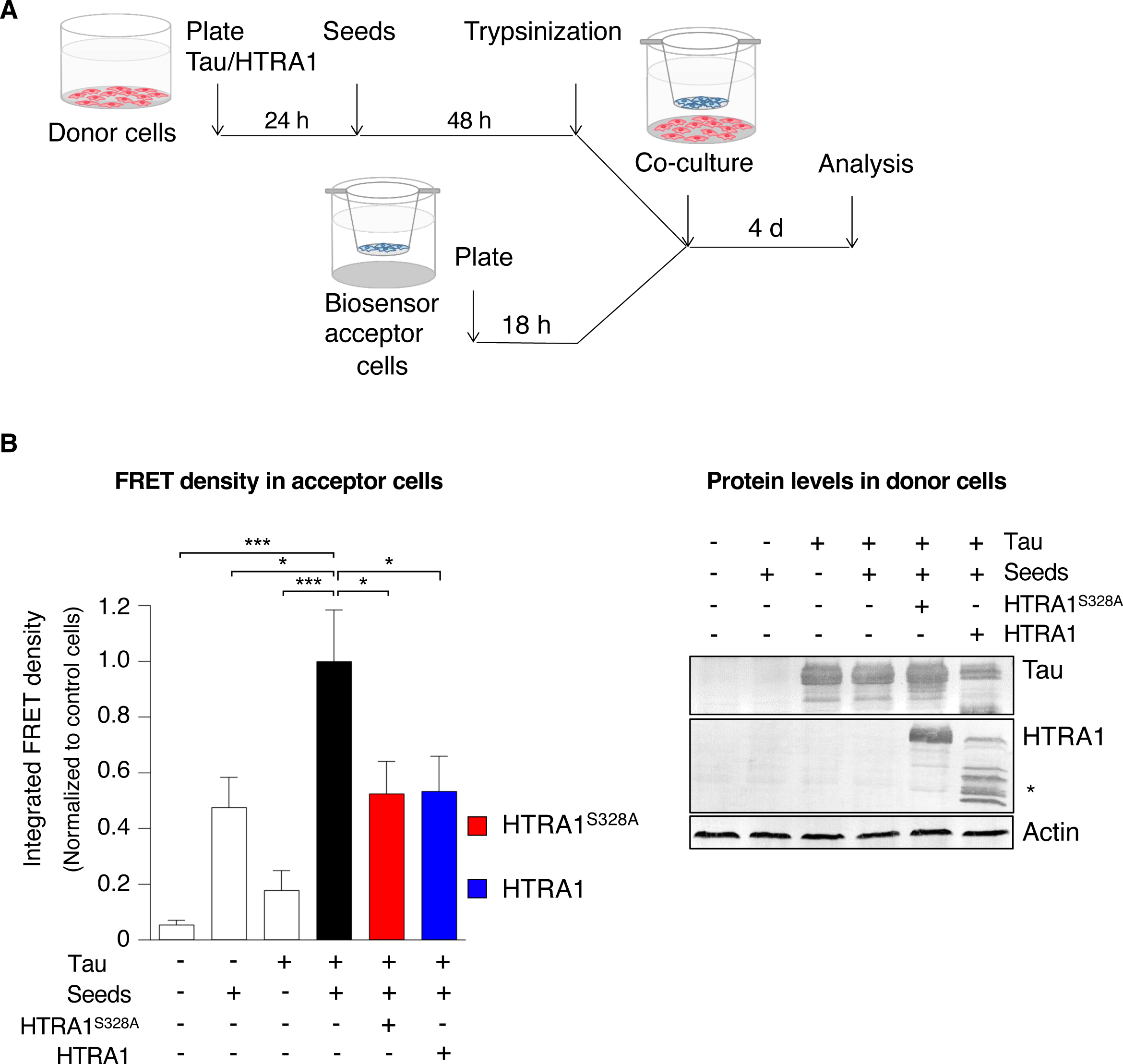
**Cellular model of tau spreading reveals inhibitory effect of HTRA1.** (A) Tau spreading was modeled by co-culturing donor cells harboring tau aggregates and untreated acceptor (biosensor) cells separated by a semipermeable membrane that allowed medium exchange, but not cell migration. Donor cells overexpressed tau and either myc-tagged HTRA1 or HTRA1^S328A^ respectively, or as a control, GFP after induction for 24 h. Intracellular aggregation of tau was induced by treating donor cells with 750 nM tau seeds. After 48 h, donor cells were trypsinized to remove extracellular seeds. Subsequently, biosensor acceptor cells were co-cultivated for 4 days (d). Tau spreading was assessed by FRET flow cytometry analysis of biosensor acceptor cells. (B) After co-culture as in (A), biosensor cells were analyzed by FRET flow cytometry as described in Fig. 3A. Integrated FRET density of cells overexpressing tau and treated with seeds (black bar) was used to normalize values and analyze significant differences. As a control, donor cells were untreated or treated with seeds only, and tau or GFP overexpression was induced in the absence of seeds. Error bars, s.e.m.; n = 3. * P < 0.05, *** P < 0.001, one-way ANOVA with Dunnett’s post-test. Donor cells were subjected to SDS-PAGE and immunoblotting using antibodies against tau, myc- tag of HTRA1 and actin, respectively. Bands marked with an asterisk represent autoproteolytic products of HTRA1.

### HTRA1 is transferred between cells

Since HTRA1 is a secreted protease that is internalized by various cell lines (10, 21), its cell-to-cell transmission was investigated by co-cultivation (Fig. 5A). Myc-tagged HTRA1^S328A^ was overexpressed in HEK293T donor cells and its transfer to acceptor cells was monitored using HEK293T or SH-SY5Y cells. Two types of acceptor cells were used to investigate potential cell type-specific differences in the transfer of HTRA1. Donor and acceptor cells were co-cultured for up to 4 days. Subsequently, whole cell extracts and media were subjected to Western blotting using an antibody against the myc-tag to exclude endogenously expressed HTRA1 from the analyses (Fig. 5B). HTRA1^S328A^ produced in donor cells appeared in the medium after 24 h and was internalized by SH-SY5Y acceptor cells after 24 h and by HEK293T acceptor cells after 72 h. Transferred HTRA1^S328A^ was detectable in both cell types for up to 4 days. To exclude that the HTRA1 detected in the acceptor cells was due to residual lipotransfection vesicles containing HTRA1 plasmids, HTRA1^S328A^ lacking its secretion signal was overexpressed in donor cells. The empty vector was used as an additional negative control. In both controls, acceptor cells did not contain myc-tagged HTRA1 (Fig. 5B). To quantify the efficiency of intercellular HTRA1 transfer, the number of acceptor cells that internalized HTRA1 was determined by confocal microscopy in co-culture assays using mCherry-tagged HTRA1 for detection. When co-cultured with HTRA1^S328A^-mCherry overexpressing donor cells, 93% of SH-SY5Y and 77% of HEK293T acceptor cells, respectively were mCherry positive (Fig. 5C). To demonstrate that mCherry itself is not transferred between cells, mCherry fused to the HTRA1 signal sequence was used as a control. Indeed, secreted mCherry did not generate fluorescent signals in acceptor cells. These data suggest that HTRA1 is efficiently transferred between cells. The fact that more SH-SY5Y cells internalized HTRA1^S328A^ is consistent with its faster uptake in this cell type as shown by Western blotting (Fig. 5B). In both acceptor cell lines, HTRA1^S328A^-mCherry was localized to punctate structures that were similarly seen in biosensor cells treated with recombinant HTRA1^S328A^ (Fig. 5D).

**Fig. 5.**
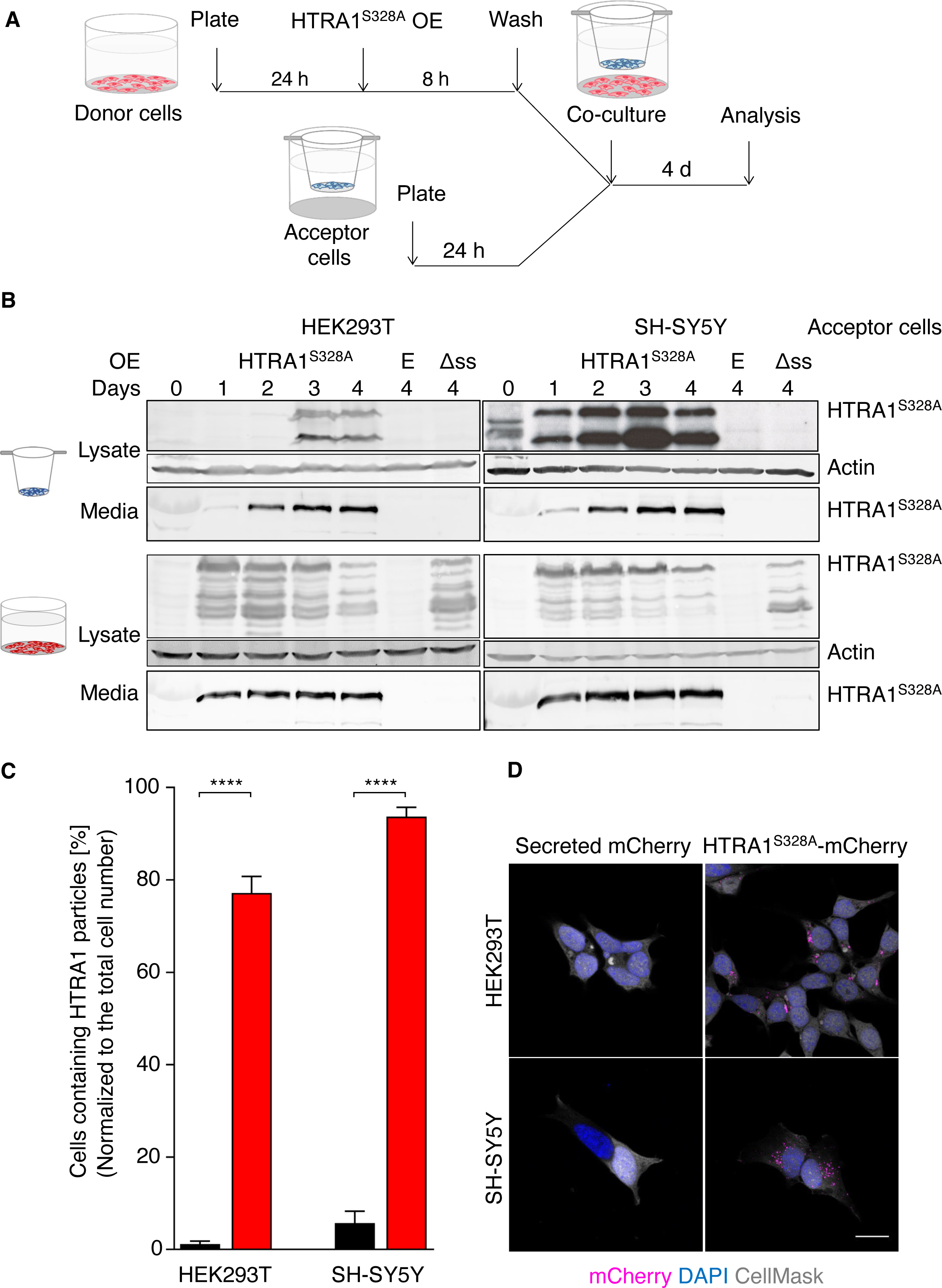
**Intercellular spreading of HTRA1.** (A) Intercellular transfer of HTRA1 was examined by co-culturing HEK293T donor cells transiently overexpressing myc-tagged HTRA1^S328A^ and untreated HEK293T or SH-SY5Y acceptor cells. (B) Media and lysates of donor and acceptor cells were subjected to SDS-PAGE and immunoblotting against the myc tag of HTRA1 and actin, respectively. As a control, donor cells were transfected with the empty vector (E) or myc-tagged HTRA1^S328A^ lacking the signal sequence for secretion (Δss). Note that extracellular HTRA1 (49 kDa) is partially or completely processed upon internalization into the cytosol into a smaller form. This ca. 35 kDa form does not contain the disulfide-bondend N-terminus for which no function has yet been identified. Note that the faint band in the Media samples at the 0 days timepoint is serum albumin present in the cell culture media. (C) Confocal microscopy analysis of the intercellular HTRA1 transfer. HEK293T or SH- SY5Y acceptor cells were co-cultured for 3 days with donor cells overexpressing either HTRA1^S328A^-mCherry or mCherry fused to the HTRA1 signal sequence. Secretion and fluorescence of both proteins are shown in SI Appendix, Fig. S2. Acceptor cells were fixed and cell nuclei and membranes were counterstained. Quantification of HTRA1 positive cells is based on 10 images taken per condition. Error bars, s.e.m.; n = 3. **** P < 0.0001, Mann-Whitney U test using acceptor cells co-cultured with donor cells overexpressing secreted mCherry as a control. (D) Representative images of HEK293T and SH-SY5Y acceptor cells from (C). DAPI (blue) and CellMask Deep Red dye (grey) were used to label nuclear DNA and the cell membrane, respectively. Scale bar: 10 µm.

Several additional control experiments were performed. Donor cells overexpressing mCherry fused to the HTRA1 signal sequence were analyzed by confocal microscopy. The vesicular distribution of the mCherry signal indicated its localization to secretory vesicles (SI Appendix, Fig. S5A). Fluorescence intensity measurements of the cell culture supernatant showed that secretion did not eliminate the fluorescence of the hybrid proteins (SI Appendix, Fig. S5B). In addition, Western blotting of the donor cell medium confirmed the secretion of mCherry and the presence of the HTRA1-mCherry hybrid protein (SI Appendix, Fig. S5C). As a previous study suggested a tag-dependent intracellular HTRA1 localization (21), HEK293T and SH-SY5Y cells were co-cultured with donor cells overexpressing myc-tagged HTRA1^S328A^. Immunostaining against the myc tag revealed no difference in the intracellular distribution of HTRA1^S328A^, demonstrating that the fluorescent tag did not affect the location of the transferred HTRA1 (SI Appendix, Fig. S5D).

### Intercellularly transferred HTRA1 colocalizes with tau seeds in recipient cells and reduces seeding of tau *in vitro*

Live cell spinning disk confocal microscopy was used to quantify the colocalization of intracellular HTRA1 during the uptake of tau seeds and the uptake of HTRA1 into cells already containing tau seeds over 30 h. SH-SY5Y cells were treated either with conditioned media containing secreted HTRA1^S328A^-mCherry followed by DyLight 488- labeled tau seeds or vice versa (termed HS and SH, respectively) (SI Appendix, videos S1 and S2). Morphological changes of the cells and cytotoxic effects were observed at longer imaging times, most likely due to starvation in media containing 1% FCS. The number of detectable discrete fluorescent spots was quantified over time in both, the mCherry and the DyLight 488 channels.

Efficient uptake was observed for extracellular tau seeds (HS treatment, Fig. 6A and C, Fig. S6A) and secreted HTRA1^S328A^-mCherry (SH treatment, Fig. 6B and 6D, SI Appendix, Fig. S6B). For both conditions, the number of discrete intracellular spots increased over time. Strikingly, a rapid co-localization was observed shortly after the internalization of the extracellularly applied proteins for both conditions, which slightly decreased over time, most likely due to the oversaturation compared to the population already present in the cytosol (SI Appendix, Fig. S6G, HTRA1-positive tau spots and Fig.S6H, tau-positive HTRA1 spots). For the subpopulation already present in the cytosol (HTRA1-positive Tau spots in SH treatment, tau-positive HTRA1 spots in HS treatment), the degree of co-localization increased steadily over time. Based on >10,500 HTRA1 and tau particles, respectively, we detected up to 60% colocalization in both conditions. For the subpopulation already present in the cytosol (HTRA1 in HS and Tau in SH treatment), we observed a decrease in the number of spots over time. A similar change was observed for the respective control treatments, where DMEM was applied instead of extracellular protein (SI Appendix, Fig. S6 C-F). Despite photobleaching effects, this is most likely due to the condensation of aggregates into larger complexes. In addition, the reduction in detectable tau seed spots in the control cells was less pronounced than in the cells treated with secreted HTRA1^S328A^-mCherry, suggesting that the presence of HTRA1^S328A^ reduces tau aggregation.

**Fig. 6.**
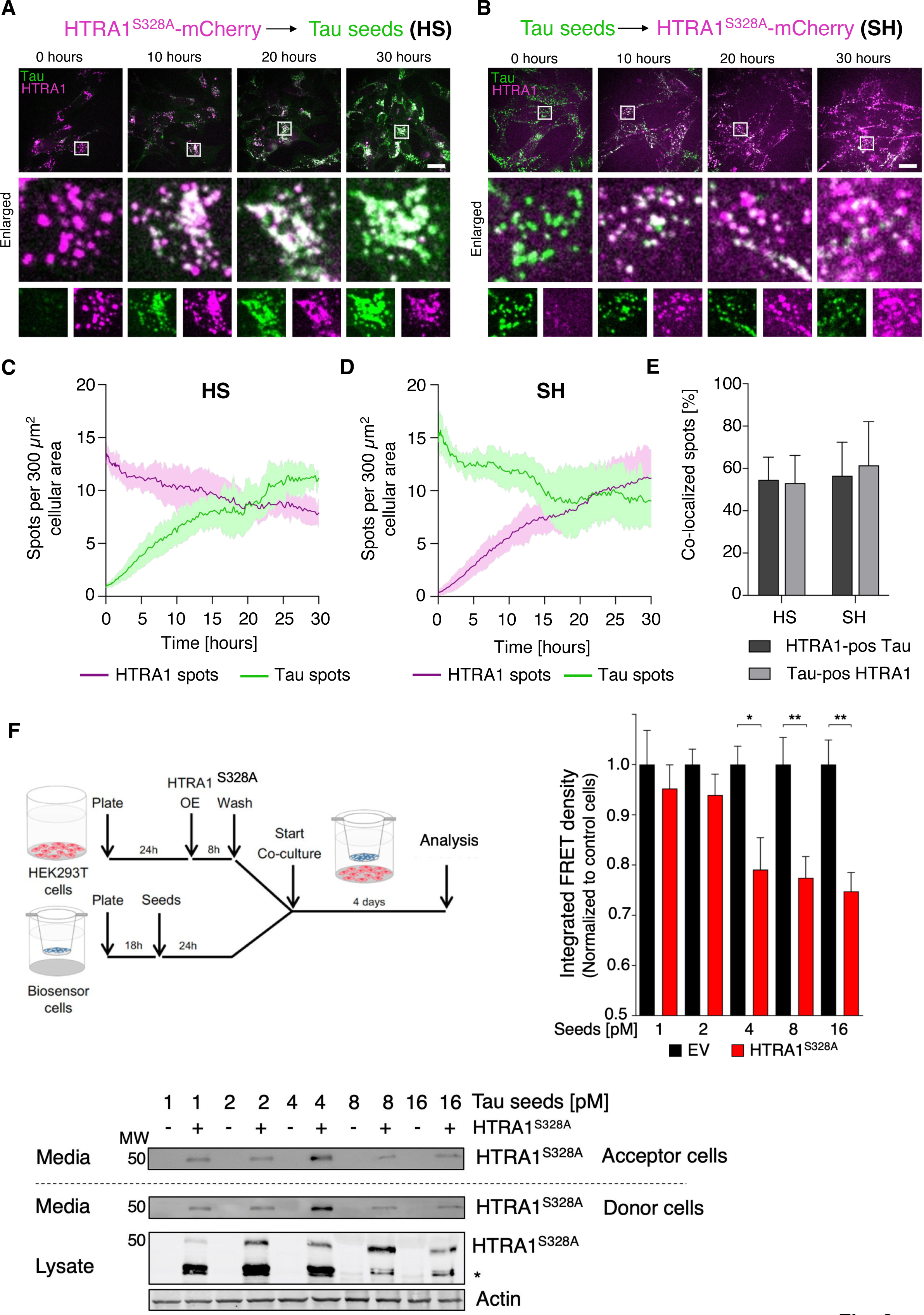
**Intercellularly transferred HTRA1 colocalizes with tau seeds in recipient cells.** (A) Live cell imaging of SH-SY5Y cells treated with conditioned medium containing secreted HTRA1S328A-mCherry (magenta) 24 hours before imaging. Cells were treated with DyLight 488-labeled tau seeds (green) immediately prior to imaging (HS treatment). Enlarged regions labeled in the top panel are shown as overlay (middle panel) and single channels (lower panel). (B) Live cell imaging of SH-SY5Y cells treated with DyLight 488-labeled tau seeds (green) 24 hours before imaging. Subsequently, cells were treated with conditioned medium containing secreted HTRA1S328A-mCherry (magenta) immediately prior to imaging (SH treatment). Enlarged regions marked in the upper panel are shown as overlay (middle panel) and single channels (lower panel). Scale bars: 20 µm. The intensity of the green (tau) channel is scaled differently for sub-figures A and B (A - HS: 0 – 350, B – SH: 0 – 140), to allow better visibility of individual spots. As the treatments were conducted 20 minutes before the start of imaging, there were already a very few spots visible at the 0-hour timepoint. (C-D) Time-resolved analysis of HTRA1^S328A^-mCherry and DyLight 488-labeled tau seed spots in SH-SY5Y cells (10 random positions per condition/experiment, n=3; in total, >10,500 individual HTRA1 and Tau spots were analyzed). Cells were either treated first with HTRA1^S328A^-mCherry followed by tau seeds (HS shown in C) or vice versa (SH shown in D). In each frame, the CellTracker Deep Red stain was used to detect the image area that was covered with cells (total cell area). Discrete fluorescent spots were detected in the mCherry and DyLight 488 channel in this region only. The number of detected discrete HTRA1^S328A^-mCherry (magenta line) and/or tau seed spots (green line) was normalized by the measured cell area (number of spots/µm^2^) and multiplied by the factor 300, which reflects the average cell area of an individual SH-SY5Y cell. Data show mean ± SD. Quantification of controls and example movies are shown in SI Appendix, Fig. S6 and Videos S1 and S2. (E) Co-localization analysis of HTRA1^S328A^-mCherry and DyLight 488-labeled tau seed spots at timepoint 20 hours after start of imaging. Discrete spots of HTRA1^S328A^- mCherry were detected and masked with DyLight 488-labeled tau seed spots to identify double-positive species (Tau-positive HTRA1 spots) or vice versa (HTRA1- positive Tau spots) (mean ± SD). Time-resolved analysis of co-localization data is shown in SI Appendix, Fig. S6. (F) Co-culture of donor cells overproducing HTRA1^S328A^ and biosensor cells containing tau aggregates. HEK293T donor cells were either transfected with HTRA1^S328A^ (OE) or the corresponding empty vector (EV). Prior to co-cultivation, biosensor acceptor cells were lipotransfected with between 1-16 pM of tau seeds. Co-cultivation was performed for 4 days, followed by the analysis of intracellular tau aggregation in biosensor cells by FRET flow cytometry as described in Fig. 3A. The EV control was used for the normalization of values and the analysis of significant differences. Error bars, s.e.m.; n = 2. * P < 0.05, ** P < 0.01, unpaired t-test. To show HTRA1^S328A^ levels, media of donor and acceptor cells and cell lysates of donor cells were subjected to SDS PAGE and immunoblotting using antibodies against its Myc-tag. For cell lysates, actin was used as a loading control. MW, molecular weight [kDa]. Note that upon internalization into the cytosol, extracellular 49 kDa form of HTRA1 is partially processed into a smaller form (35 kDa, marked by an asterisk) not containing the originally disulfide-bonded N-terminus. See text for details.

So far, our data suggest that cytosolic HTRA1 captures internalized tau aggregates during the seeding process. In addition, HTRA1 localizes to cytosolic tau seeds and tau aggregates after its own uptake. Thus, the intercellular transfer of HTRA1 may rescue acceptor cells from tau pathology. To test this hypothesis, the co-culture system was used (Fig. 6F). Acceptor cells were lipotransfected with tau seeds and then co-cultured with donor cells overexpressing HTRA1^S328A^. Western blotting confirmed the presence of HTRA1^S328A^ in donor cells and the medium (Fig. 6F). Co-culture of biosensor acceptor cells harboring seeded tau aggregates with HTRA1^S328A^- overexpressing donor cells resulted in a reduction of the FRET signal by up to 25% (Fig. 6F). Titration of tau seeds indicated that the protective effect of HTRA1^S328A^ is stronger at higher seed concentrations, probably because higher seed concentrations improve binding to HTRA1. These data support the hypothesis that the cell-to-cell transfer of HTRA1 counteracts tau propagation by reducing the seeded aggregation of tau and/or by clearing tau aggregates in acceptor cells.

## DISCUSSION

The secreted HTRA1 protease has the ability to diffuse and enter neighboring cells. The production and secretion of HTRA1 in a donor cell, and its uptake into neighboring acceptor cells with compromised proteostasis will decrease the levels of tau fibrils, thus reducing the ability to spread the pathogenic tau conformation. In addition, healthy acceptor cells taking up HTRA1 could be protected from tau aggregation and tau pathology. Along its way, HTRA1 dissociates and degrades tau seeds and amyloid fibrils. A more general protective role for HTRA1 is suggested by a recent report indicating that HTRA1 prevents the conversion of α-synuclein monomers to amyloid fibrils, dissociates α-synuclein amyloid fibrils, and interferes with seeding (26). Our data establish HTRA1 as a migrating protein quality control factor that interferes with several critical steps of the spreading process. This response to a particularly problematic cascade of protein misfolding and propagation events exemplifies the power of evolution, which in this case appears to be directed at delaying the onset of fatal neurodegenerative diseases. Interestingly, the production of HTRA1 by healthy e.g. non-neuronal cells that do not harbor tau fibrils, allows for non-autonomous proteostasis in neuronal cells. The concept of cell non-autonomous proteostasis was first established and is most widely studied in non-vertebrates but at least some aspects seem to be conserved in mammals (27, 28). Therefore, transcellular chaperone and protease signaling might represent an attractive direction of future research. In the case of HTRA1, transcellular signaling could modulate HTRA1 expression through activation of specific transcription factors or epigenetic mechanisms, as the HTRA1 promoter contains a CgG island (29).

As an ATP-independent enzyme, HTRA1 is uniquely suited to function intra- and extracellularly and has several additional advantages over other factors previously identified to dissociate or degrade fibrils. For example, while ATP-dependent heat shock chaperones and the AAA^+^ ATPase VCP are capable of dissociating fibrils from various sources, including patient brains, the extracted individual tau polypeptides remain seeding competent, presumably because they are not degraded by a protease (30–33). However, a collaboration between HTRA1 and these or other proteostasis factors with the sole ability to dissociate fibrils should not be excluded. One relevant example is the Calpain 2-HTRA1 complex, in which the binding of the cysteine protease Calpain 2 confers allosteric activation of HTRA1 (14).

Time-resolved mass spectrometry analysis demonstrates that the proteolytic degradation of amyloid fibrils by HTRA1 is initiated at regions surrounding the β-sheet core, followed by multiple cuts within the core region. This finding suggests a mechanism by which HTRA1 particularly targets aggregation-prone conformations, which can be explained by the induced fit mechanism of proteolysis. This model is supported by the stronger activation of the proteolytic activity by tau fibrils compared to unstructured soluble tau. In general, substrates bind to the active site of proteases in an extended β-strand conformation, which was also shown for HtrA proteases (34, 35). The linear extension of a substrate optimizes its interaction with the substrate binding pockets and optimally exposes the main chain amide atoms to the catalytic residue of the protease. However, the tight packing and extensive intramolecular hydrogen bonding of the sheet structures of the fibril core make it less favorable for interaction with the active site. Therefore, proteolysis by HTRA1 is preceded by fibril dissociation before extracted individual peptides can interact productively with the active site of the protease (10). Regarding the differences and similarities between the fibrillar and soluble substrates, we find a greater number of peptides and cleavage sites when fibrillar tau is proteolyzed. This finding can be explained by the optimized binding of preformed β-strands in peptides derived from fibrillar structures. However, the frequency of cuts is lower in fibrillar structures, which can be explained by the fact that individual peptides have to be extracted from the fibrils, which is expected to slow down proteolysis. In addition, the comparison of preferences for residues surrounding the scissile bond shows only minor differences between soluble and fibrillar tau, which makes sense because the substrate binding pockets of the active site are the same. In addition to HTRA1, numerous other proteases have been shown to cleave the tau protein, but the majority of the identified cleavage sites of 17 proteases were located outside of the β-sheet-forming regions, except for residue I308 that is targeted by the chymotrypsin-like activity of the 20S proteasome, and residue D314, which is targeted by caspase 2 (36). It is therefore not surprising that numerous tau fragments produced by a variety of proteases, targeting proteolytic sites located outside the microtubule-binding repeats and thus the β-sheet-forming regions have been shown to mediate oligomer formation, aggregation, and toxicity in cells or have been identified in patient brain samples and thus are suspected to be implicated in tauopathies (37, 38).

Our data highlight the therapeutic potential of HTRA1, not only because of its ability to track and degrade even moving targets, such as spreading tau seeds, but also because of its accumulation at sites of amyloid fibril deposition (13). Consistently, HTRA1 levels are elevated in AD patient samples and tau transgenic mouse models (12, 39–42). In addition, HTRA1 has previously been shown to degrade Aβ oligomers and ApoE4, the latter being a risk factor for late onset of AD (13, 43). As observed with tau, HTRA1 levels were elevated in Aβ-associated mouse models (44, 45). In addition, a large proteomics study of 610 brain tissues from individuals with clinical diagnoses of no cognitive impairment, mild cognitive impairment, and AD dementia detected elevated HTRA1 levels in ApoE4 carriers (46). Our data emphasize that the identification and characterization of evolutionarily conserved protein quality control factors such as HTRA1 warrants further efforts to translate the mechanistic insights obtained from basic research into treatment.

## MATERIALS and METHODS

### Antibodies and chemicals

For the detection of tau, a rabbit polyclonal antibody from Abcam (no. ab-64193) was applied. Rabbit polyclonal antibody against HTRA1 was published earlier (11). Protein tags were detected using mouse monoclonal antibodies against GFP, mCherry, and myc purchased from Hoffmann-La Roche (no. 11814460001, mixture of clone 7.1 and 13.1) and Cell Signaling (no. 213511, clone EPR20579; no. 2276S, clone 9B11). Actin was detected by a mouse monoclonal antibody from MP Biomedicals (no. 691001, clone 4) or a rabbit monoclonal antibody from Abcam (no. ab-198991, clone EPR16770). Alkaline phosphatase labeled polyclonal goat antibodies against mouse and rabbit were purchased from Sigma Aldrich and DaKoCytomation, respectively. For immunofluorescence, polyclonal goat anti-mouse and rabbit antibodies labeled with Alexa Fluor 488 as well as DAPI and CellMask Deep Red Plasma membrane stain were purchased from Thermo Fisher Scientific. Amine-reactive protein labeling dyes DyLight 488 NHS Ester, Alexa Fluor 568 NHS Ester and DyLight 633 NHS Ester were obtained from Thermo Fisher Scientific. Heparin and ThT were from Sigma Aldrich and AAT Bioquest, respectively.

### Plasmids

Bacterial expression plasmids for the purification of 4R2N tau as well as Strep-tagged HTRA1 and HTRA1^S328A^ lacking the mac domain were published previously (10). Plasmids based on the pcDNA5/FRT/TO vector backbone (Thermo Fisher Scientific) were used for the generation of Flp-In T-REx cell lines and the transient overexpression in mammalian cells. Plasmids were produced by PCR and standard cloning techniques. The pcDNA5/FRT/TO plasmids containing the sequence of *4R2N tau^P301L^*, *HTRA1*, one or two *HTRA1^S328A^* cistrons or *HTRA1^S328A^* lacking the secretion signal sequence were generated from prior constructs (10). Bi-cistronic constructs of *HTRA1* and *tau* were obtained by cloning the genes into a previously published plasmid (48). Multi-cistronic vectors harbored a 2A peptide sequence to ensure protein co-expression. Plasmids containing a mCherry or GFP sequence were generated using published constructs (48, 49).

### Purification of recombinant HTRA1 and tau

Recombinant 4R2N tau (441-residue isoform) as well as Strep-tagged HTRA1 and HTRA1^S328A^ (residues 158–480) were purified as described (10).

### Heparin induced fibril formation of recombinant tau, generation of tau seeds and effects of HTRA1

Fibril formation and generation of seeds were done as described (10) To study the effects of HTRA1, HTRA1^S328A^ was pre-incubated with seeds for 3 h, while HTRA1 was directly added to tau and seeds at concentrations indicated. As a control, tau, seeds, HTRA1 and HTRA1^S328A^ were incubated alone. Fibril formation was analyzed by sedimentation assays as described (10). Protein levels of tau were determined by densitometry analysis using Image J (50). For transmission electron microscopy of fibrils, samples were diluted, subjected to a formvar-coated copper grid and incubated for 60 sec. After removal of excess liquid, the grids were incubated with a staining solution of 0.75% Uranyl formate, 6 mM NaOH and dried at room temperature. The samples were analyzed with a JEOL JEM 1400 PLUS at 120 kV. At least 25 images (7.89 µm x 7.89 µm) were taken per condition. Quantification of the total fibril length per image was performed using the ridge detection plugin of Image J (51). The kinetics of fibril formation were investigated by measuring ThT fluorescence after 4, 24, 48 and 72 h.

### HTRA1 activity assay

The specific activity of recombinant HTRA1 in presence of Tau species was measured using a synthetic HTRA1 substrate consisting of para-nitroanilin (pNA) coupled to the C-terminus of the peptide VFNTLPMMGKASPV (16). HTRA1 (1 µM) was mixed with various concentrations of Tau species to a final volume of 100 µl in 100 mM HEPES 100 mM NaCl pH 7.5. After incubation for 1 min at 37 °C and 700 rpm, 500 µM of the pNA substrate was added. Cleavage of the substrate was continuously monitored at a wavelength of 405 nm for 2 h using the SpectraMax iD5 spectrophotometer. The specific enzyme activities of HTRA1 were derived from at least three independent measurements and calculated with the following formula:

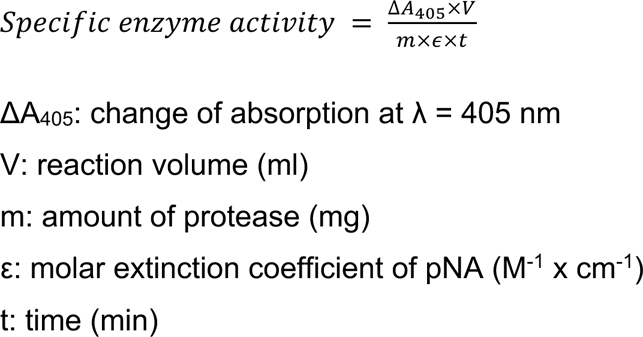

### Crosslinking-mass spectrometry

5 µM tau fibril seeds and 5 µM HTRA1S328A were preincubated in 100 mM HEPES, 100 mM NaCl pH 7.5 for 15 min at 37°C to allow binding. Subsequently, 2 mM (final concentration) of the cross-linking agent PhoX was added. After 15 min, the reaction was stopped by quenching with Tris-HCl. Samples were subjected to trypsin digests and prepared for mass spectrometry. For peptide identification, the RAW-files were loaded into Proteome Discoverer (version 2.5.0.400, Thermo Scientific). All created MS/MS spectra were searched using MSAmanda v2.0.0.16129 (52). Initially, the RAW-files were searched against the databases ID1416.fasta (2 sequences; 782 residues), tags_v11.fasta (28 sequences; 2 153 residues), uniprot_reference_E-coli_k12_2023-09-19.fasta (4 362 sequences; 1 354 446 residues) and PD_Contaminants_TAGs_v20_tagsremoved.fasta, using following search parameters: The peptide mass tolerance was set to ±10 ppm and the fragment mass tolerance to ±10ppm. The maximal number of missed cleavages was set to 2. The result was filtered to 1% FDR on protein level using Percolator algorithm integrated in Thermo Proteome Discoverer. For the 2nd step, the RAW-files were searched against the created sub-databases termed ID1416.fasta (Tau and HTRA1 sequences; 782 residues), using the following search parameters: iodoacetamide derivative on cysteine was set as a fixed modification, oxidation on methionine, phosphorylation on serine, threonine and tyrosine, deamidation on asparagine and glutamine, PhoX on lysine, pyro-glu from q on peptide N-terminal glutamine, acetylation on protein N-Terminus were set as variable modifications.

Monoisotopic masses were searched within unrestricted protein masses for tryptic enzymatic specificity. The peptide mass tolerance was set to ±10 ppm and the fragment mass tolerance to ±10 ppm. The maximal number of missed cleavages was set to 2. The result was filtered to 1% FDR on protein level using Percolator algorithm integrated in Thermo Proteome Discoverer. Additional high quality filtering by setting a minimum MS Amanda Score of 150 on PSM level was applied. Protein areas have been quantified using IMP-apQuant (53) by summing unique and razor peptides and applying intensity-based absolute quantification (54) with subsequent normalization based on the MaxLFQ algorithm (55). Proteins were filtered to be identified by a minimum of 2 PSMs in at least 1 sample.

### Time-resolved proteolysis and analysis by mass spectrometry

For proteolysis, 10 µM of tau soluble or fibrillar tau protein was mixed with 2 µM of HTRA1 in 100 mM HEPES 100 mM NaCl pH 7.5. and incubated at 37 °C and 350 rpm. Samples (20 µl) were taken at the indicated timepoints and directly added to 120 µl ice-cold acetone for overnight precipitation at -80°C. Precipitated proteins were removed by centrifugation (15,000 rpm, 1 h, 4 °C) and the peptide-containing supernatant was lyophilized in a SpeedVac concentrator at 30 °C. Dried samples were stored at -80 °C.

### LC/MS/MS

Experiments were performed on an Orbitrap Elite or Fusion Lumos mass spectrometer (Thermo Fischer Scientific, Waltham, Massachusetts, USA) that was coupled to an Evosep One liquid chromatography (LC) system (Evosep Biosystems, Odense, Denmark). Analysis on the Evosep One was performed on a commercially available EV-1064 Analytical Column – 60 & 100 samples/day (Length (LC) 8 cm; ID 100 µm; OD 360 mm; emitter EV-1086 Stainless steel emitter). The LC system was equipped with two mobile phases: solvent A (0.1% formic acid, FA, in water) and solvent B (0.1% FA in acetonitrile, ACN). All solvents were of UHPLC (ultra-high-performance liquid chromatography) grade (Honeywell, Seelze, Germany). For analysis with the Evosep One, samples were first loaded onto Evotips by following the manufacturer’s guidelines. For peptide separation, we used the 60 samples per day gradient which has an effective gradient of 21 min.

The mass spectrometers were operated using Xcalibur software (Elite: v2.2 SP1.48 or Lumos: v4.5.445.18). The mass spectrometers were set in the positive ion mode. Precursor ion scanning (MS1) was performed in the Orbitrap analyzer (FTMS; Fourier Transform Mass Spectrometry with the internal lock mass option turned on (lock mass was 445.120025 m/z, polysiloxane) (56). MS2 Product ion spectra were recorded only from ions with a charge greater +1 and in a data dependent fashion in the Ion Trap Mass Spectrometry. All relevant MS settings (Resolution, scan range, AGC, ion acquisition time, charge states isolation window, fragmentation type and details, cycle time, number of scans performed, and various other settings) for the individual experiments can be found in Supplemental data 5.

### Peptide and Protein identification using MaxQuant

RAW spectra were submitted to an Andromeda (57) search in MaxQuant (version 2.0.2.0 or 2.0.3.0) using the default settings (58). Label-free quantification (LFQ)(55) and match between runs was activated. Normalization in MaxQuant was switched off. MS/MS spectra data for Project ACE_0711 were searched against the ACE_0699_UP000000625_83333.fasta (4450 entries) custom database. For project ACE_0809 MS/MS spectra data were searched against the ACE_0809_SOI_v01.fasta (5 entries) custom database (contains sequences of interest) and the contaminants database as implemented in MaxQuant. Andromeda searches allowed oxidation of methionine residues (16 Da) and acetylation of the protein N-terminus (42 Da). No static modifications were set. Enzyme specificity was set to “unspecific”. The instrument type in Andromeda searches was set to Orbitrap and the precursor mass tolerance was set to ±20 ppm (first search) and ±4.5 ppm (main search). The MS/MS match tolerance was set to ±20 ppm. The peptide spectrum match FDR and the protein FDR were set to 0.01 (based on target-decoy approach). Minimum peptide length was 7 amino acids. For protein quantification, unique and razor peptides were allowed. In addition to unmodified peptides, modified peptides with dynamic modifications were allowed for quantification. The minimum score for modified peptides was set to 40.

### Data availability

The mass spectrometry data have been deposited to the ProteomeXchange Consortium via the PRIDE(59) partner repository (https://www.ebi.ac.uk/pride/archive/) with the dataset identifier PXD052323.

### UMSAP

MS-data were evaluated with the Targeted Proteolysis module of UMSAP 2.2.1 (19). MS-data of the tau alone samples served as reference for all UMSAP calculations. The significance level was set to 0.05 and the minimum score value to 50. A log2 transformation was applied to the data before the analysis. The amino acid distribution around the cleavage sites included 5 residues in each direction. The amino acid sequence used for this calculation are shown in SI Appendix, Supplemental data 1.

### Calculation of the relative frequency of cuts by UMSAP

The calculation of the frequency of cuts is performed in 2 steps. First, UMSAP groups all MS-detected peptides that share the same P1-P1’ bond. Subsequently, the average intensities for each peptide in each experiment is calculated. Subsequently, for each peptide the average intensity ratios are calculated taking the first average intensity >0 along the timepoints for each peptide as reference. Subsequently, the relative frequency of cuts for a P1 site at a specific timepoint is computed by summing the average intensity ratios of all peptides sharing the same P1 site.

If the peptide was not detected at a timepoint or the intensity values are not significantly different to the control experiments the average intensity is set to zero for this timepoint. A numeric example is provided in Supplemental data 6.

### Cell lines and culture media

SH-SY5Y and HEK293T cells were purchased from American Type Culture Collection (CRL-3216, CRL-2266). Biosensor cells were published previously (20). HEK293 based cell lines were cultured in Dulbecco’s Modified Eagle Medium (DMEM) and SH-SY5Y cells in a 1:1 mixture of DMEM and Ham’s F-12 Medium. Both media were supplemented with 10% fetal calf serum and 1% penicillin/streptomycin unless described otherwise. Co-culture of HEK293T and SH-SY5Y cells was performed with DMEM mixed with Ham’s F-12 Medium. For live cell imaging experiments, medium lacking phenol red was applied. The generation of Flp-In T-REx expression cells was performed as described by Thermo Fisher Scientific. For experiments performed with Flp-In T-Rex cells tet system approved serum (Takara Bio) was used.

### Protein labeling

Recombinant HTRA1^S328A^ was labeled with the amine reactive DyLight 633 NHS Ester, while fibrils made of recombinant 4R2N tau were labeled with DyLight 488 NHS Ester (used for the treatment of SH-SY5Y cells) or Alexa Fluor 568 NHS Ester (used for the treatment of biosensor cells), respectively. The labeling reaction was performed as described (10).

### Biosensor assay

To quantify the intracellular aggregation of tau, a FRET-based flow cytometry approach was applied (20). Biosensor cells were used that stably overexpressed *MTBD tau^P301S^* tagged with CFP or YFP. The sensitivity of the assay was examined by plating 3.5 x 10^4^ biosensor cells onto 96-wells using 130 µl medium. Cells were grown for 24 h and treated with the indicated amounts of recombinant tau seeds. Seeds were either added directly in a volume of 20 µl or by transfection with Lipofectamine 2000 as specified (20). After 18 h, FRET flow cytometry analysis was performed as described (20) to determine the integrated FRET density, which is the product of the percentage of FRET positive cells and their median fluorescence intensity.

### **Analysis of** *in cellulo* **tau aggregation in the presence of recombinant HTRA1**

The effects of recombinant HTRA1 on the seeding of tau aggregation and on pre- formed tau aggregates in cells were analyzed using the biosensor assay described above. For analyzing seeded tau aggregation, 3.5 x 10^4^ biosensor cells were plated onto poly-L-lysine coated 96-wells. On the following day, cells were washed and treated with 95 µl serum-free medium containing recombinant HTRA1 or HTRA1^S328A^ in the indicated concentrations. As a control, the medium was supplemented with the corresponding buffer. After 24 h, 100 nM recombinant tau seeds or Dulbecco’s phosphate-buffered saline (DPBS) were added in a volume of 5 µl to the medium of cells. Analysis was performed after 18 h by FRET flow cytometry as described above. The effect of HTRA1 on pre-formed aggregates was analyzed by adding 5 µl recombinant tau seeds (final concentration: 25 nM) or DPBS to the medium of cells. After 24 h, cells were washed and 100 µl serum-free medium containing recombinant HTRA1 or HTRA1^S328A^ in the concentrations indicated or the corresponding buffer were added to cells. After 18 h, FRET flow cytometry analysis was performed.

### Internalization of recombinant HTRA1 by biosensor cells

5 × 10^5^ biosensor cells were seeded onto poly-L-lysine–coated 6-wells. Cells were grown for 24 h, washed and treated with 1 ml serum-free medium containing recombinant HTRA1 or HTRA1^S328A^ in the indicated concentrations. On the following day, cells were detached by trypsin-EDTA treatment, centrifuged and cell pellets were washed with PBS. Pellets were lysed with RIPA buffer containing protease inhibitor (Roche) and analyzed by SDS-PAGE and immunoblotting. For microscopy analysis, 3.5 x 10^4^ biosensor cells were plated onto poly-L-lysine–coated µ-slide 8-wells (Ibidi). Cells were grown for 24 h, washed and subsequently treated with 200 µl serum-free medium containing 25 µg/ml recombinant DyLight 633 labeled HTRA1^S328A^ or the corresponding buffer. On the following day, cells were incubated with 300 nM Alexa Fluor 568 labeled tau seeds or DPBS for 8 h prior to live cell imaging. Images were acquired using a Leica TCS SP8X Falcon confocal laser scanning microscope equipped with a HC PL APO 63x/1.4 oil-immersion objective. An argon laser was applied at 458 nm (CFP) and a WLL E laser at 568 and 633 nm (Alexa Fluor 568 and DyLight 633) for the excitation of samples. The LAS X Core software (Leica Microsystems) was used for image acquisition and hardware controlling. For samples and control conditions, the same detector sensitivity settings were used. Images of different channels were taken using the serial acquisition mode to avoid inter-channel bleed-through.

### Tau spreading model

Intercellular tau spreading was analyzed by an indirect co-culture system comprising a well containing donor cells harboring tau aggregates and an inlaid insert containing biosensor cells as acceptors. Well and insert were separated by a semi-permeable membrane (Corning). For the generation of donor cells, Flp-In T-REx expression cells were used overexpressing *4R2N tau^P301L^* and simultaneously full-length *HTRA1* or *HTRA1^S328A^* upon doxycycline induction. As a control, *tau* was co-expressed with *GFP*. To trigger tau aggregation, 2.5 x 10^5^ donor cells were plated onto poly-L-lysine coated 24-wells and treated with 100 ng/ml doxycycline for 24 h followed by the addition of 750 nM recombinant tau seeds in 250 µl serum-free medium. On the following day, cells were supplied with 10% serum. After 2 days of incubation with tau seeds, donor cells were washed and treated with 150 µl trypsin-EDTA for 3 minutes to remove extracellular seeds. Trypsin was inactivated by adding 450 µl medium containing 1% serum. Subsequently, donor cells were plated onto poly-L-lysine coated co-culture 24-wells using 600 µl medium and an identical cell number for all conditions. After 6 h of cell seeding, co-culture with biosensor cells was started that were plated 18 h earlier using 100 µl medium and 3.5 x 10^4^ cells per insert. Co-culture was performed in medium containing 1% serum and 100 ng/ml doxycycline for 4 days. After 2 days of co-culture, doxycycline was re-added to the medium. As a control, biosensor cells were co-cultured with donor cells that were untreated or treated only with tau seeds or doxycycline. Biosensor acceptor cells were analyzed by FRET flow cytometry, while donor cells were trypsinized, lysed in RIPA buffer containing protease inhibitor and subjected to SDS-PAGE and immunoblotting.

### Digest of recombinant tau seeds by trypsin-EDTA

To verify the efficient clearance of recombinant tau seeds by trypsin-EDTA treatment performed in the tau spreading assay, 750 nM recombinant tau seeds were incubated in trypsin-EDTA. As a control, seeds were added to DPBS and trypsin-EDTA was incubated alone. Samples were taken at the time points indicated and analyzed by SDS-PAGE and Coomassie staining to detect trypsin and immunoblotting to detect tau.

### Live cell imaging of SH-SY5Y cells treated with recombinant tau and HTRA1

For analyzing the localization of internalized HTRA1 in cells containing tau seeds, 8 x 10^4^ SH-SY5Y cells, which were pretreated with “deep red cell tracker” (Thermofisher, C34565), were plated onto µ-slide 8-well plates. On the following day, 300 nM DyLight 488 labeled tau seeds or DPBS were added to cells. After 24 h, cells were washed and treated with 300 µl conditioned medium containing 1% serum and HTRA1^S328A^- mCherry or secreted mCherry, respectively, followed by live cell microscopy. Tau seeding in the presence of HTRA1 was investigated by treating cells with 300 µl conditioned medium containing 1% serum and HTRA1^S328A^-mCherry or secreted mCherry. After 24 h, fresh DMEM imaging medium with 1% FCS and 300 nM DyLight 488 labeled tau seeds or DPBS were added to cells.

Live-cell timelapse confocal spinning disk microscopy was performed using an inverted Nikon Eclipse TiE microscope system (Nikon Europe B.V.) equipped with an Andor REVOLUTION 500 AOTF Laser Combiner, a CSU-X1 Yokogawa spinning disk unit, and an iXon3 897 single photon detection EMCCD camera (Andor Technology). Laser lines used for excitation were diode laser cw 640 nm for CellTracker Deep Red, 561 nm for mCherry, and 488 nm for DyLight 488. The images were acquired using a CFI Plan Apo 40×/0.95NA dry objective lens (Nikon) and emission filters from Semrock. Specifically, the single-band pass filters used were Cy5 (BP 685/40 nm) and RFP (617/73 nm), and the dual-band pass filter used was EGFP/Cy5 (BP 538/50 nm + BP 685/45 nm). Acquisition was controlled by Andor IQ software (Andor Technology). Images were acquired at 10 random positions per condition with a time interval of 5 minutes for the first 2 hours and 15 minutes for the following 46 hours. Nikons hardware-based autofocus system PFS (Nikon Europe B.V.) was used to prevent drift and keep focus position stable. Live-cell imaging was performed at 37°C and 5% CO2 in imaging medium.

### Image Analysis

Automated image analysis was performed with CellProfiler (version 4.2.6) (60). Briefly, cells were detected using the Run Cellpose plugin for CellProfiler using the cyto2 model (expected object size 100, cell probability threshold -3 and flow threshold 0.8) with Omnipose for mask reconstruction(61, 62) on the rescaled (full intensity range) and smoothed (median filter artifact diameter 2) CellMask Deep Red image. The morphological challenges of SH-SY5Y cells, such as their partially overlapping spindle shape, can make it difficult to reliably define individual cells and cell boundaries. Therefore, we used the entire area covered by cells instead of quantifying on a per-cell basis. Thus, we identified regions covered by cells using the CellTracker Deep Red whole cell stain and combined them to form a single entity (total cell area) and detected discrete fluorescent spots within this region. To account for variations in cellular coverage over time within the image area, we normalized the number of spots by the measured cell area (number of spots/µm^2^) and multiplied this factor by 300, which reflects the average cell area of an individual SH-SY5Y cell. To detect discrete fluorescent Tau seed spots that co-localize with HTRA1S328A-mCherry, the segmented Tau spots were masked for HTRA1 detected spots. Only spots with a minimal overlap of 40% were considered double-positive (HTRA1-positive Tau spots). The same criteria were applied for the opposite combination (Tau-positive HTRA1 spots). To enhance vesicular structures of both, Tau seeds and HTRA1 spots, difference of Gaussians (DoG) was applied to the respective channels (gaussian filters of size 2 and 8). Discrete spots of Tau and/or HTRA1 signal were detected and related to the respective total cell area. The MaskObjects module was used to detect colocalized spots if their overlap was at least 40%. Data extraction, analysis and graph plotting were conducted using Microsoft Office Excel and GraphPad Prism 10.

### Analysis of the tag-dependence of internalized HTRA1

To exclude an impact of the mCherry-tag on the distribution of internalized HTRA1, co-culture was performed as described above with the exception that donor cells were transfected with plasmids expressing *myc-tagged HTRA1^S328A^* or with the corresponding empty vector. Microscopy analysis of SH-SY5Y and HEK293T acceptor cells was performed after 3 days of co-culture by staining cells using an antibody against the myc-tag, DAPI and CellMask Deep Red Plasma membrane stain. Images were acquired at the Leica TCS SP8 confocal laser scanning microscope as described earlier, however, the Alexa Fluor 488 labeled secondary antibody used for the detection of the my-tag antibody was excited with an argon laser.

### The intercellular transfer of HTRA1 in tau pathology

The effects of transferred HTRA1 on cells harboring tau aggregates were examined in the co-culture system. Either biosensor or SH-SY5Y cells were used as acceptors. For the biosensor setup, donor cell population was prepared by plating 2.5 x 10^5^ HEK293T cells onto poly-L-lysine coated co-culture 24-wells. After 24 h, donor cells were lipotransfected with HTRA1^S328A^ or the corresponding empty vector, followed by co-cultivation with acceptor cells. The acceptor cell population was prepared by plating 4 x 10^4^ biosensor cells per insert. On the following day, cells were lipotransfected with recombinant tau seeds in the indicated concentrations or DPBS. After 24 h, acceptor cells were subjected to co-culture for 4 days. Co-culture was performed in serum-free media added in a volume of 600 µl to wells and 100 µl to inserts. Biosensor cells were analyzed by FRET flow cytometry, while donor cells were trypsinized, lysed in RIPA buffer containing protease inhibitor and subjected to immunoblotting. The medium of wells and inserts was centrifuged at 1,000 g, 4°C for 5 minutes. The resulting supernatant was analyzed by immunoblotting.

The effect of transferred HTRA1 was further tested on pre-formed aggregates and tau seeding in SH-SY5Y acceptor cells. HEK293T donor cells were plated onto poly-L-lysine coated co-culture 6-wells. Prior to co-culture, donor cells were transfected either with a plasmid containing one or two *HTRA1^S328A^* cistrons or with the corresponding empty vector. Pre-formed tau aggregates were analyzed by plating 5x10^5^ SH-SY5Y cells onto co-culture inserts. After 24 h, cells were washed and treated with medium containing 1% serum and 750 nM recombinant tau seeds. On the following day, SH- SY5Y cells were washed before starting co-culture. For analyzing tau seeding, SH- SY5Y cells were plated as described above, however, treatment with 500 nM recombinant tau seeds was performed at the beginning of co-culture. Donor and acceptor cells were co-cultured for 4 days using medium supplemented with 1% serum added in a volume of 2.6 ml to wells and 1.5 ml to inserts. Analysis of SH-SY5Y acceptor cells was performed by sarkosyl extraction and sedimentation analysis as described (10). The supernatant and pellet fractions were analyzed by immunoblotting. Donor cells were trypsinized, lysed with RIPA buffer containing protease inhibitor and washed with PBS. Medium was harvested, centrifuged at 1,000 g, 4°C for 5 minutes. The supernatants and cell lysates were subjected to immunoblotting.

## Supporting information

Movie S1, Cyto tau fibrils (green), extracell HTRA1 (red)

Movie S2, Cyto HTRA1 (red), extracell tau fibrils (green)

## ACKNOWLEDGEMENTS

This work was funded by grant EH 100/19-1 (to M.E.) and SFB1430 - Project-ID 424228829 (to M.E., F.K., M.K., D.H. and S.P.) from Deutsche Forschungsgemeinschaft (DFG, German Research Foundation).

## AUTHOR CONTRIBUTIONS

A.B, B.H., K.R., N.S., M.S., M.C. carried out experiments and analyzed data; M.M., S.P., D.H., S.B. analyzed data; F.K, M.K. performed mass spectrometry; M.Er., S.S. performed TEM; A.B., S.P., D.H., S.B., M.Eh. outlined the work and wrote the paper with contributions of all authors.

