## Supplemental information for "A non-autonomous protein quality control mechanism targeting tau aggregate propagation"

Supporting text

Figures S1 to S6

Tables S1 to S3

Legends for Movies S1 to S2 SI References

Datasets S1 to S6

**Supporting Information Text**

**High-resolution analysis of proteolytic processing of soluble tau and tau fibrils**

Each proteolytic product is the result of two cuts, one at its N-terminus, where the N-terminal residue represents the P1' residue, and another at its C-terminus, representing the P1 residue (Fig. 2C). UMSAP also calculates the relative frequency of cuts at each P1 residue for each time point (see Methods for details). These data provide quantitative information on how well HTRA1 cleaves the substrate at each position within its primary amino acid sequence and how proteolysis progresses over time (SI Appendix, Fig. S2A). In addition, individual P1 residues can be grouped into several classes, based on histograms (SI Appendix, Fig. S2E-G). The first class consisted of 7 P1 residues (V256, C291, I308, C322, L376, I392, L408) where the relative frequency of cuts reached ≥20, i.e. 22-51 (SI Appendix, Fig. S2, Table S1). The second class consisted of 10 P1 residues (L266, I277, I278, V287, S289, I328, L344, V411, L428, V432) where the relative frequency of cuts ranged from 10 to 19 (SI Appendix, Fig. S2, Table S2). The third class consisted of residues where the relative frequency of cuts ranged from 1 to 10. These were considered poor sites at all time points, probably due to low affinity to the active site. The fourth class consisted of residues for which no peptidic products could be detected. This lack of detection could be explained by the models: these sites did not bind to the active site, or the resulting products were not detected by MS.

**Proteolysis of fibrillar tau by HTRA1**

Based on histograms (SI Appendix, Fig. S2E-G), P1 sites were grouped into classes according to the maximum relative frequency of cuts performed at each site, class 1 (≥20 cuts, i.e. 25 to 59) comprised 5 residues (SI Appendix, Table S1) and class 2 (10-19 cuts) comprised 9 residues (SI Appendix, Fig. S2B, Table S2). The P1 sites with the highest relative number of cuts are V256, L376, I392, L408 and V411. Of these, L408 and A392 are expected to be cleaved first, as they are associated with the highest number of cuts at 15 sec, i.e. 11 and 10, respectively), followed by L376, which reaches 28 cuts at 1 min of incubation, suggesting that proteolysis is initiated at the C-terminus of tau. The next cleavage should occur after V256, reaching 23 cuts after 5 min of incubation. Since V256 is located upstream of the fibril core, these first 4 cuts could produce a fragment of, or fragments similar to, K257-L376, containing the β-sheet stacks of the fibrils comprising V275-C291, S305-Y310, V313-D314, K317-V318, and N327-I328.

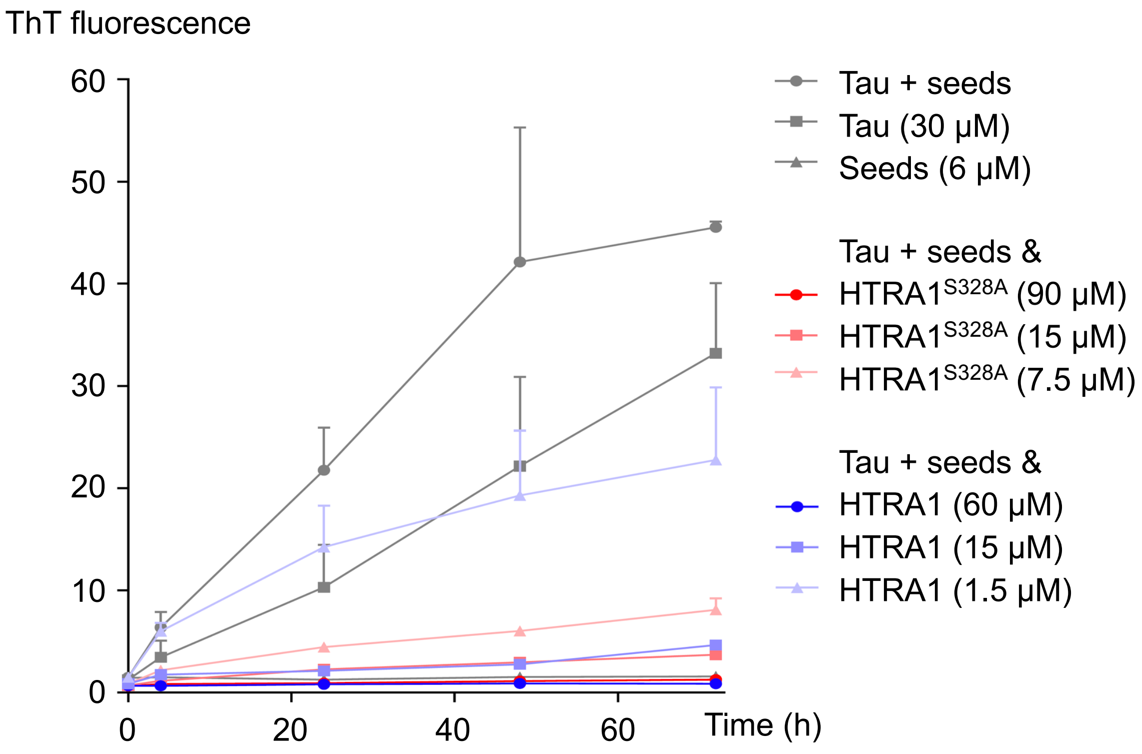


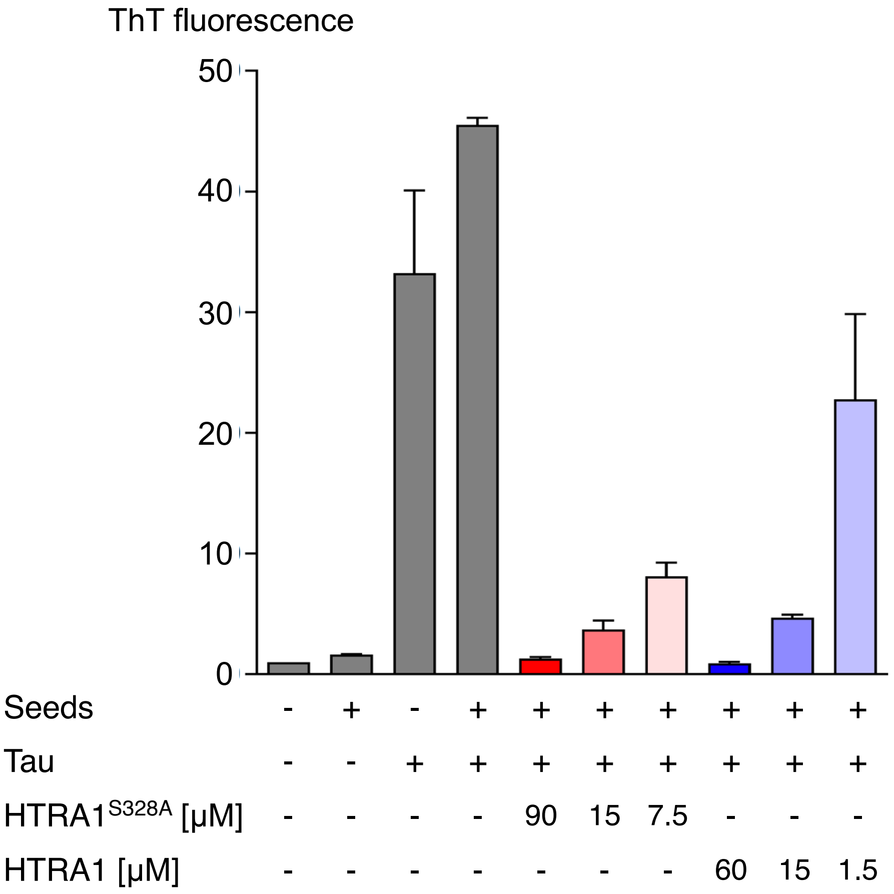


**Fig. S1. Quantification of tau seeding based on ThT fluorescence. Related to Fig.1** Top, In vitro seeding of tau aggregation was performed as described in Fig. 1. ThT fluorescence was measured at 480 nm at the time points indicated. Measured ThT signals were normalized to the corresponding buffer controls. Bottom, For better comparison, endpoint values (72 h) were plotted as bar diagrams. Error bars, s.e.m.; n = 3.

### Fig. S2A Proteolysis of soluble tau by HTRA1

| | 1 | 2 | 3 | 4 | 5 | 6 | 7 | 8 | 9 | 10 | 11 | 12 | 13 | 14 | 15 | 16 | 17 | 18 | 19 | 20 | 21 | 22 | 23 | 24 | 25 | 26 | 27 | 28 | 29 | 30 | 31 | 32 | 33 | 34 | 35 | 36 | 37 | 38 | 39 | 40 | 41 | 42 | 43 | 44 | 45 | 46 | 47 | 48 | 49 | 50 | Residue no. |
|---|---|---|---|---|---|---|---|---|---|---|---|---|---|---|---|---|---|---|---|---|---|---|---|---|---|---|---|---|---|---|---|---|---|---|---|---|---|---|---|---|---|---|---|---|---|---|---|---|---|---|---|
| | M | A | E | P | R | Q | E | F | E | V | M | E | D | H | A | G | T | Y | G | L | G | D | R | K | D | Q | G | G | Y | T | M | H | Q | D | Q | E | G | D | T | D | A | G | L | K | E | S | P | L | Q | T | Sequence |
| Min | * | * | : | * | : | * | * | * | : | . | | * | * | : | . | | | | | | | | | | | | | . | * | | | | * | * | : | * | * | * | | : | | * | * | * | * | * | * | | * | | Conservation |
| 0.25 | 0 | | | | | | | | | 0 | 0 | | | | | | | | | | | | | | | | | | | | | | | | | | | | | | | | | | | | | | | | Rel. freq. cuts |
| 1 | 0 | | | | | | | | | 0 | 0 | | | | | | | | | | | | | | | | | | | | | | | | | | | | | | | | | | | | | | | | |
| 2 | 1 | | | | | | | | | 1 | 0 | | | | | | | | | | | | | | | | | | | | | | | | | | | | | | | | | | | | | | | | |
| 5 | 1 | | | | | | | | | 0 | 1 | | | | | | | | | | | | | | | | | | | | | | | | | | | | | | | | | | | | | | | | |
| 10 | 0 | | | | | | | | | 0 | 0 | | | | | | | | | | | | | | | | | | | | | | | | | | | | | | | | | | | | | | | | |
| 120 | 2 | | | | | | | | | 1 | 1 | | | | | | | | | | | | | | | | | | | | | | | | | | | | | | | | | | | | | | | | |

| | 51 | 52 | 53 | 54 | 55 | 56 | 57 | 58 | 59 | 60 | 61 | 62 | 63 | 64 | 65 | 66 | 67 | 68 | 69 | 70 | 71 | 72 | 73 | 74 | 75 | 76 | 77 | 78 | 79 | 80 | 81 | 82 | 83 | 84 | 85 | 86 | 87 | 88 | 89 | 90 | 91 | 92 | 93 | 94 | 95 | 96 | 97 | 98 | 99 | 100 | Residue no. |
|---|---|---|---|---|---|---|---|---|---|---|---|---|---|---|---|---|---|---|---|---|---|---|---|---|---|---|---|---|---|---|---|---|---|---|---|---|---|---|---|---|---|---|---|---|---|---|---|---|---|---|---|
| | P | T | E | D | G | S | E | E | P | G | S | E | T | S | D | A | K | S | T | P | T | A | E | D | V | T | A | P | L | V | D | E | G | A | P | G | K | Q | A | A | A | Q | P | H | T | E | I | P | E | G | Sequence |
| Min | * | : | : | * | * | : | * | * | | * | * | * | * | * | * | * | * | * | * | * | * | * | * | | | | | | | | | | | | | | | | | | | | | | | | | | | | Conservation |
| 0.25 | | | | | | | | | | | | | | | | | | | | | | | | | | | | | | | | | | | | | | | | | | | | | | | | | | | Rel. freq. cuts |
| 1 | | | | | | | | | | | | | | | | | | | | | | | | | | | | | | | | | | | | | | | | | | | | | | | | | | | |
| 2 | | | | | | | | | | | | | | | | | | | | | | | | | | | | | | | | | | | | | | | | | | | | | | | | | | | |
| 5 | | | | | | | | | | | | | | | | | | | | | | | | | | | | | | | | | | | | | | | | | | | | | | | | | | | |
| 10 | | | | | | | | | | | | | | | | | | | | | | | | | | | | | | | | | | | | | | | | | | | | | | | | | | | |
| 120 | | | | | | | | | | | | | | | | | | | | | | | | | | | | | | | | | | | | | | | | | | | | | | | | | | | |

| | 101 | 102 | 103 | 104 | 105 | 106 | 107 | 108 | 109 | 110 | 111 | 112 | 113 | 114 | 115 | 116 | 117 | 118 | 119 | 120 | 121 | 122 | 123 | 124 | 125 | 126 | 127 | 128 | 129 | 130 | 131 | 132 | 133 | 134 | 135 | 136 | 137 | 138 | 139 | 140 | 141 | 142 | 143 | 144 | 145 | 146 | 147 | 148 | 149 | 150 | Residue no. |
|---|---|---|---|---|---|---|---|---|---|---|---|---|---|---|---|---|---|---|---|---|---|---|---|---|---|---|---|---|---|---|---|---|---|---|---|---|---|---|---|---|---|---|---|---|---|---|---|---|---|---|---|
| | T | T | A | E | E | A | G | I | G | D | T | P | S | L | E | D | E | A | A | G | H | V | T | Q | A | R | M | V | S | K | S | K | D | G | T | G | S | D | D | K | K | A | K | G | A | D | G | K | T | K | Sequence |
| Min | | | * | * | * | * | * | * | * | * | * | | . | | * | * | : | * | * | * | * | * | * | * | * | | | . | | | . | * | * | | * | * | | * | : | | | | | . | . | . | | | : | * | Conservation |
| 0.25 | | | | | | 0 | | | | | | | | | | | | | | | | | | | 0 | | | | | | | | | | | | | | | | | | | | | | | | | | Rel. freq. cuts |
| 1 | | | | | | 0 | | | | | | | | | | | | | | | | | | | 0 | | | | | | | | | | | | | | | | | | | | | | | | | | |
| 2 | | | | | | 0 | | | | | | | | | | | | | | | | | | | 0 | | | | | | | | | | | | | | | | | | | | | | | | | | |
| 5 | | | | | | 0 | | | | | | | | | | | | | | | | | | | 0 | | | | | | | | | | | | | | | | | | | | | | | | | | |
| 10 | | | | | | 0 | | | | | | | | | | | | | | | | | | | 0 | | | | | | | | | | | | | | | | | | | | | | | | | | |
| 120 | | | | | | 1 | | | | | | | | | | | | | | | | | | | 1 | | | | | | | | | | | | | | | | | | | | | | | | | | |

| | 151 | 152 | 153 | 154 | 155 | 156 | 157 | 158 | 159 | 160 | 161 | 162 | 163 | 164 | 165 | 166 | 167 | 168 | 169 | 170 | 171 | 172 | 173 | 174 | 175 | 176 | 177 | 178 | 179 | 180 | 181 | 182 | 183 | 184 | 185 | 186 | 187 | 188 | 189 | 190 | 191 | 192 | 193 | 194 | 195 | 196 | 197 | 198 | 199 | 200 | Residue no. |
|---|---|---|---|---|---|---|---|---|---|---|---|---|---|---|---|---|---|---|---|---|---|---|---|---|---|---|---|---|---|---|---|---|---|---|---|---|---|---|---|---|---|---|---|---|---|---|---|---|---|---|---|
| | I | A | T | P | R | G | A | A | P | P | G | Q | K | G | Q | A | N | A | T | R | I | P | A | K | T | P | P | A | P | K | T | P | P | S | S | G | E | P | P | K | S | G | D | R | S | G | Y | S | S | P | Sequence |
| Min | * | * | * | * | * | * | * | * | | * | | * | * | * | | : | * | * | * | * | * | * | * | * | * | | * | : | * | * | * | | * | | | * | * | | | * | * | * | : | * | * | * | * | * | * | * | Conservation |
| 0.25 | | | | | | | | | | | | | | | | | | | | | | | | | | | | | | | | | | | | | | | | | | | | | | | | | | | Rel. freq. cuts |
| 1 | | | | | | | | | | | | | | | | | | | | | | | | | | | | | | | | | | | | | | | | | | | | | | | | | | | |
| 2 | | | | | | | | | | | | | | | | | | | | | | | | | | | | | | | | | | | | | | | | | | | | | | | | | | | |
| 5 | | | | | | | | | | | | | | | | | | | | | | | | | | | | | | | | | | | | | | | | | | | | | | | | | | | |
| 10 | | | | | | | | | | | | | | | | | | | | | | | | | | | | | | | | | | | | | | | | | | | | | | | | | | | |
| 120 | | | | | | | | | | | | | | | | | | | | | | | | | | | | | | | | | | | | | | | | | | | | | | | | | | | |

| | 201 | 202 | 203 | 204 | 205 | 206 | 207 | 208 | 209 | 210 | 211 | 212 | 213 | 214 | 215 | 216 | 217 | 218 | 219 | 220 | 221 | 222 | 223 | 224 | 225 | 226 | 227 | 228 | 229 | 230 | 231 | 232 | 233 | 234 | 235 | 236 | 237 | 238 | 239 | 240 | 241 | 242 | 243 | 244 | 245 | 246 | 247 | 248 | 249 | 250 | Residue no. |
|---|---|---|---|---|---|---|---|---|---|---|---|---|---|---|---|---|---|---|---|---|---|---|---|---|---|---|---|---|---|---|---|---|---|---|---|---|---|---|---|---|---|---|---|---|---|---|---|---|---|---|---|
| | G | S | P | G | T | P | G | S | R | S | R | T | P | S | L | P | T | P | P | T | R | E | P | K | K | V | A | V | V | R | T | P | P | K | S | P | S | S | A | K | S | R | L | Q | T | A | P | V | P | M | Sequence |
| Min | * | * | * | * | * | * | * | * | * | * | * | * | * | * | * | * | * | * | * | : | * | * | * | * | * | * | * | * | * | * | * | * | * | * | * | * | * | : | : | * | * | * | * | * | : | * | * | | * | * | Conservation |
| 0.25 | | | | | | | | | | | | | | | | | | | | | | | | | | 1 | | 0 | 0 | | | | | | | | 0 | | 0 | | 0 | | 0 | 1 | | | | | | | Rel. freq. cuts |
| 1 | | | | | | | | | | | | | | | | | | | | | | | | | | 1 | | 0 | 0 | | | | | | | | 0 | | 0 | | 0 | | 0 | 3 | | | | | | | |
| 2 | | | | | | | | | | | | | | | | | | | | | | | | | | 3 | | 0 | 0 | | | | | | | | 0 | | 0 | | 0 | | 0 | 4 | | | | | | | |
| 5 | | | | | | | | | | | | | | | | | | | | | | | | | | 2 | | 0 | 0 | | | | | | | | 0 | | 0 | | 0 | | 0 | 3 | | | | | | | |
| 10 | | | | | | | | | | | | | | | | | | | | | | | | | | 3 | | 0 | 1 | | | | | | | | 0 | | 0 | | 0 | | 0 | 4 | | | | | | | |
| 120 | | | | | | | | | | | | | | | | | | | | | | | | | | 5 | | 1 | 3 | | | | | | | | 1 | | 1 | | 1 | | 1 | 7 | | | | | | | |

| | 251 | 252 | 253 | 254 | 255 | 256 | 257 | 258 | 259 | 260 | 261 | 262 | 263 | 264 | 265 | 266 | 267 | 268 | 269 | 270 | 271 | 272 | 273 | 274 | 275 | 276 | 277 | 278 | 279 | 280 | 281 | 282 | 283 | 284 | 285 | 286 | 287 | 288 | 289 | 290 | 291 | 292 | 293 | 294 | 295 | 296 | 297 | 298 | 299 | 300 | Residue no. |
|---|---|---|---|---|---|---|---|---|---|---|---|---|---|---|---|---|---|---|---|---|---|---|---|---|---|---|---|---|---|---|---|---|---|---|---|---|---|---|---|---|---|---|---|---|---|---|---|---|---|---|---|
| | P | D | L | K | N | V | K | S | K | I | G | S | T | E | N | L | K | H | Q | P | G | G | G | K | V | Q | I | I | N | K | K | L | D | L | S | N | V | Q | S | K | C | G | S | K | D | N | I | K | H | V | Sequence |
| Min | * | * | * | * | * | * | : | * | * | * | * | * | * | * | * | * | * | * | * | * | * | * | * | * | * | * | * | * | * | * | * | * | * | * | * | * | * | * | * | * | * | * | * | * | * | * | * | * | * | * | Conservation |
| 0.25 | | | | | | 3 | | | | | | | 0 | | | 1 | | | | | | | | | 0 | | 1 | 1 | | 1 | | 1 | | 0 | 0 | | 0 | | 0 | | 4 | | | | 0 | | | | | | Rel. freq. cuts |
| 1 | | | | | | 5 | | | | | | | 0 | | | 1 | | | | | | | | | 0 | | 2 | 1 | | 1 | | 3 | | 0 | 0 | | 3 | | 0 | | 4 | | | | 0 | | | | | | |
| 2 | | | | | | 6 | | | | | | | 0 | | | 1 | | | | | | | | | 0 | | 5 | 3 | | 3 | | 4 | | 0 | 0 | | 5 | | 3 | | 9 | | | | 0 | | | | | | |
| 5 | | | | | | 6 | | | | | | | 0 | | | 5 | | | | | | | | | 1 | | 5 | 5 | | 2 | | 4 | | 1 | 1 | | 7 | | 0 | | 10 | | | | 0 | | | | | | |
| 10 | | | | | | 8 | | | | | | | 0 | | | 5 | | | | | | | | | 2 | | 8 | 6 | | 3 | | 4 | | 1 | 1 | | 7 | | 4 | | 15 | | | | 1 | | | | | | |
| 120 | | | | | | 22 | | | | | | | 1 | | | 11 | | | | | | | | | 6 | | 13 | 12 | | 7 | | 6 | | 2 | 2 | | 13 | | 10 | | 24 | | | | 2 | | | | | | |

| | 301 | 302 | 303 | 304 | 305 | 306 | 307 | 308 | 309 | 310 | 311 | 312 | 313 | 314 | 315 | 316 | 317 | 318 | 319 | 320 | 321 | 322 | 323 | 324 | 325 | 326 | 327 | 328 | 329 | 330 | 331 | 332 | 333 | 334 | 335 | 336 | 337 | 338 | 339 | 340 | 341 | 342 | 343 | 344 | 345 | 346 | 347 | 348 | 349 | 350 | Residue no. |
|---|---|---|---|---|---|---|---|---|---|---|---|---|---|---|---|---|---|---|---|---|---|---|---|---|---|---|---|---|---|---|---|---|---|---|---|---|---|---|---|---|---|---|---|---|---|---|---|---|---|---|---|
| | P | G | G | G | S | V | Q | I | V | Y | K | P | V | D | L | S | K | V | T | S | K | C | G | S | L | G | N | I | H | H | K | P | G | G | G | Q | V | E | V | K | S | E | K | L | D | F | K | D | R | V | Sequence |
| Min | * | * | * | * | * | * | : | * | * | * | * | * | * | * | * | * | * | * | * | * | * | * | * | * | * | * | * | * | * | * | * | * | * | * | * | * | * | * | * | * | * | * | * | * | * | * | * | * | * | * | Conservation |
| 0.25 | | 0 | | | | 0 | 0 | 4 | 0 | 1 | 0 | | 0 | 0 | 0 | 0 | 0 | 1 | 0 | | 0 | 3 | 0 | 0 | 0 | 0 | 0 | 0 | 0 | 0 | | | | 0 | 0 | | 0 | | 0 | | 0 | 0 | 0 | 2 | 0 | | | | | 0 | Rel. freq. cuts |
| 1 | | 1 | | | | 0 | 0 | 8 | 1 | 0 | 0 | | 2 | 0 | 0 | 1 | 0 | 2 | 1 | | 0 | 5 | 0 | 0 | 0 | 0 | 0 | 2 | 0 | 0 | | | | 0 | 0 | | 0 | | 0 | | 0 | 0 | 0 | 2 | 0 | | | | | 1 | |
| 2 | | 1 | | | | 0 | 0 | 17 | 4 | 0 | 0 | | 4 | 0 | 0 | 2 | 0 | 3 | 2 | | 1 | 11 | 0 | 0 | 1 | 0 | 0 | 5 | 0 | 0 | | | | 0 | 0 | | 0 | | 2 | | 0 | 0 | 0 | 6 | 0 | | | | | 0 | |
| 5 | | 2 | | | | 0 | 0 | 17 | 4 | 0 | 0 | | 4 | 0 | 0 | 2 | 0 | 5 | 2 | | 1 | 10 | 0 | 0 | 1 | 0 | 0 | 5 | 1 | 0 | | | | 0 | 0 | | 0 | | 1 | | 0 | 0 | 0 | 3 | 0 | | | | | 2 | |
| 10 | | 2 | | | | 0 | 0 | 21 | 5 | 0 | 1 | | 5 | 0 | 0 | 2 | 0 | 5 | 2 | | 1 | 14 | 1 | 0 | 1 | 0 | 0 | 6 | 1 | 0 | | | | 0 | 0 | | 0 | | 2 | | 1 | 0 | 0 | 5 | 0 | | | | | 2 | |
| 120 | | 4 | | | | 2 | 1 | 42 | 9 | 2 | 3 | | 8 | 1 | 1 | 3 | 1 | 9 | 3 | | 2 | 32 | 1 | 2 | 5 | 1 | 1 | 13 | 1 | 1 | | | | 1 | 1 | | 1 | | 3 | | 2 | 1 | 1 | 15 | 1 | | | | | 2 | |

| | 351 | 352 | 353 | 354 | 355 | 356 | 357 | 358 | 359 | 360 | 361 | 362 | 363 | 364 | 365 | 366 | 367 | 368 | 369 | 370 | 371 | 372 | 373 | 374 | 375 | 376 | 377 | 378 | 379 | 380 | 381 | 382 | 383 | 384 | 385 | 386 | 387 | 388 | 389 | 390 | 391 | 392 | 393 | 394 | 395 | 396 | 397 | 398 | 399 | 400 | Residue no. |
|---|---|---|---|---|---|---|---|---|---|---|---|---|---|---|---|---|---|---|---|---|---|---|---|---|---|---|---|---|---|---|---|---|---|---|---|---|---|---|---|---|---|---|---|---|---|---|---|---|---|---|---|
| | Q | S | K | I | G | S | L | D | N | I | T | H | V | P | G | G | G | N | K | K | I | E | T | H | K | L | T | F | R | E | N | A | K | A | K | T | D | H | G | A | E | I | V | Y | K | S | P | V | V | S | Sequence |
| Min | * | * | * | * | * | * | * | * | * | * | * | * | * | * | * | * | * | : | * | * | * | * | * | * | * | * | * | * | * | * | * | * | * | * | * | * | * | * | * | * | * | * | * | * | * | * | * | * | * | * | Conservation |
| 0.25 | | | | | | | 0 | | | 2 | 0 | | 0 | 0 | 0 | 0 | | | | | | | | | | 5 | 0 | | | | | | | | | | 0 | 0 | | 0 | | 12 | 1 | 1 | 1 | | | 0 | 0 | | Rel. freq. cuts |
| 1 | | | | | | | 0 | | | 4 | 0 | | 0 | 0 | 0 | 0 | | | | | | | | | | 6 | 0 | | | | | | | | | | 0 | 0 | | 0 | | 24 | 1 | 1 | 1 | | | 0 | 0 | | |
| 2 | | | | | | | 0 | | | 4 | 0 | | 0 | 0 | 2 | 0 | | | | | | | | | | 9 | 0 | | | | | | | | | | 1 | 0 | | 0 | | 39 | 3 | 1 | 0 | | | 2 | 0 | | |
| 5 | | | | | | | 0 | | | 5 | 0 | | 0 | 0 | 0 | 0 | | | | | | | | | | 8 | 0 | | | | | | | | | | 0 | 0 | | 0 | | 44 | 2 | 1 | 2 | | | 1 | 1 | | |
| 10 | | | | | | | 1 | | | 5 | 0 | | 1 | 0 | 1 | 0 | | | | | | | | | | 12 | 1 | | | | | | | | | | 1 | 1 | | 0 | | 47 | 3 | 1 | 1 | | | 3 | 1 | | |
| 120 | | | | | | | 3 | | | 6 | 1 | | 2 | 1 | 3 | 1 | | | | | | | | | | 27 | 2 | | | | | | | | | | 2 | 2 | | 1 | | 51 | 4 | 1 | 2 | | | 6 | 1 | | |

| | 401 | 402 | 403 | 404 | 405 | 406 | 407 | 408 | 409 | 410 | 411 | 412 | 413 | 414 | 415 | 416 | 417 | 418 | 419 | 420 | 421 | 422 | 423 | 424 | 425 | 426 | 427 | 428 | 429 | 430 | 431 | 432 | 433 | 434 | 435 | 436 | 437 | 438 | 439 | 440 | 441 | | | | | | | | | | Residue no. |
|---|---|---|---|---|---|---|---|---|---|---|---|---|---|---|---|---|---|---|---|---|---|---|---|---|---|---|---|---|---|---|---|---|---|---|---|---|---|---|---|---|---|---|---|---|---|---|---|---|---|---|---|
| | G | D | T | S | P | R | H | L | S | N | V | S | S | T | G | S | I | D | M | V | D | S | P | Q | L | A | T | L | A | D | E | V | S | A | S | L | A | K | Q | G | L | | | | | | | | | | Sequence |
| Min | * | * | * | * | * | * | * | * | * | * | * | * | * | * | * | * | * | * | * | * | * | * | * | * | * | * | * | * | * | * | * | * | * | * | * | * | * | * | * | * | * | | | | | | | | | | Conservation |
| 0.25 | 1 | 0 | 0 | 0 | 0 | 0 | 0 | 7 | 1 | 1 | 1 | 1 | 0 | 0 | 1 | 0 | | 0 | 1 | 0 | 1 | 0 | 0 | 0 | 3 | 2 | 1 | 3 | | 0 | 1 | 2 | | 1 | | | | | 0 | 0 | | | | | | | | | | | Rel. freq. cuts |
| 1 | 1 | 1 | 1 | 1 | 0 | 1 | 1 | 15 | 2 | 2 | 4 | 2 | 0 | 1 | 1 | 0 | | 1 | 2 | 2 | 2 | 1 | 2 | 2 | 4 | 3 | 1 | 4 | | 0 | 0 | 4 | | 2 | | | | | 0 | 0 | | | | | | | | | | | |
| 2 | 1 | 2 | 2 | 2 | 2 | 2 | 2 | 28 | 2 | 2 | 10 | 2 | 1 | 5 | 2 | 1 | | 1 | 3 | 3 | 3 | 2 | 3 | 2 | 4 | 8 | 4 | 10 | | 2 | 2 | 8 | | 1 | | | | | 1 | 1 | | | | | | | | | | | |
| 5 | 1 | 1 | 1 | 2 | 1 | 2 | 1 | 22 | 2 | 2 | 8 | 2 | 1 | 2 | 2 | 1 | | 2 | 3 | 3 | 3 | 2 | 4 | 4 | 5 | 6 | 2 | 5 | | 1 | 1 | 6 | | 2 | | | | | 0 | 0 | | | | | | | | | | | |
| 10 | 1 | 2 | 2 | 2 | 1 | 2 | 2 | 28 | 2 | 2 | 12 | 2 | 1 | 4 | 2 | 2 | | 2 | 3 | 4 | 3 | 2 | 4 | 3 | 5 | 7 | 4 | 7 | | 2 | 2 | 8 | | 3 | | | | | 1 | 0 | | | | | | | | | | | |
| 120 | 2 | 2 | 2 | 2 | 0 | 3 | 2 | 29 | 3 | 3 | 16 | 2 | 1 | 9 | 3 | 2 | | 2 | 3 | 5 | 5 | 3 | 5 | 4 | 6 | 9 | 2 | 13 | | 3 | 2 | 12 | | 3 | | | | | 1 | 1 | | | | | | | | | | | |

### Fig. S2B Proteolysis of fibrillar tau by HTRA1

| | 1 | 2 | 3 | 4 | 5 | 6 | 7 | 8 | 9 | 10 | 11 | 12 | 13 | 14 | 15 | 16 | 17 | 18 | 19 | 20 | 21 | 22 | 23 | 24 | 25 | 26 | 27 | 28 | 29 | 30 | 31 | 32 | 33 | 34 | 35 | 36 | 37 | 38 | 39 | 40 | 41 | 42 | 43 | 44 | 45 | 46 | 47 | 48 | 49 | 50 | Residue no. |
|---|---|---|---|---|---|---|---|---|---|---|---|---|---|---|---|---|---|---|---|---|---|---|---|---|---|---|---|---|---|---|---|---|---|---|---|---|---|---|---|---|---|---|---|---|---|---|---|---|---|---|---|
| | M | A | E | P | R | Q | E | F | E | V | M | E | D | H | A | G | T | Y | G | L | G | D | R | K | D | Q | G | G | Y | T | M | H | Q | D | Q | E | G | D | T | D | A | G | L | K | E | S | P | L | Q | T | Sequence |
| Min | * | * | : | * | : | * | * | * | : | . | | * | * | : | . | | | | | | | | | | | | | . | * | | | | * | * | : | * | * | * | | : | | * | * | * | * | * | * | | * | | Conservation |
| 0.25 | 0 | | | | | | | | | | | | | | | | | | | 0 | | | | | | | | | | | | 0 | | | | | | | | | | | | | | | | | | | Rel. freq. cuts |
| 1 | 0 | | | | | | | | | | | | | | | | | | | 0 | | | | | | | | | | | | 0 | | | | | | | | | | | | | | | | | | | |
| 2 | 0 | | | | | | | | | | | | | | | | | | | 0 | | | | | | | | | | | | 0 | | | | | | | | | | | | | | | | | | | |
| 5 | 1 | | | | | | | | | | | | | | | | | | | 0 | | | | | | | | | | | | 1 | | | | | | | | | | | | | | | | | | | |
| 10 | 2 | | | | | | | | | | | | | | | | | | | 1 | | | | | | | | | | | | 1 | | | | | | | | | | | | | | | | | | | |
| 120 | 0 | | | | | | | | | | | | | | | | | | | 0 | | | | | | | | | | | | 0 | | | | | | | | | | | | | | | | | | | |

| | 51 | 52 | 53 | 54 | 55 | 56 | 57 | 58 | 59 | 60 | 61 | 62 | 63 | 64 | 65 | 66 | 67 | 68 | 69 | 70 | 71 | 72 | 73 | 74 | 75 | 76 | 77 | 78 | 79 | 80 | 81 | 82 | 83 | 84 | 85 | 86 | 87 | 88 | 89 | 90 | 91 | 92 | 93 | 94 | 95 | 96 | 97 | 98 | 99 | 100 | Residue no. |
|---|---|---|---|---|---|---|---|---|---|---|---|---|---|---|---|---|---|---|---|---|---|---|---|---|---|---|---|---|---|---|---|---|---|---|---|---|---|---|---|---|---|---|---|---|---|---|---|---|---|---|---|
| | P | T | E | D | G | S | E | E | P | G | S | E | T | S | D | A | K | S | T | P | T | A | E | D | V | T | A | P | L | V | D | E | G | A | P | G | K | Q | A | A | A | Q | P | H | T | E | I | P | E | G | Sequence |
| Min | * | : | : | * | * | : | * | * | | * | * | * | * | * | * | * | * | * | * | * | * | * | * | | | | | | | | | | | | | | | | | | | | | | | | | | | | Conservation |
| 0.25 | | | | | | | | | | | | | | | | | | | | | | | | | | | | | | | | | | | | | | | | | | | | | | | | | | | Rel. freq. cuts |
| 1 | | | | | | | | | | | | | | | | | | | | | | | | | | | | | | | | | | | | | | | | | | | | | | | | | | | |
| 2 | | | | | | | | | | | | | | | | | | | | | | | | | | | | | | | | | | | | | | | | | | | | | | | | | | | |
| 5 | | | | | | | | | | | | | | | | | | | | | | | | | | | | | | | | | | | | | | | | | | | | | | | | | | | |
| 10 | | | | | | | | | | | | | | | | | | | | | | | | | | | | | | | | | | | | | | | | | | | | | | | | | | | |
| 120 | | | | | | | | | | | | | | | | | | | | | | | | | | | | | | | | | | | | | | | | | | | | | | | | | | | |

| | 101 | 102 | 103 | 104 | 105 | 106 | 107 | 108 | 109 | 110 | 111 | 112 | 113 | 114 | 115 | 116 | 117 | 118 | 119 | 120 | 121 | 122 | 123 | 124 | 125 | 126 | 127 | 128 | 129 | 130 | 131 | 132 | 133 | 134 | 135 | 136 | 137 | 138 | 139 | 140 | 141 | 142 | 143 | 144 | 145 | 146 | 147 | 148 | 149 | 150 | Residue no. |
|---|---|---|---|---|---|---|---|---|---|---|---|---|---|---|---|---|---|---|---|---|---|---|---|---|---|---|---|---|---|---|---|---|---|---|---|---|---|---|---|---|---|---|---|---|---|---|---|---|---|---|---|
| | T | T | A | E | E | A | G | I | G | D | T | P | S | L | E | D | E | A | A | G | H | V | T | Q | A | R | M | V | S | K | S | K | D | G | T | G | S | D | D | K | K | A | K | G | A | D | G | K | T | K | Sequence |
| Min | | | * | * | * | * | * | * | * | * | * | | . | | * | * | : | * | * | * | * | * | * | * | * | | | . | | | . | * | * | | * | * | | * | : | | | | | . | . | . | | | : | * | Conservation |
| 0.25 | | | | | | 0 | | | | | | 0 | | | | | | | | | | | | | 0 | | | | | | | | | | | | | | | | | | | | | | | | | | Rel. freq. cuts |
| 1 | | | | | | 0 | | | | | | 0 | | | | | | | | | | | | | 0 | | | | | | | | | | | | | | | | | | | | | | | | | | |
| 2 | | | | | | 0 | | | | | | 0 | | | | | | | | | | | | | 0 | | | | | | | | | | | | | | | | | | | | | | | | | | |
| 5 | | | | | | 1 | | | | | | 0 | | | | | | | | | | | | | 1 | | | | | | | | | | | | | | | | | | | | | | | | | | |
| 10 | | | | | | 2 | | | | | | 0 | | | | | | | | | | | | | 2 | | | | | | | | | | | | | | | | | | | | | | | | | | |
| 120 | | | | | | 2 | | | | | | 1 | | | | | | | | | | | | | 3 | | | | | | | | | | | | | | | | | | | | | | | | | | |

| | 151 | 152 | 153 | 154 | 155 | 156 | 157 | 158 | 159 | 160 | 161 | 162 | 163 | 164 | 165 | 166 | 167 | 168 | 169 | 170 | 171 | 172 | 173 | 174 | 175 | 176 | 177 | 178 | 179 | 180 | 181 | 182 | 183 | 184 | 185 | 186 | 187 | 188 | 189 | 190 | 191 | 192 | 193 | 194 | 195 | 196 | 197 | 198 | 199 | 200 | Residue no. |
|---|---|---|---|---|---|---|---|---|---|---|---|---|---|---|---|---|---|---|---|---|---|---|---|---|---|---|---|---|---|---|---|---|---|---|---|---|---|---|---|---|---|---|---|---|---|---|---|---|---|---|---|
| | I | A | T | P | R | G | A | A | P | P | G | Q | K | G | Q | A | N | A | T | R | I | P | A | K | T | P | P | A | P | K | T | P | P | S | S | G | E | P | P | K | S | G | D | R | S | G | Y | S | S | P | Sequence |
| Min | * | * | * | * | * | * | * | * | | * | | * | * | * | | : | * | * | * | * | * | * | * | * | * | | * | : | * | * | * | | * | | | * | * | | | * | * | * | : | * | * | * | * | * | * | * | Conservation |
| 0.25 | | | | | | | | | | | | | | | | | | | | | | | | | | | | | | | | | | | | | | | | | | | | | | | | | | | Rel. freq. cuts |
| 1 | | | | | | | | | | | | | | | | | | | | | | | | | | | | | | | | | | | | | | | | | | | | | | | | | | | |
| 2 | | | | | | | | | | | | | | | | | | | | | | | | | | | | | | | | | | | | | | | | | | | | | | | | | | | |
| 5 | | | | | | | | | | | | | | | | | | | | | | | | | | | | | | | | | | | | | | | | | | | | | | | | | | | |
| 10 | | | | | | | | | | | | | | | | | | | | | | | | | | | | | | | | | | | | | | | | | | | | | | | | | | | |
| 120 | | | | | | | | | | | | | | | | | | | | | | | | | | | | | | | | | | | | | | | | | | | | | | | | | | | |

| | 201 | 202 | 203 | 204 | 205 | 206 | 207 | 208 | 209 | 210 | 211 | 212 | 213 | 214 | 215 | 216 | 217 | 218 | 219 | 220 | 221 | 222 | 223 | 224 | 225 | 226 | 227 | 228 | 229 | 230 | 231 | 232 | 233 | 234 | 235 | 236 | 237 | 238 | 239 | 240 | 241 | 242 | 243 | 244 | 245 | 246 | 247 | 248 | 249 | 250 | Residue no. |
|---|---|---|---|---|---|---|---|---|---|---|---|---|---|---|---|---|---|---|---|---|---|---|---|---|---|---|---|---|---|---|---|---|---|---|---|---|---|---|---|---|---|---|---|---|---|---|---|---|---|---|---|
| | G | S | P | G | T | P | G | S | R | S | R | T | P | S | L | P | T | P | P | T | R | E | P | K | K | V | A | V | V | R | T | P | P | K | S | P | S | S | A | K | S | R | L | Q | T | A | P | V | P | M | Sequence |
| Min | * | * | * | * | * | * | * | * | * | * | * | * | * | * | * | * | * | * | * | : | * | * | * | * | * | * | * | * | * | * | * | * | * | * | * | * | * | : | : | * | * | * | * | * | : | * | * | | * | * | Conservation |
| 0.25 | | | | | | | 0 | | | | 0 | | | | | | | | | | | | | | | 0 | 0 | 0 | 0 | | | | | 0 | | 0 | 0 | | 0 | 0 | 0 | 0 | 0 | 1 | 0 | 0 | 0 | | | | Rel. freq. cuts |
| 1 | | | | | | | 0 | | | | 0 | | | | | | | | | | | | | | | 2 | 0 | 0 | 1 | | | | | 0 | | 0 | 0 | | 1 | 0 | 0 | 0 | 0 | 2 | 1 | 0 | 0 | | | | |
| 2 | | | | | | | 0 | | | | 0 | | | | | | | | | | | | | | | 2 | 0 | 0 | 1 | | | | | 0 | | 0 | 1 | | 1 | 0 | 1 | 1 | 1 | 3 | 2 | 0 | 0 | | | | |
| 5 | | | | | | | 0 | | | | 0 | | | | | | | | | | | | | | | 6 | 0 | 1 | 2 | | | | | 1 | | 1 | 3 | | 2 | 0 | 4 | 1 | 1 | 5 | 2 | 0 | 0 | | | | |
| 10 | | | | | | | 0 | | | | 1 | | | | | | | | | | | | | | | 8 | 0 | 1 | 3 | | | | | 2 | | 1 | 3 | | 3 | 0 | 5 | 1 | 2 | 6 | 2 | 0 | 1 | | | | |
| 120 | | | | | | | 1 | | | | 2 | | | | | | | | | | | | | | | 15 | 1 | 2 | 3 | | | | | 2 | | 2 | 4 | | 6 | 1 | 4 | 2 | 3 | 8 | 4 | 1 | 2 | | | | |

| | 251 | 252 | 253 | 254 | 255 | 256 | 257 | 258 | 259 | 260 | 261 | 262 | 263 | 264 | 265 | 266 | 267 | 268 | 269 | 270 | 271 | 272 | 273 | 274 | 275 | 276 | 277 | 278 | 279 | 280 | 281 | 282 | 283 | 284 | 285 | 286 | 287 | 288 | 289 | 290 | 291 | 292 | 293 | 294 | 295 | 296 | 297 | 298 | 299 | 300 | Residue no. |
|---|---|---|---|---|---|---|---|---|---|---|---|---|---|---|---|---|---|---|---|---|---|---|---|---|---|---|---|---|---|---|---|---|---|---|---|---|---|---|---|---|---|---|---|---|---|---|---|---|---|---|---|
| | P | D | L | K | N | V | K | S | K | I | G | S | T | E | N | L | K | H | Q | P | G | G | G | K | V | Q | I | I | N | K | K | L | D | L | S | N | V | Q | S | K | C | G | S | K | D | N | I | K | H | V | Sequence |
| | | | | | | | | | | | | | | | | | | | | | | G | G | K | V | Q | I | I | N | K | K | L | D | L | S | N | V | Q | S | K | C | G | S | K | D | N | I | K | H | V | B-factors |
| | | | | | | | | | | | | | | | | | | | | | | | | | b | b | b | b | b | b | b | b | b | b | b | b | b | b | b | b | b | | | | | | | | | | Fib. β sheet |
| Min | * | * | * | * | * | * | : | * | * | * | * | * | * | * | * | * | * | * | * | * | * | * | * | * | * | * | * | * | * | * | * | * | * | * | * | * | * | * | * | * | * | * | * | * | * | * | * | * | * | * | Conservation |
| 0.25 | | | | | | 2 | | | | | | | 0 | | | 1 | | | | | | | | | 0 | 0 | 0 | 0 | | | | | | | 0 | | 0 | | 0 | | 0 | | | | | | 0 | | | | Rel. freq. cuts |
| 1 | | | | | | 7 | | | | | | | 0 | | | 5 | | | | | | | | | 0 | 0 | 2 | 2 | | | | | | | 0 | | 2 | | 0 | | 1 | | | | | | 1 | | | | |
| 2 | | | | | | 13 | | | | | | | 0 | | | 5 | | | | | | | | | 0 | 0 | 2 | 2 | | | | | | | 0 | | 2 | | 0 | | 1 | | | | | | 0 | | | | |
| 5 | | | | | | 23 | | | | | | | 0 | | | 11 | | | | | | | | | 1 | 1 | 4 | 6 | | | | | | | 1 | | 7 | | 0 | | 1 | | | | | | 3 | | | | |
| 10 | | | | | | 30 | | | | | | | 0 | | | 14 | | | | | | | | | 1 | 1 | 6 | 8 | | | | | | | 2 | | 9 | | 2 | | 3 | | | | | | 4 | | | | |
| 120 | | | | | | 40 | | | | | | | 4 | | | 15 | | | | | | | | | 2 | 2 | 7 | 11 | | | | | | | 2 | | 9 | | 4 | | 2 | | | | | | 2 | | | | |

| | 301 | 302 | 303 | 304 | 305 | 306 | 307 | 308 | 309 | 310 | 311 | 312 | 313 | 314 | 315 | 316 | 317 | 318 | 319 | 320 | 321 | 322 | 323 | 324 | 325 | 326 | 327 | 328 | 329 | 330 | 331 | 332 | 333 | 334 | 335 | 336 | 337 | 338 | 339 | 340 | 341 | 342 | 343 | 344 | 345 | 346 | 347 | 348 | 349 | 350 | Residue no. |
|---|---|---|---|---|---|---|---|---|---|---|---|---|---|---|---|---|---|---|---|---|---|---|---|---|---|---|---|---|---|---|---|---|---|---|---|---|---|---|---|---|---|---|---|---|---|---|---|---|---|---|---|
| | P | G | G | G | S | V | Q | I | V | Y | K | P | V | D | L | S | K | V | T | S | K | C | G | S | L | G | N | I | H | H | K | P | G | G | G | Q | V | E | V | K | S | E | K | L | D | F | K | D | R | V | Sequence |
| | P | G | G | G | S | V | Q | I | V | Y | K | P | V | D | L | S | K | V | T | S | K | C | G | S | L | G | N | I | H | H | | | | | | | | | | | | | | | | | | | | | B-factors |
| | | | | | b | b | b | b | b | b | | | b | b | | | b | b | | | | | | | | | b | b | | | | | | | | | | | | | | | | | | | | | | | Fib. β sheet |
| Min | * | * | * | * | * | * | : | * | * | * | * | * | * | * | * | * | * | * | * | * | * | * | * | * | * | * | * | * | * | * | * | * | * | * | * | * | * | * | * | * | * | * | * | * | * | * | * | * | * | * | Conservation |
| 0.25 | | | | | | | | 1 | 0 | | | | | | 0 | 0 | 0 | 1 | 0 | 0 | 0 | 0 | | | 0 | | | 0 | | | | | | | | | | | 0 | | | 0 | | 0 | | | | | 0 | 0 | Rel. freq. cuts |
| 1 | | | | | | | | 7 | 1 | | | | | | 0 | 0 | 1 | 2 | 1 | 0 | 1 | 1 | | | 0 | | | 0 | | | | | | | | | | | 0 | | | 0 | | 1 | | | | | 0 | 1 | |
| 2 | | | | | | | | 7 | 1 | | | | | | 0 | 0 | 1 | 3 | 1 | 0 | 1 | 2 | | | 0 | | | 0 | | | | | | | | | | | 0 | | | 0 | | 1 | | | | | 0 | 1 | |
| 5 | | | | | | | | 15 | 2 | | | | | | 1 | 1 | 2 | 4 | 1 | 1 | 2 | 4 | | | 0 | | | 1 | | | | | | | | | | | 1 | | | 1 | | 11 | | | | | 0 | 3 | |
| 10 | | | | | | | | 17 | 2 | | | | | | 1 | 1 | 2 | 4 | 2 | 2 | 2 | 5 | | | 1 | | | 1 | | | | | | | | | | | 2 | | | 2 | | 18 | | | | | 1 | 4 | |
| 120 | | | | | | | | 17 | 2 | | | | | | 1 | 1 | 2 | 4 | 2 | 2 | 2 | 6 | | | 1 | | | 1 | | | | | | | | | | | 3 | | | 1 | | 18 | | | | | 0 | 5 | |

| | 351 | 352 | 353 | 354 | 355 | 356 | 357 | 358 | 359 | 360 | 361 | 362 | 363 | 364 | 365 | 366 | 367 | 368 | 369 | 370 | 371 | 372 | 373 | 374 | 375 | 376 | 377 | 378 | 379 | 380 | 381 | 382 | 383 | 384 | 385 | 386 | 387 | 388 | 389 | 390 | 391 | 392 | 393 | 394 | 395 | 396 | 397 | 398 | 399 | 400 | Residue no. |
|---|---|---|---|---|---|---|---|---|---|---|---|---|---|---|---|---|---|---|---|---|---|---|---|---|---|---|---|---|---|---|---|---|---|---|---|---|---|---|---|---|---|---|---|---|---|---|---|---|---|---|---|
| | Q | S | K | I | G | S | L | D | N | I | T | H | V | P | G | G | G | N | K | K | I | E | T | H | K | L | T | F | R | E | N | A | K | A | K | T | D | H | G | A | E | I | V | Y | K | S | P | V | V | S | Sequence |
| Min | * | * | * | * | * | * | * | * | * | * | * | * | * | * | * | * | * | : | * | * | * | * | * | * | * | * | * | * | * | * | * | * | * | * | * | * | * | * | * | * | * | * | * | * | * | * | * | * | * | * | Conservation |
| 0.25 | 0 | 0 | 0 | 0 | 0 | 0 | 0 | 0 | 0 | 1 | 0 | 0 | 0 | 0 | 0 | 0 | 0 | 0 | | | | | | 0 | 0 | 5 | 0 | | | | | 0 | | 1 | | | 0 | 1 | | 0 | 0 | 10 | 1 | 0 | 0 | 0 | | 0 | 0 | 0 | Rel. freq. cuts |
| 1 | 0 | 0 | 0 | 0 | 0 | 0 | 1 | 0 | 0 | 4 | 0 | 0 | 1 | 0 | 1 | 1 | 0 | 0 | | | | | | 0 | 0 | 28 | 1 | | | | | 0 | | 1 | | | 1 | 3 | | 0 | 1 | 34 | 4 | 1 | 1 | 0 | | 4 | 2 | 3 | |
| 2 | 1 | 0 | 0 | 0 | 0 | 0 | 2 | 0 | 0 | 7 | 0 | 0 | 1 | 0 | 2 | 1 | 0 | 1 | | | | | | 0 | 0 | 22 | 1 | | | | | 0 | | 1 | | | 1 | 3 | | 0 | 1 | 38 | 5 | 1 | 1 | 0 | | 6 | 1 | 4 | |
| 5 | 2 | 0 | 1 | 1 | 1 | 0 | 4 | 0 | 1 | 11 | 2 | 1 | 1 | 1 | 3 | 0 | 0 | 3 | | | | | | 1 | 1 | 50 | 2 | | | | | 0 | | 1 | | | 1 | 3 | | 1 | 2 | 50 | 7 | 2 | 1 | 1 | | 11 | 5 | 5 | |
| 10 | 3 | 1 | 1 | 1 | 1 | 1 | 6 | 0 | 1 | 14 | 2 | 1 | 1 | 1 | 3 | 2 | 1 | 1 | | | | | | 0 | 2 | 62 | 4 | | | | | 1 | | 2 | | | 1 | 2 | | 0 | 2 | 51 | 8 | 3 | 2 | 2 | | 12 | 7 | 7 | |
| 120 | 4 | 1 | 0 | 1 | 1 | 2 | 6 | 1 | 1 | 14 | 2 | 2 | 3 | 1 | 3 | 2 | 0 | 2 | | | | | | 1 | 1 | 59 | 3 | | | | | 0 | | 1 | | | 1 | 2 | | 1 | 1 | 49 | 7 | 3 | 2 | 1 | | 12 | 6 | 8 | |

| | 401 | 402 | 403 | 404 | 405 | 406 | 407 | 408 | 409 | 410 | 411 | 412 | 413 | 414 | 415 | 416 | 417 | 418 | 419 | 420 | 421 | 422 | 423 | 424 | 425 | 426 | 427 | 428 | 429 | 430 | 431 | 432 | 433 | 434 | 435 | 436 | 437 | 438 | 439 | 440 | 441 | Residue no. |
|---|---|---|---|---|---|---|---|---|---|---|---|---|---|---|---|---|---|---|---|---|---|---|---|---|---|---|---|---|---|---|---|---|---|---|---|---|---|---|---|---|---|---|
| | G | D | T | S | P | R | H | L | S | N | V | S | S | T | G | S | I | D | M | V | D | S | P | Q | L | A | T | L | A | D | E | V | S | A | S | L | A | K | Q | G | L | Sequence |
| Min | * | * | * | * | * | * | * | * | * | * | * | * | * | * | * | * | * | * | * | * | * | * | * | * | * | * | * | * | * | * | * | * | * | * | * | * | * | * | * | * | * | Conservation |
| 0.25 | 1 | 1 | 1 | 1 | 1 | 0 | 1 | 11 | 0 | 1 | 5 | 0 | 0 | 0 | 0 | 0 | | 0 | 1 | 0 | 1 | 0 | 1 | 1 | 2 | 2 | 0 | 3 | 0 | 0 | 0 | 2 | 1 | 1 | 0 | | | 0 | | 1 | | Rel. freq. cuts |
| 1 | 2 | 2 | 2 | 3 | 3 | 1 | 4 | 24 | 1 | 3 | 13 | 1 | 1 | 3 | 1 | 1 | | 1 | 2 | 2 | 2 | 1 | 2 | 4 | 4 | 5 | 2 | 10 | 0 | 1 | 0 | 4 | 1 | 2 | 1 | | | 1 | | 1 | | |
| 2 | 2 | 2 | 2 | 2 | 2 | 1 | 3 | 30 | 1 | 2 | 18 | 1 | 1 | 5 | 1 | 0 | | 3 | 4 | 4 | 3 | 1 | 3 | 4 | 5 | 7 | 3 | 13 | 2 | 2 | 1 | 8 | 1 | 3 | 1 | | | 1 | | 1 | | |
| 5 | 3 | 3 | 3 | 4 | 4 | 2 | 4 | 37 | 2 | 5 | 24 | 2 | 1 | 9 | 1 | 2 | | 5 | 4 | 7 | 3 | 2 | 4 | 6 | 7 | 9 | 4 | 15 | 2 | 2 | 1 | 13 | 2 | 4 | 3 | | | 1 | | 1 | | |
| 10 | 4 | 4 | 3 | 5 | 3 | 2 | 5 | 40 | 3 | 6 | 26 | 3 | 0 | 9 | 1 | 2 | | 5 | 5 | 7 | 3 | 3 | 4 | 7 | 7 | 11 | 4 | 17 | 1 | 2 | 2 | 15 | 2 | 5 | 3 | | | 1 | | 1 | | |
| 120 | 4 | 3 | 4 | 3 | 3 | 2 | 5 | 40 | 3 | 4 | 25 | 2 | 1 | 9 | 2 | 0 | | 3 | 5 | 8 | 3 | 3 | 4 | 5 | 6 | 11 | 3 | 15 | 1 | 2 | 1 | 14 | 2 | 5 | 3 | | | 1 | | 1 | | |

### Fig. S2C Proteolysis of fibrillar tau CN by HTRA1.

| wt tau | 1 | 2 | 3 | 4 | 5 | 6 | 7 | 8 | 9 | 10 | 11 | 12 | 13 | 14 | 15 | 16 | 17 | 18 | 19 | 20 | 21 | 22 | 23 | 24 | 25 | 26 | 27 | 28 | 29 | 30 | 31 | 32 | 33 | 34 | 35 | 36 | 37 | 38 | 39 | 40 | 41 | 42 | 43 | 44 | 45 | 46 | 47 | 48 | 49 | 50 | Residue no. |
|---|---|---|---|---|---|---|---|---|---|---|---|---|---|---|---|---|---|---|---|---|---|---|---|---|---|---|---|---|---|---|---|---|---|---|---|---|---|---|---|---|---|---|---|---|---|---|---|---|---|---|---|
| | M | A | E | P | R | Q | E | F | E | V | M | E | D | H | A | G | T | Y | G | L | G | D | R | K | D | Q | G | G | Y | T | M | H | Q | D | Q | E | G | D | T | D | A | E | I | V | Y | K | S | P | V | V | Sequence |
| Min | 1 | 2 | 3 | 4 | 5 | 6 | 7 | 8 | 9 | 10 | 11 | 12 | 13 | 14 | 15 | 16 | 17 | 18 | 19 | 20 | 21 | 22 | 23 | 24 | 25 | 26 | 27 | 28 | 29 | 30 | 31 | 32 | 33 | 34 | 35 | 36 | 37 | 38 | 39 | 40 | 41 | 391 | 392 | 393 | 394 | 395 | 396 | 397 | 398 | 399 | Residue no. wt tau |
| 0 | 0 | | | | | | | | | 0 | 0 | | | | | | | | | | | | | | | | | | | | | | | | | | | | | | | 0 | 0 | | 0 | | | 0 | 0 | Rel. freq. cuts |
| 0.25 | 1 | | | | | | | | | 1 | 0 | | | | | | | | | | | | | | | | | | | | | | | | | | | | | | | 4 | 0 | | 0 | | | 0 | 0 | |
| 1 | 1 | | | | | | | | | 1 | 0 | | | | | | | | | | | | | | | | | | | | | | | | | | | | | | | 8 | 1 | | 0 | | | 1 | 1 | |
| 10 | 1 | | | | | | | | | 1 | 0 | | | | | | | | | | | | | | | | | | | | | | | | | | | | | | | 6 | 0 | | 0 | | | 3 | 1 | |
| 2 | 0 | | | | | | | | | 0 | 0 | | | | | | | | | | | | | | | | | | | | | | | | | | | | | | | 11 | 2 | | 0 | | | 2 | 1 | |
| 5 | 0 | | | | | | | | | 0 | 0 | | | | | | | | | | | | | | | | | | | | | | | | | | | | | | | 14 | 3 | | 1 | | | 4 | 3 | |
| 120 | 1 | | | | | | | | | 0 | 1 | | | | | | | | | | | | | | | | | | | | | | | | | | | | | | | 15 | 3 | | 1 | | | 6 | 3 | |

| wt tau | 51 | 52 | 53 | 54 | 55 | 56 | 57 | 58 | 59 | 60 | 61 | 62 | 63 | 64 | 65 | 66 | 67 | 68 | 69 | 70 | 71 | 72 | 73 | 74 | 75 | 76 | 77 | 78 | 79 | 80 | 81 | 82 | 83 | 84 | 85 | 86 | 87 | 88 | 89 | 90 | 91 | 92 | 93 | 94 | 95 | 96 | 97 | 98 | 99 | 100 | Residue no. |
|---|---|---|---|---|---|---|---|---|---|---|---|---|---|---|---|---|---|---|---|---|---|---|---|---|---|---|---|---|---|---|---|---|---|---|---|---|---|---|---|---|---|---|---|---|---|---|---|---|---|---|---|
| | S | G | D | T | S | P | R | H | L | S | N | V | S | S | T | G | S | I | D | M | V | D | S | P | Q | L | A | T | L | A | D | E | V | S | A | S | L | A | K | Q | G | L | E | G | T | T | A | E | E | A | Sequence |
| Min | 400 | 401 | 402 | 403 | 404 | 405 | 406 | 407 | 408 | 409 | 410 | 411 | 412 | 413 | 414 | 415 | 416 | 417 | 418 | 419 | 420 | 421 | 422 | 423 | 424 | 425 | 426 | 427 | 428 | 429 | 430 | 431 | 432 | 433 | 434 | 435 | 436 | 437 | 438 | 439 | 440 | 441 | 99 | 100 | 101 | 102 | 103 | 104 | 105 | 106 | Residue no. wt tau |
| 0 | 0 | | | 0 | | | 0 | 0 | 0 | 0 | 0 | 0 | 0 | | 0 | 0 | | | | | 0 | | | | 0 | 0 | 0 | 0 | 0 | 0 | 0 | | 3 | | 0 | 0 | | 0 | | 0 | | 1 | | | 0 | | | | | 1 | Rel. freq. cuts |
| 0.25 | 0 | | | 0 | | | 0 | 1 | 4 | 1 | 0 | 3 | 0 | | 0 | 0 | | | | | 0 | | | | 0 | 0 | 0 | 0 | 3 | 0 | 0 | | 6 | | 0 | 0 | | 0 | | 0 | | 3 | | | 0 | | | | | 1 | |
| 1 | 0 | | | 0 | | | 1 | 2 | 8 | 1 | 1 | 3 | 1 | | 0 | 0 | | | | | 1 | | | | 0 | 1 | 1 | 1 | 5 | 0 | 0 | | 2 | | 0 | 0 | | 0 | | 1 | | 8 | | | 0 | | | | | 0 | |
| 10 | 1 | | | 1 | | | 1 | 1 | 20 | 0 | 0 | 9 | 0 | | 3 | 0 | | | | | 3 | | | | 1 | 2 | 7 | 2 | 10 | 3 | 1 | | 9 | | 1 | 2 | | 1 | | 3 | | 13 | | | 1 | | | | | 1 | |
| 2 | 0 | | | 1 | | | 1 | 2 | 15 | 1 | 1 | 8 | 1 | | 1 | 0 | | | | | 2 | | | | 1 | 2 | 3 | 1 | 7 | 1 | 0 | | 7 | | 0 | 0 | | 1 | | 1 | | 10 | | | 1 | | | | | 0 | |
| 5 | 1 | | | 0 | | | 1 | 2 | 23 | 1 | 1 | 13 | 1 | | 2 | 1 | | | | | 4 | | | | 1 | 2 | 7 | 2 | 9 | 3 | 1 | | 8 | | 1 | 2 | | 1 | | 2 | | 14 | | | 1 | | | | | 1 | |
| 120 | 1 | | | 1 | | | 4 | 2 | 25 | 1 | 1 | 14 | 1 | | 4 | 2 | | | | | 4 | | | | 0 | 1 | 8 | 2 | 14 | 2 | 0 | | 13 | | 0 | 2 | | 1 | | 1 | | 7 | | | 2 | | | | | 4 | |

| | 101 | 102 | 103 | 104 | 105 | 106 | 107 | 108 | 109 | 110 | 111 | 112 | 113 | 114 | 115 | 116 | 117 | 118 | 119 | 120 | 121 | 122 | 123 | 124 | 125 | 126 | 127 | 128 | 129 | 130 | 131 | 132 | 133 | 134 | 135 | 136 | 137 | 138 | 139 | 140 | 141 | 142 | 143 | 144 | 145 | 146 | 147 | 148 | 149 | 150 | Residue no. |
|---|---|---|---|---|---|---|---|---|---|---|---|---|---|---|---|---|---|---|---|---|---|---|---|---|---|---|---|---|---|---|---|---|---|---|---|---|---|---|---|---|---|---|---|---|---|---|---|---|---|---|---|
| | G | I | G | D | T | P | S | L | E | D | E | A | A | G | H | V | T | Q | A | R | M | V | S | K | S | K | D | G | T | G | S | D | D | K | K | A | K | G | A | D | G | K | T | K | I | A | T | P | R | G | Sequence |
| Min | 107 | 108 | 109 | 110 | 111 | 112 | 113 | 114 | 115 | 116 | 117 | 118 | 119 | 120 | 121 | 122 | 123 | 124 | 125 | 126 | 127 | 128 | 129 | 130 | 131 | 132 | 133 | 134 | 135 | 136 | 137 | 138 | 139 | 140 | 141 | 142 | 143 | 144 | 145 | 146 | 147 | 148 | 149 | 150 | 151 | 152 | 153 | 154 | 155 | 156 | Residue no. wt tau |
| 0 | | | | 0 | | | | | | | | | | | | | | | 1 | | | 0 | | | | | | | | | | | | | | | | | | | | | | | | | | | | | Rel. freq. cuts |
| 0.25 | | | | 0 | | | | | | | | | | | | | | | 2 | | | 0 | | | | | | | | | | | | | | | | | | | | | | | | | | | | | |
| 1 | | | | 0 | | | | | | | | | | | | | | | 1 | | | 0 | | | | | | | | | | | | | | | | | | | | | | | | | | | | | |
| 10 | | | | 0 | | | | | | | | | | | | | | | 0 | | | 0 | | | | | | | | | | | | | | | | | | | | | | | | | | | | | |
| 2 | | | | 0 | | | | | | | | | | | | | | | 1 | | | 0 | | | | | | | | | | | | | | | | | | | | | | | | | | | | | |
| 5 | | | | 0 | | | | | | | | | | | | | | | 0 | | | 0 | | | | | | | | | | | | | | | | | | | | | | | | | | | | | |
| 120 | | | | 1 | | | | | | | | | | | | | | | 2 | | | 1 | | | | | | | | | | | | | | | | | | | | | | | | | | | | | |

| | 151 | 152 | 153 | 154 | 155 | 156 | 157 | 158 | 159 | 160 | 161 | 162 | 163 | 164 | 165 | 166 | 167 | 168 | 169 | 170 | 171 | 172 | 173 | 174 | 175 | 176 | 177 | 178 | 179 | 180 | 181 | 182 | 183 | 184 | 185 | 186 | 187 | 188 | 189 | 190 | 191 | 192 | 193 | 194 | 195 | 196 | 197 | 198 | 199 | 200 | Residue no. |
|---|---|---|---|---|---|---|---|---|---|---|---|---|---|---|---|---|---|---|---|---|---|---|---|---|---|---|---|---|---|---|---|---|---|---|---|---|---|---|---|---|---|---|---|---|---|---|---|---|---|---|---|
| | A | A | P | P | G | Q | K | G | Q | A | N | A | T | R | I | P | A | K | T | P | P | A | P | K | T | P | P | S | S | G | E | P | P | K | S | G | D | R | S | G | Y | S | S | P | G | S | P | G | T | P | Sequence |
| Min | 157 | 158 | 159 | 160 | 161 | 162 | 163 | 164 | 165 | 166 | 167 | 168 | 169 | 170 | 171 | 172 | 173 | 174 | 175 | 176 | 177 | 178 | 179 | 180 | 181 | 182 | 183 | 184 | 185 | 186 | 187 | 188 | 189 | 190 | 191 | 192 | 193 | 194 | 195 | 196 | 197 | 198 | 199 | 200 | 201 | 202 | 203 | 204 | 205 | 206 | Residue no. wt tau |
| 0 | | | | | | | | | | | | | | | | | | | | | | | | | | | | | | | | | | | | | | | | | | | | | | | | | | | Rel. freq. cuts |
| 0.25 | | | | | | | | | | | | | | | | | | | | | | | | | | | | | | | | | | | | | | | | | | | | | | | | | | | |
| 1 | | | | | | | | | | | | | | | | | | | | | | | | | | | | | | | | | | | | | | | | | | | | | | | | | | | |
| 2 | | | | | | | | | | | | | | | | | | | | | | | | | | | | | | | | | | | | | | | | | | | | | | | | | | | |
| 5 | | | | | | | | | | | | | | | | | | | | | | | | | | | | | | | | | | | | | | | | | | | | | | | | | | | |
| 10 | | | | | | | | | | | | | | | | | | | | | | | | | | | | | | | | | | | | | | | | | | | | | | | | | | | |
| 120 | | | | | | | | | | | | | | | | | | | | | | | | | | | | | | | | | | | | | | | | | | | | | | | | | | | |

| | 201 | 202 | 203 | 204 | 205 | 206 | 207 | 208 | 209 | 210 | 211 | 212 | 213 | 214 | 215 | 216 | 217 | 218 | 219 | 220 | 221 | 222 | 223 | 224 | 225 | 226 | 227 | 228 | 229 | 230 | 231 | 232 | 233 | 234 | 235 | 236 | 237 | 238 | 239 | 240 | 241 | 242 | 243 | 244 | 245 | 246 | 247 | 248 | 249 | 250 | Residue no. |
|---|---|---|---|---|---|---|---|---|---|---|---|---|---|---|---|---|---|---|---|---|---|---|---|---|---|---|---|---|---|---|---|---|---|---|---|---|---|---|---|---|---|---|---|---|---|---|---|---|---|---|---|
| | G | S | R | S | R | T | P | S | L | P | T | P | P | T | R | E | P | K | K | V | A | V | V | R | T | P | P | K | S | P | S | S | A | K | S | R | L | Q | T | A | P | V | P | M | P | D | L | K | N | V | Sequence |
| Min | 207 | 208 | 209 | 210 | 211 | 212 | 213 | 214 | 215 | 216 | 217 | 218 | 219 | 220 | 221 | 222 | 223 | 224 | 225 | 226 | 227 | 228 | 229 | 230 | 231 | 232 | 233 | 234 | 235 | 236 | 237 | 238 | 239 | 240 | 241 | 242 | 243 | 244 | 245 | 246 | 247 | 248 | 249 | 250 | 251 | 252 | 253 | 254 | 255 | 256 | Residue no. wt tau |
| 0 | | | | | | | | | | | | | | | | | | | | 0 | 0 | 0 | 0 | 0 | | 0 | | 0 | | 0 | 0 | | 0 | 0 | 0 | 1 | 0 | 0 | 0 | 0 | 0 | 0 | | | | | 0 | | | 0 | Rel. freq. cuts |
| 0.25 | | | | | | | | | | | | | | | | | | | | 1 | 0 | 0 | 0 | 0 | | 0 | | 0 | | 0 | 1 | | 0 | 0 | 0 | 2 | 0 | 0 | 0 | 0 | 0 | 0 | | | | | 0 | | | 2 | |
| 1 | | | | | | | | | | | | | | | | | | | | 2 | 1 | 0 | 1 | 0 | | 0 | | 0 | | 0 | 1 | | 0 | 1 | 0 | 3 | 0 | 1 | 0 | 0 | 0 | 0 | | | | | 0 | | | 9 | |
| 2 | | | | | | | | | | | | | | | | | | | | 1 | 0 | 0 | 0 | 0 | | 0 | | 0 | | 0 | 1 | | 0 | 1 | 1 | 2 | 0 | 2 | 2 | 1 | 1 | 1 | | | | | 0 | | | 13 | |
| 5 | | | | | | | | | | | | | | | | | | | | 3 | 1 | 0 | 0 | 0 | | 0 | | 0 | | 0 | 2 | | 0 | 1 | 1 | 4 | 0 | 2 | 1 | 0 | 0 | 0 | | | | | 0 | | | 10 | |
| 10 | | | | | | | | | | | | | | | | | | | | 6 | 1 | 1 | 2 | 0 | | 0 | | 0 | | 0 | 3 | | 0 | 1 | 2 | 4 | 0 | 4 | 2 | 1 | 0 | 0 | | | | | 0 | | | 20 | |
| 120 | | | | | | | | | | | | | | | | | | | | 13 | 2 | 2 | 6 | 1 | | 1 | | 1 | | 1 | 6 | | 2 | 2 | 4 | 4 | 2 | 8 | 4 | 1 | 1 | 1 | | | | | 1 | | | 33 | |

| | 251 | 252 | 253 | 254 | 255 | 256 | 257 | 258 | 259 | 260 | 261 | 262 | 263 | 264 | 265 | 266 | 267 | 268 | 269 | 270 | 271 | 272 | 273 | 274 | 275 | 276 | 277 | 278 | 279 | 280 | 281 | 282 | 283 | 284 | 285 | 286 | 287 | 288 | 289 | 290 | 291 | 292 | 293 | 294 | 295 | 296 | 297 | 298 | 299 | 300 | Residue no. |
|---|---|---|---|---|---|---|---|---|---|---|---|---|---|---|---|---|---|---|---|---|---|---|---|---|---|---|---|---|---|---|---|---|---|---|---|---|---|---|---|---|---|---|---|---|---|---|---|---|---|---|---|
| | K | S | K | I | G | S | T | E | N | L | K | H | Q | P | G | G | G | K | V | Q | I | I | N | K | K | L | D | L | S | N | V | Q | S | K | C | G | S | K | D | N | I | K | H | V | P | G | G | G | S | V | Sequence |
| | 257 | 258 | 259 | 260 | 261 | 262 | 263 | 264 | 265 | 266 | 267 | 268 | 269 | 270 | 271 | 272 | 273 | 274 | 275 | 276 | 277 | 278 | 279 | 280 | 281 | 282 | 283 | 284 | 285 | 286 | 287 | 288 | 289 | 290 | 291 | 292 | 293 | 294 | 295 | 296 | 297 | 298 | 299 | 300 | 301 | 302 | 303 | 304 | 305 | 306 | Residue no. wt tau |
| Min | | | | | | | | | | | | | | | | G | G | K | V | Q | I | I | N | K | K | L | D | L | S | N | V | Q | S | K | C | G | S | K | D | N | I | K | H | V | P | G | G | G | S | V | B-factors |
| 0 | 0 | | | | 1 | | 0 | | | 0 | 0 | | | | | | | | 0 | 0 | 0 | 0 | 0 | 0 | | | | 0 | | | 0 | | 0 | | 0 | 0 | | | | | 0 | | | | | | | | | | Rel. freq. cuts |
| 0.25 | 0 | | | | 2 | | 0 | | | 0 | 0 | | | | | | | | 0 | 1 | 0 | 0 | 0 | 0 | | | | 0 | | | 1 | | 0 | | 1 | 0 | | | | | 0 | | | | | | | | | | |
| 1 | 1 | | | | 2 | | 0 | | | 4 | 0 | | | | | | | | 0 | 1 | 2 | 1 | 0 | 0 | | | | 0 | | | 3 | | 0 | | 1 | 1 | | | | | 0 | | | | | | | | | | |
| 2 | 1 | | | | 0 | | 0 | | | 3 | 0 | | | | | | | | 0 | 1 | 2 | 0 | 0 | 1 | | | | 0 | | | 3 | | 0 | | 0 | 0 | | | | | 1 | | | | | | | | | | |
| 5 | 1 | | | | 2 | | 1 | | | 5 | 0 | | | | | | | | 0 | 1 | 3 | 1 | 0 | 0 | | | | 0 | | | 4 | | 0 | | 1 | 2 | | | | | 1 | | | | | | | | | | |
| 10 | 1 | | | | 2 | | 0 | | | 8 | 0 | | | | | | | | 0 | 3 | 4 | 1 | 1 | 1 | | | | 0 | | | 8 | | 1 | | 1 | 2 | | | | | 2 | | | | | | | | | | |
| 120 | 2 | | | | 0 | | 1 | | | 12 | 1 | | | | | | | | 1 | 3 | 6 | 2 | 1 | 3 | | | | 1 | | | 10 | | 2 | | 1 | 2 | | | | | 2 | | | | | | | | | | |

| | 301 | 302 | 303 | 304 | 305 | 306 | 307 | 308 | 309 | 310 | 311 | 312 | 313 | 314 | 315 | 316 | 317 | 318 | 319 | 320 | 321 | 322 | 323 | 324 | 325 | 326 | 327 | 328 | 329 | 330 | 331 | 332 | 333 | 334 | 335 | 336 | 337 | 338 | 339 | 340 | 341 | 342 | 343 | 344 | 345 | 346 | 347 | 348 | 349 | 350 | Residue no. |
|---|---|---|---|---|---|---|---|---|---|---|---|---|---|---|---|---|---|---|---|---|---|---|---|---|---|---|---|---|---|---|---|---|---|---|---|---|---|---|---|---|---|---|---|---|---|---|---|---|---|---|---|
| | Q | I | V | Y | K | P | V | D | L | S | K | V | T | S | K | C | G | S | L | G | N | I | H | H | K | P | G | G | G | Q | V | E | V | K | S | E | K | L | D | F | K | D | R | V | Q | S | K | I | G | S | Sequence |
| | 307 | 308 | 309 | 310 | 311 | 312 | 313 | 314 | 315 | 316 | 317 | 318 | 319 | 320 | 321 | 322 | 323 | 324 | 325 | 326 | 327 | 328 | 329 | 330 | 331 | 332 | 333 | 334 | 335 | 336 | 337 | 338 | 339 | 340 | 341 | 342 | 343 | 344 | 345 | 346 | 347 | 348 | 349 | 350 | 351 | 352 | 353 | 354 | 355 | 356 | Residue no. wt tau |
| Min | Q | I | V | Y | K | P | V | D | L | S | K | V | T | S | K | C | G | S | L | G | N | I | H | H | | | | | | | | | | | | | | | | | | | | | | | | | | | B-factors |
| 0 | 0 | 0 | 0 | 0 | | | | | 0 | 0 | | 0 | 0 | 0 | 0 | 0 | 0 | 0 | 0 | | | | | | | | | | | 0 | | 0 | 0 | | | | 0 | 0 | | | | | 0 | 0 | 0 | 0 | 0 | | 0 | | Rel. freq. cuts |
| 0.25 | 1 | 3 | 0 | 0 | | | | | 0 | 0 | | 2 | 0 | 1 | 0 | 0 | 0 | 0 | 0 | | | | | | | | | | | 0 | | 0 | 0 | | | | 0 | 1 | | | | | 0 | 0 | 0 | 0 | 0 | | 0 | | |
| 1 | 1 | 7 | 0 | 0 | | | | | 1 | 0 | | 3 | 1 | 1 | 1 | 2 | 0 | 0 | 0 | | | | | | | | | | | 1 | | 0 | 1 | | | | 0 | 2 | | | | | 0 | 1 | 0 | 0 | 0 | | 0 | | |
| 2 | 1 | 5 | 0 | 0 | | | | | 1 | 1 | | 2 | 0 | 1 | 0 | 0 | 0 | 0 | 0 | | | | | | | | | | | 0 | | 0 | 0 | | | | 0 | 1 | | | | | 0 | 2 | 1 | 0 | 0 | | 0 | | |
| 5 | 2 | 10 | 1 | 0 | | | | | 1 | 0 | | 4 | 1 | 1 | 2 | 3 | 0 | 0 | 0 | | | | | | | | | | | 0 | | 0 | 2 | | | | 0 | 5 | | | | | 0 | 1 | 0 | 1 | 0 | | 0 | | |
| 10 | 3 | 14 | 2 | 0 | | | | | 1 | 1 | | 5 | 1 | 2 | 2 | 6 | 1 | 0 | 0 | | | | | | | | | | | 1 | | 1 | 2 | | | | 0 | 9 | | | | | 0 | 3 | 1 | 1 | 1 | | 1 | | |
| 120 | 2 | 15 | 3 | 1 | | | | | 2 | 0 | | 7 | 1 | 1 | 2 | 6 | 0 | 1 | 1 | | | | | | | | | | | 1 | | 0 | 2 | | | | 1 | 14 | | | | | 1 | 4 | 3 | 1 | 3 | | 0 | | |

| | 351 | 352 | 353 | 354 | 355 | 356 | 357 | 358 | 359 | 360 | 361 | 362 | 363 | 364 | 365 | 366 | 367 | 368 | 369 | 370 | 371 | 372 | 373 | 374 | 375 | 376 | 377 | 378 | 379 | 380 | 381 | 382 | 383 | 384 | 385 | 386 | 387 | 388 | 389 | 390 | 391 | 392 | 393 | 394 | 395 | 396 | 397 | 398 | 399 | 400 | Residue no. |
|---|---|---|---|---|---|---|---|---|---|---|---|---|---|---|---|---|---|---|---|---|---|---|---|---|---|---|---|---|---|---|---|---|---|---|---|---|---|---|---|---|---|---|---|---|---|---|---|---|---|---|---|
| | L | D | N | I | T | H | V | P | G | G | G | N | K | K | I | E | T | H | K | L | T | F | R | E | N | A | K | A | K | T | D | H | G | A | G | L | K | E | S | P | L | Q | T | P | T | E | D | G | S | E | Sequence |
| Min | 357 | 358 | 359 | 360 | 361 | 362 | 363 | 364 | 365 | 366 | 367 | 368 | 369 | 370 | 371 | 372 | 373 | 374 | 375 | 376 | 377 | 378 | 379 | 380 | 381 | 382 | 383 | 384 | 385 | 386 | 387 | 388 | 389 | 390 | 42 | 43 | 44 | 45 | 46 | 47 | 48 | 49 | 50 | 51 | 52 | 53 | 54 | 55 | 56 | 57 | Residue no. wt tau |
| 0 | 0 | 0 | | 0 | 0 | 0 | 0 | 0 | 0 | 0 | | | | | | 0 | 0 | 0 | 0 | 0 | 0 | | | | | | | | | | 0 | | | 0 | | | 0 | | | | | | | | | | | | | | Rel. freq. cuts |
| 0.25 | 0 | 0 | | 2 | 0 | 0 | 0 | 0 | 0 | 0 | | | | | | 0 | 0 | 1 | 0 | 1 | 0 | | | | | | | | | | 1 | | | 0 | | | 0 | | | | | | | | | | | | | | |
| 1 | 2 | 0 | | 8 | 0 | 1 | 1 | 1 | 1 | 1 | | | | | | 2 | 0 | 2 | 1 | 8 | 1 | | | | | | | | | | 1 | | | 1 | | | 0 | | | | | | | | | | | | | | |
| 2 | 0 | 0 | | 4 | 0 | 0 | 1 | 1 | 1 | 0 | | | | | | 0 | 0 | 0 | 0 | 3 | 0 | | | | | | | | | | 0 | | | 0 | | | 0 | | | | | | | | | | | | | | |
| 5 | 2 | 0 | | 12 | 0 | 2 | 2 | 2 | 2 | 1 | | | | | | 1 | 1 | 2 | 2 | 13 | 2 | | | | | | | | | | 1 | | | 1 | | | 0 | | | | | | | | | | | | | | |
| 10 | 4 | 0 | | 15 | 1 | 2 | 3 | 3 | 3 | 3 | | | | | | 5 | 1 | 3 | 3 | 20 | 3 | | | | | | | | | | 1 | | | 1 | | | 1 | | | | | | | | | | | | | | |
| 120 | 5 | 1 | | 17 | 2 | 2 | 5 | 3 | 3 | 3 | | | | | | 5 | 1 | 3 | 2 | 20 | 2 | | | | | | | | | | 1 | | | 1 | | | 1 | | | | | | | | | | | | | | |

| | 401 | 402 | 403 | 404 | 405 | 406 | 407 | 408 | 409 | 410 | 411 | 412 | 413 | 414 | 415 | 416 | 417 | 418 | 419 | 420 | 421 | 422 | 423 | 424 | 425 | 426 | 427 | 428 | 429 | 430 | 431 | 432 | 433 | 434 | 435 | 436 | 437 | 438 | 439 | 440 | 441 | | | | | | | | | | Residue no. |
|---|---|---|---|---|---|---|---|---|---|---|---|---|---|---|---|---|---|---|---|---|---|---|---|---|---|---|---|---|---|---|---|---|---|---|---|---|---|---|---|---|---|---|---|---|---|---|---|---|---|---|---|
| | E | P | G | S | E | T | S | D | A | K | S | T | P | T | A | E | D | V | T | A | P | L | V | D | E | G | A | P | G | K | Q | A | A | A | Q | P | H | T | E | I | P | | | | | | | | | | Sequence |
| Min | 58 | 59 | 60 | 61 | 62 | 63 | 64 | 65 | 66 | 67 | 68 | 69 | 70 | 71 | 72 | 73 | 74 | 75 | 76 | 77 | 78 | 79 | 80 | 81 | 82 | 83 | 84 | 85 | 86 | 87 | 88 | 89 | 90 | 91 | 92 | 93 | 94 | 95 | 96 | 97 | 98 | | | | | | | | | | Residue no. wt tau |
| 0 | | | | | 0 | | | | | | | | | | | | | | | | | | | | | | | | | | 0 | | | | | | | | | | | | | | | | | | | | Rel. freq. cuts |
| 0.25 | | | | | 0 | | | | | | | | | | | | | | | | | | | | | | | | | | 0 | | | | | | | | | | | | | | | | | | | | |
| 1 | | | | | 0 | | | | | | | | | | | | | | | | | | | | | | | | | | 0 | | | | | | | | | | | | | | | | | | | | |
| 2 | | | | | 0 | | | | | | | | | | | | | | | | | | | | | | | | | | 0 | | | | | | | | | | | | | | | | | | | | |
| 5 | | | | | 0 | | | | | | | | | | | | | | | | | | | | | | | | | | 0 | | | | | | | | | | | | | | | | | | | | |
| 10 | | | | | 1 | | | | | | | | | | | | | | | | | | | | | | | | | | 0 | | | | | | | | | | | | | | | | | | | | |
| 120 | | | | | 0 | | | | | | | | | | | | | | | | | | | | | | | | | | 1 | | | | | | | | | | | | | | | | | | | | |

D

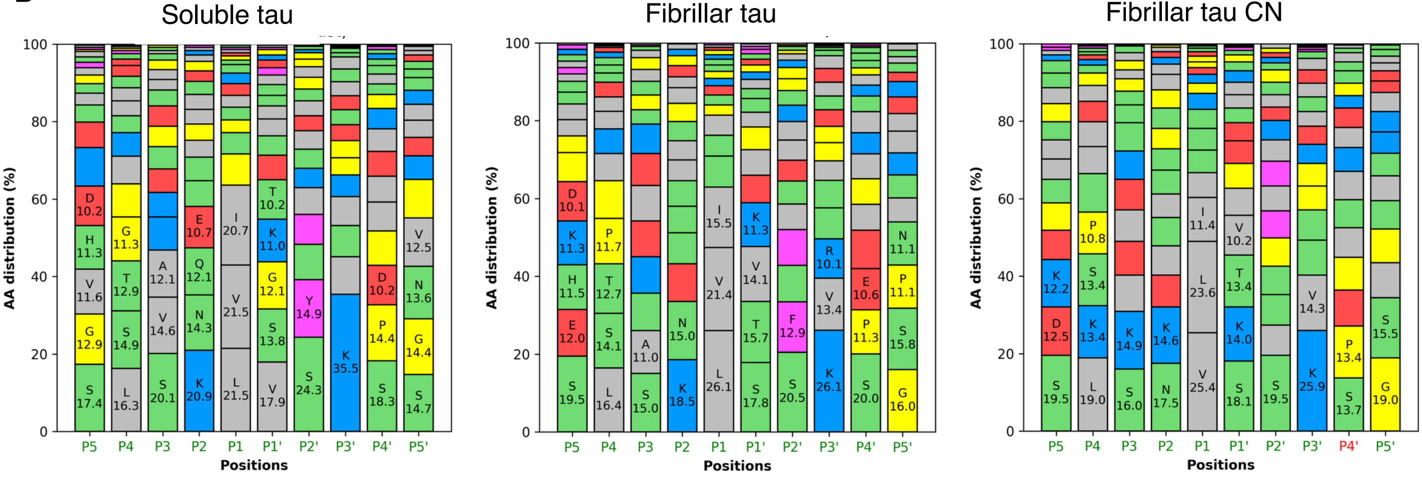


E

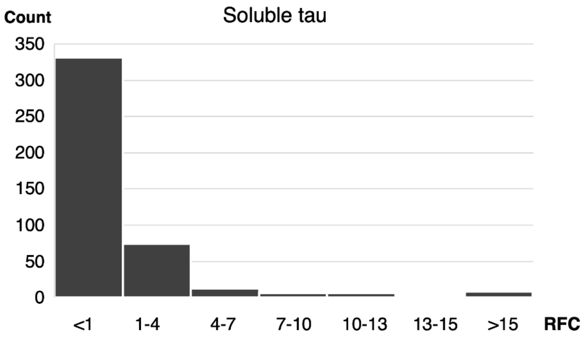


F

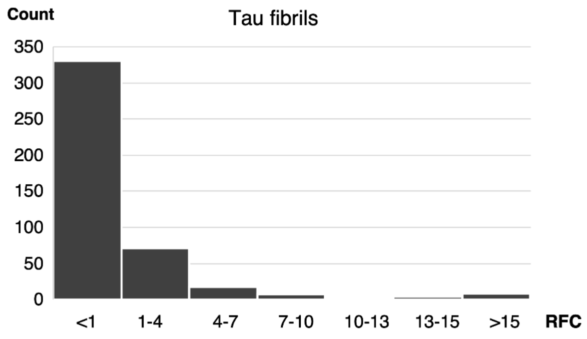


G

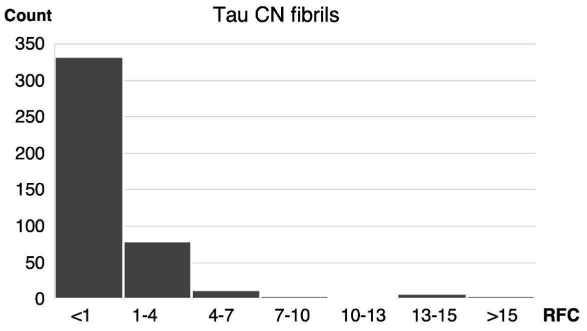


**Fig. S2. Proteolysis of soluble, fibrillar and fibrillar tau CN by HTRA1. Related to Fig. 2**

Proteolysis of soluble, fibrillar and fibrillar tau CN by HTRA1 was done as described in Fig. 2A. Samples were taken at the timepoints indicated (Min) and proteolytic products of tau were identified by LC-MS. Structural elements i.e. helices and loops (2ndary struct.), B-factors as detected in the cryo EM structure of fibrillar tau (pdb:6QJH) are indicated. Amino acid sequence conservation (Conservation) is derived from a multiple sequence alignment (Supplemental data 2); * = identical, : = conserved residues. P1 residues identified in n=4 experiments as significantly enriched compared to controls are represented by numbers indicating the relative frequency of cuts (Rel. freq. cuts) at each time point. No number indicates that no cleavage was detected at any of the time points investigated.

(A) Proteolysis of soluble tau by HTRA1.

(B) Proteolysis of fibrillar tau by HTRA1.

(C) Proteolysis of fibrillar tau CN by HTRA1. In fibrillar tau CN, parts of the N- and C-termini have been switched i.e. the C-terminal residues 391-441 were swapped with the N-terminal residues 42-98 (highlighted in grey).

(D) Relative amino acid distribution at P1-P5 and P1'-P5' sites of tau variants used in this study. X-axis: red letter, no protease selectivity toward specific amino acids at this position; green letter, protease selectivity toward certain amino acids at this position. Amino acid residue color code: Hydrophobic, grey; hydrophilic, green; positively charged, red; negatively charged, blue; Gly and Pro, yellow.

(E-G) Histograms of relative cleavage efficiencies. Related to Fig. 2 and supplemental text. To classify cleavage sites, the relative cleavage efficiencies after each residue was calculated by UMSAP using data of the 120 min timepoints. The histograms were generated by Xcel using width of interval: 3, underflow bin: 0.99; overflow bin: 20 as parameters. RFC, relative frequency of cuts.

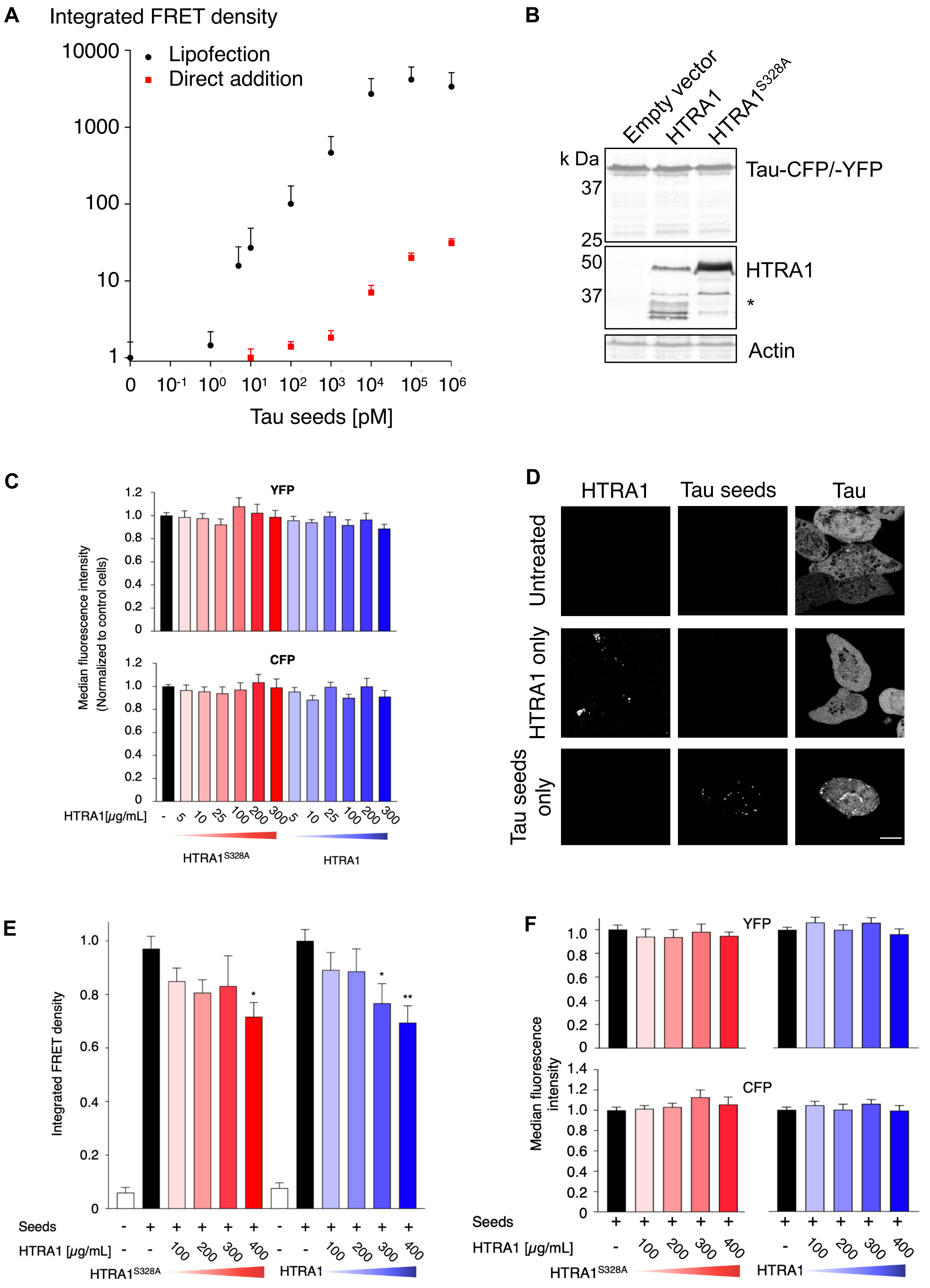

A
Integrated FRET density
Lipofection
Direct addition
Tau seeds [pM]
B
Empty vector
HTRA1
HTRA1S328A
k Da
Tau-CFP/-YFP
HTRA1
Actin
C
YFP
CFP
Median fluorescence intensity (Normalized to control cells)
HTRA1[μg/mL]
HTRA1S328A
HTRA1
D
HTRA1
Tau seeds
Tau
Untreated
HTRA1 only
Tau seeds only
E
Integrated FRET density
Seeds
HTRA1 [μg/mL]
HTRA1S328A
HTRA1
F
YFP
CFP
Median fluorescence intensity
Seeds
HTRA1 [μg/mL]
HTRA1S328A
HTRA1

**Fig. S3. FRET flow cytometry-based quantification of tau aggregation in cells. Related to Fig. 3.**
(A) Biosensor cells were treated with increasing amounts of tau seeds that were either lipotransfected or added directly. After 24 h, intracellular tau aggregation was investigated by FRET flow cytometry. Error bars, s.e.m.; $n \geq 2$.

(B) To test the stability of hybrid proteins, biosensor cells expressing the microtubule binding domain (MTBD) of tau tagged to CFP or YFP, respectively, were transfected with either empty vector control or plasmids overexpressing myc-tagged HTRA1 or HTRA1$^{S328A}$. Cell extracts were subjected to SDS-PAGE and immunoblotted with anti-GFP, myc-tag and actin antibodies, respectively. Asterisk: HTRA1 degradation products.

(C) Biosensor cells were treated with recombinant HTRA1 or HTRA1$^{S328A}$ and tau seeds as described in Fig. 4A. Flow cytometry-based analysis of the median fluorescence intensity of YFP and CFP revealed no significant differences between samples (one-way-ANOVA with Dunnett's posttest). Values are normalized to biosensor cells treated only with seeds (highlighted in black). Error bars, s.e.m.; $n = 4$.

(D) Confocal microscopy analysis of biosensor cells treated with fluorescently labeled HTRA1 and tau seeds. Biosensor cells were untreated, incubated with 25 µg/mL DyLight 633-labeled HTRA1$^{S328A}$ or with 300 nM Alexa Fluor 568-labeled tau seeds. Analysis was performed by confocal microscopy. Scale bar: 10 µm.

(E) HTRA1-mediated decrease of intracellular tau aggregates. Biosensor cells were treated with 25 nM tau seeds for 24 h followed by the addition of recombinant HTRA1$^{S328A}$ or HTRA1 at the concentrations indicated. As a control, buffer alone was added. After 24 h, cells were subjected to flow cytometry to quantify the integrated FRET density and the median fluorescence intensity of YFP and CFP. The signal of biosensor cells treated only with seeds (highlighted in black) was used for the normalization of values and the analysis of significant differences. Error bars, s.e.m.; $n \geq 3$. *$P < 0.05$, ** $P < 0.01$, one-way-ANOVA with Dunnett's posttest.

(F) The median fluorescence intensity of YFP and CFP was not significantly different between samples.

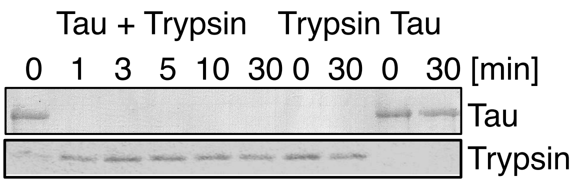


**Fig. S4. Trypsin digestion of tau seeds. Related to Fig. 4.** Seeds (Tau) were incubated with trypsin at 37°C. Volumes and concentrations were identical to the cell culture assay (A). Samples were collected at time points indicated and subjected to SDS-PAGE, Coomassie staining and immunoblotting against tau. Tau seeds incubated in buffer and trypsin alone served as controls.

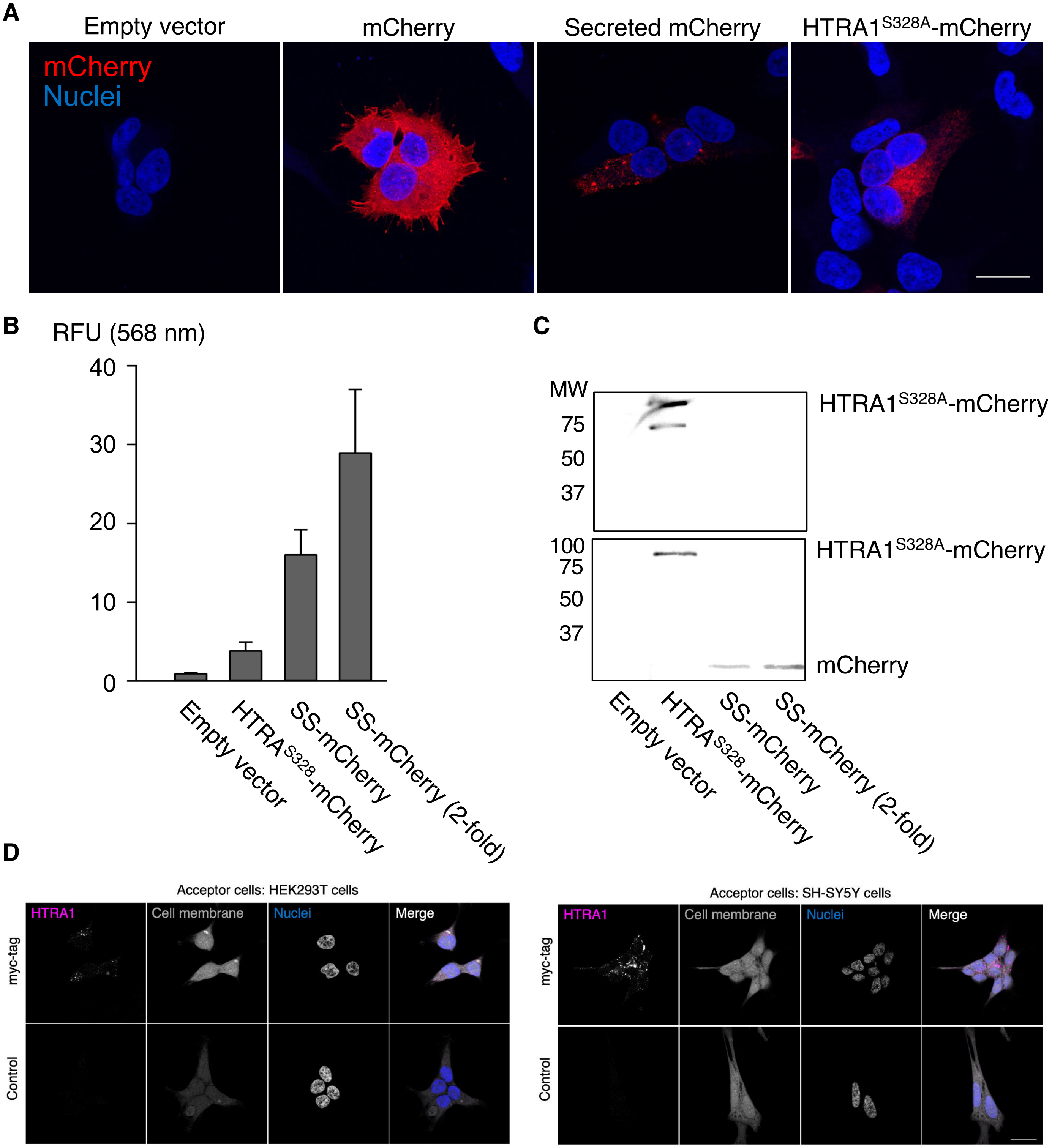


**Fig. S5. Analysis of the secretion and fluorescence of secreted HTRA1-mcherry or mCherry alone. Related to Fig. 5.**

(A) Confocal microscopy analysis of HEK293T donor cells overexpressing mCherry, secreted mCherry or HTRA1$^{S328A}$-mCherry. As a control, cells were transfected with the empty vector. Scale bar: 10 µm.

(B) Quantification of the fluorescence intensity at 568 nm in the cell culture supernatant. Error bars, s.e.m.; n = 3.

(C) HEK293T cells were transiently transfected with HTRA1$^{S328A}$-mCherry, mCherry containing the signal peptide of HTRA1 (SS-mCherry), or the corresponding empty vector. After 4 days, the medium was centrifuged to remove cells and the supernatant was subjected to SDS-PAGE and immunoblotting against HTRA1 (upper panel) and mCherry (lower panel). The medium containing secreted mCherry was concentrated to adjust the protein amount to HTRA1$^{S328A}$-mCherry. MW, molecular weight [kDa]. 2-fold indicates a two-fold concentration of the medium.

(D) The cellular location of internalized HTRA1 is tag-independent. HEK293T and SH-SY5Y acceptor cells were co-cultured with HEK293T cells overexpressing myc-tagged HTRA1$^{S328A}$. After 3 days, recipient cells were fixed and immunostained against the myc-tag. Cell nuclei and membranes were counterstained. As a control, the primary antibody was omitted. Analysis was performed by confocal microscopy. Scale bar: 10 µm.

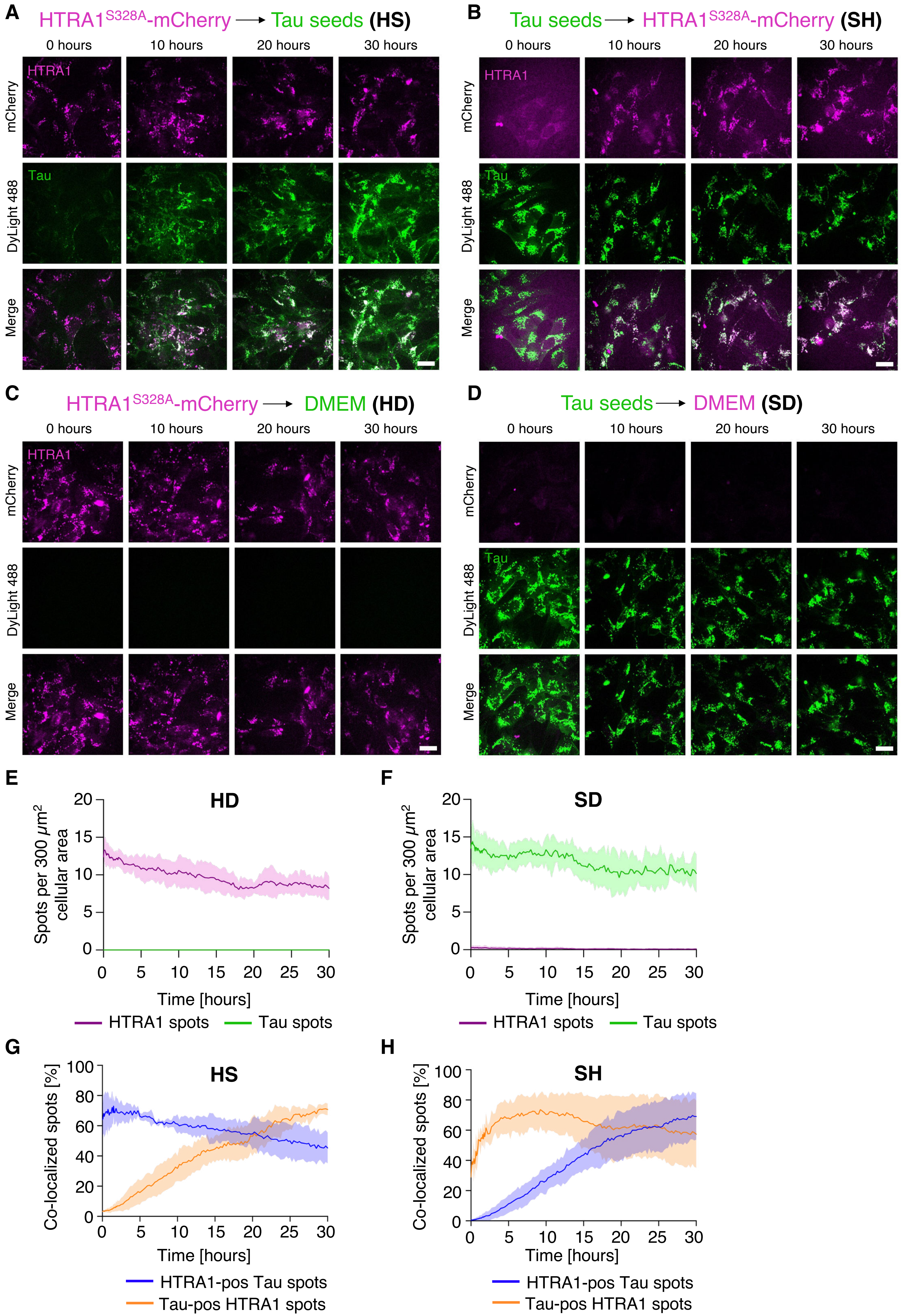

A
HTRA1S328A-mCherry → Tau seeds (HS)
0 hours
10 hours
20 hours
30 hours
mCherry
HTRA1
DyLight 488
Tau
Merge
B
Tau seeds → HTRA1S328A-mCherry (SH)
C
HTRA1S328A-mCherry → DMEM (HD)
D
Tau seeds → DMEM (SD)
E
HD
Spots per 300 μm2 cellular area
Time [hours]
HTRA1 spots
Tau spots
F
SD
G
HS
Co-localized spots [%]
HTRA1-pos Tau spots
Tau-pos HTRA1 spots
H
SH

**Fig. S6. Live cell analysis of SH-SY5Y cells treated with secreted HTRA1$^{S328A}$-mCherry and/or fluorescently labeled tau seeds. Related to Fig. 6. (**A-D) Live cell imaging of SH-SY5Y cells. Scale bars: 20 µm.

(A) HS treatment: Cells were treated with conditioned medium containing secreted HTRA1S328A-mCherry (magenta) 24 hours before imaging and DyLight 488-labeled tau seeds (green) immediately prior to imaging.

(B) SH treatment: Cells were treated with DyLight 488-labeled tau seeds (green) 24 hours before imaging and conditioned medium containing secreted HTRA1S328A-mCherry (magenta) immediately prior to imaging.

(C) HD treatment: Cells were treated with conditioned medium containing secreted HTRA1S328A-mCherry (magenta) 24 hours before imaging and DMEM immediately prior to imaging.

(D) SD treatment: Cells were treated with DyLight 488-labeled tau seeds (green) 24 hours before imaging and DMEM immediately prior to imaging.

(E–F) Time-resolved analysis of HTRA1S328A-mCherry and DyLight 488-labeled tau seed spots in SH-SY5Y control treatment cells (10 random positions per condition/experiment, n=3; in total, more than 10.500 individual HTRA1 and Tau spots were detected and subsequently analyzed). Cells were either treated first with HTRA1S328A-mCherry followed by DMEM (HD shown in E) or DyLight 488-labeled tau seeds followed by DMEM (SD shown in F). In each frame, the CellTracker Deep Red stain was used to detect the image area that was covered with cells (total cell area). Discrete fluorescent spots were detected in the mCherry and DyLight 488 channel in this region only. The number of detected discrete HTRA1S328A-mCherry (magenta line) and/or tau seed spots (green line) was normalized by the measured cell area (number of spots/µm2) and multiplied by the factor 300, which reflects the average cell area of an individual SH-SY5Y cell. Data show mean ± SD.

(G-H) Time-resolved co-localization analysis of HTRA1S328A-mCherry and DyLight 488-labeled tau seed spots (corresponding data from Fig. 6E). Discrete spots of HTRA1S328A-mCherry were detected and masked with DyLight 488-labeled tau seed spots to identify double-positive species (Tau-positive HTRA1 spots) or vice versa (HTRA1-positive Tau spots). Data show mean ± SD.

**Table S1. P1 residues in tau with the highest relative number of cuts**

Relative number of cuts at 120 min

| Residue | Sol tau | Fibrillar tau | Fibrillar tau CN |
|---|---|---|---|
| V256 | 22 | 40 | 33 |
| C291(β) | 24 | | |
| I308(β) | 42 | | |
| C322 | 32 | | |
| L376 | 27 | 59 | 20 |
| I392 | 51 | 49 | |
| L408 | 29 | 40 | 25 |
| V411 | | 25 | |

Class I P1 residues (> 20 cuts) detected in proteolysis experiments with various tau constructs and conformations at the 120 min timepoint. (β) indicates residues of the β-sheets of the fibrillar core.

**Table S2. P1 residues in tau with the second highest relative number of cuts**

Relative number of cuts at 120 min

| Residue | Sol tau | Fibrillar tau | Fibrillar tau CN |
|---|---|---|---|
| V226 | | 15 | 13 |
| L266 | 11 | 15 | 12 |
| I277(β) | 13 | | |
| I278(β) | 12 | 11 | |
| V287(β) | 13 | | 10 |
| S289(β) | 10 | | |
| I308(β) | >20 | 17 | 15 |
| I328 | 13 | | |
| L344 | 15 | 18 | 14 |
| I360 | | 17 | 17 |
| I392 | >20 | >20 | 15 |
| V398 | | 12 | |
| V411 | 16 | >20 | 14 |
| L428 | 13 | 15 | 14 |
| V432 | 12 | 14 | 13 |

Class II P1 residues (10 - 19 cuts) detected in proteolysis experiments with various tau constructs and conformations at the 120 min timepoint. (β) indicates residues of the β-sheets of the fibrillar core.

**Table S3. P1 residues that are differentially cleaved in soluble and fibrillar tau at the 120 min timepoint**

| | Relative number of cuts at 120 min | | |
|---|---|---|---|
| Residue | Sol tau | Fibrillar tau | Fibrillar tau CN |
| V275 | 6 | 2 | 1 |
| Q276 | - | 2 | 3 |
| I277 | 13 | 7 | 6 |
| I278(β) | 12 | 11 | 2 |
| N279(β) | - | - | 1 |
| K280(β) | 7 | - | 3 |
| L282(β) | 6 | - | - |
| L284(β) | 2 | - | 1 |
| S285(β) | 2 | 2 | - |
| V287(β) | 13 | 9 | 10 |
| S289(β) | 10 | 4 | 2 |
| C291(β) | 24 | 2 | 1 |
| V306(β) | 2 | - | - |
| Q307(β) | 1 | - | 2 |
| I308(β) | 42 | 17 | 15 |
| V309(β) | 9 | 2 | 3 |
| Y310(β) | 2 | - | 1 |
| V313(β) | 8 | - | - |
| K317(β) | 1 | 2 | - |
| V318(β) | 9 | 4 | 7 |
| N327(β) | 1 | - | - |
| I328(β) | 13 | 1 | - |

Relative number of cuts at P1 residues indicated. No number corresponds to no cleavages detected. (β) indicates residues of the β-sheets of the fibrillar core.

**Legends for Movies S1 to S2**

**Movie S1**

Live-cell timelapse confocal spinning disk microscopy of cells containing cytosolic tau seeds (green), which were treated with extracellular HTRA1 (red). Uptake and colocalization of HTRA1 with tau seeds was followed over 30 min. Colocalized particles appear white. See text for details.

**Movie S2**

Live-cell timelapse confocal spinning disk microscopy of cells containing cytosolic HTRA1 (red), which were treated with extracellular tau seeds (green). Uptake and colocalization of tau seeds with HTRA1 was followed over 30 min. Colocalized particles appear white. See text for details.

**Supplemental data 1**

**2N4R tau**

```
MAEPRQEFEVMEDHAGTYGLGDRKDQGGYTMHQDQEGDTDAGLKESPLQTPTEDGSEEPG
SETSDAKSTPTAEDVTAPLVDEGAPGKQAAAQPHTEIP EGTTAEEAGIGDTPSLEDEAAG
HVTQARMVSKSKDGTGSDDKKAKGADGKTKIATPRGAAPPGQKGQANATRIPAKTPPAPK
TPPSSGEPPKSGDRSGYSSPGSPGTPGSRSRTPSLPTPPTREPKKVAVVRTPPKSPSSAK
SRLQTAPVPMPDLKNVKSKIGSTENLKHQPGGGKVQIINKKLDLSNVQSKCGSKDNIKHV
PGGGSVQIVYKPVDLSKVTSKCGSLGNIHHKPGGGQVEVKSEKLDFKDRVQSKIGSLDNI
THVPGGGNKKIETHKLTFRENAKAKTDHGA EIVYKSPVVSGDTSPRHLSNVSSTGSIDMV
DSPQLATLADEVSASLAKQGL
```

**Soluble Tau 15 sec**

```
 1 MAEPRQEFEVMEDHAGTYGLGDRKDQGGYTMHQDQEGDTDAGLKESPLQTPTEDGSEEPG

 1 SETSDAKSTPTAEDVTAPLVDEGAPGKQAAAQPHTEIPEGTTAEEAGIGDTPSLEDEAAG

 1 HVTQARMVSKSKDGTGSDDKKAKGADGKTKIATPRGAAPPGQKGQANATRIPAKTPPAPK

 1 TPPSSGEPPKSGDRSGYSSPGSPGTPGSRSRTPSLPTPPTREPKKVAVVRTPPKSPSSAK
 2                                               AVVRTPPKSPSSAK

 1 SRLQTAPVPMPDLKNVKSKIGSTENLKHQPGGGKVQIINKKLDLSNVQSKCGSKDNIKHV
 2 SRLQTAPVPMPDLKNV
 3     TAPVPMPDLKNV
 4                 KSKIGSTENL
 5                                      INKKLDLSNVQSKC
 6                                       NKKLDLSNVQSKC
 7                                         KLDLSNVQSKC
 8                                           DLSNVQSKCGSKDNIKHV
 9                                                    GSKDNIKHV

 1 PGGGSVQIVYKPVDLSKVTSKCGSLGNIHHKPGGGQVEVKSEKLDFKDRVQSKIGSLDNI
 8 PGGGSVQI
 9 PGGGSVQI
10         VYKPVDLSKV
11         VYKPVDLSKVTSKC
12           KPVDLSKVTSKC
13                       GSLGNIHHKPGGGQVEVKSEKL
14                                             DFKDRVQSKIGSLDNI

 1 THVPGGGNKKIETHKLTFRENAKAKTDHGAEIVYKSPVVSGDTSPRHLSNVSSTGSIDMV
15 THVPGGGNKKIETHKL
16                 TFRENAKAKTDHGAEI
17                 TFRENAKAKTDHGAEIV
18                 TFRENAKAKTDHGAEIVY
19                 TFRENAKAKTDHGAEIVYK
20                                 VYKSPVVSG
21                                 VYKSPVVSGDTSPRHL
22                                 VYKSPVVSGDTSPRHLS
23                                 VYKSPVVSGDTSPRHLSN
24                                 VYKSPVVSGDTSPRHLSNV
25                                 VYKSPVVSGDTSPRHLSNVS
26                                 VYKSPVVSGDTSPRHLSNVSSTG
27                                 VYKSPVVSGDTSPRHLSNVSSTGSIDM
28                                 VYKSPVVSGDTSPRHLSNVSSTGSIDMV
29                                 VYKSPVVSGDTSPRHLSNVSSTGSIDMV
30                                 VYKSPVVSGDTSPRHLSNVSSTGSIDMV
31                                                 SNVSSTGSIDMV
32                                                 SNVSSTGSIDMV
33                                                 SNVSSTGSIDMV
34                                                 SNVSSTGSIDMV
35                                                 SNVSSTGSIDMV
36                                                 SNVSSTGSIDMV

 1 DSPQLATLADEVSASLAKQGL
28 D
29 DSPQL
30 DSPQLATL
31 DSPQL
32 DSPQLA
33 DSPQLATL
34 DSPQLATLADEV
35 DSPQLATLADEVSA
36 DSPQLATLADEVSASLAKQGL
```

| | |
|---|---|
| 37 | ATLADEVSASLAKQGL |
| 38 | TLADEVSASLAKQGL |
| 39 | LADEVSASLAKQGL |
| 40 | ADEVSASLAKQGL |
| 41 | VSASLAKQGL |
| 42 | SASLAKQGL |

**Soluble Tau 1 min**

```
 1 MAEPRQEFEVMEDHAGTYGLGDRKDQGGYTMHQDQEGDTDAGLKESPLQTPTEDGSEEPG

 1 SETSDAKSTPTAEDVTAPLVDEGAPGKQAAAQPHTEIPEGTTAEEAGIGDTPSLEDEAAG

 1 HVTQARMVSKSKDGTGSDDKKAKGADGKTKIATPRGAAPPGQKGQANATRIPAKTPPAPK

 1 TPPSSGEPPKSGDRSGYSSPGSPGTPGSRSRTPSLPTPPTREPKKVAVVRTPPKSPSSAK
 2                                               AVVRTPPKSPSSAK

 1 SRLQTAPVPMPDLKNVKSKIGSTENLKHQPGGGKVQIINKKLDLSNVQSKCGSKDNIKHV
 2 SRLQTAPVPMPDLKNV
 3     TAPVPMPDLKNV
 4                 KSKIGSTENL
 5                                      INKKLDLSNV
 6                                      INKKLDLSNVQSKC
 7                                       NKKLDLSNV
 8                                         KLDLSNVQSKC
 9                                           DLSNVQSKC
10                                           DLSNVQSKCGSKDNIKHV
11                                                QSKCGSKDNIKHV
12                                                    GSKDNIKHV

 1 PGGGSVQIVYKPVDLSKVTSKCGSLGNIHHKPGGGQVEVKSEKLDFKDRVQSKIGSLDNI
10 PGGGSVQI
11 PGGGSVQI
12 PGGGSVQI
13   GGSVQIVYKPVDLSKVTSKC
14         VYKPVDLS
15         VYKPVDLSKV
16         VYKPVDLSKVT
17         VYKPVDLSKVTSKC
18          YKPVDLSKVTSKC
19              DLSKVTSKC
20              DLSKVTSKCGSLGNI
21                   TSKCGSLGNI
22                       GSLGNIHHKPGGGQVEVKSEKL
23                                             DFKDRVQSKIGSLDNI
24                                                   QSKIGSLDNI

 1 THVPGGGNKKIETHKLTFRENAKAKTDHGAEIVYKSPVVSGDTSPRHLSNVSSTGSIDMV
25 THVPGGGNKKIETHKL
26                 TFRENAKAKTDHGAEI
27                 TFRENAKAKTDHGAEIV
28                 TFRENAKAKTDHGAEIVY
29                 TFRENAKAKTDHGAEIVYK
30                                 VYKSPVVSG
31                                 VYKSPVVSGD
32                                 VYKSPVVSGDT
33                                 VYKSPVVSGDTS
34                                 VYKSPVVSGDTSPR
35                                 VYKSPVVSGDTSPRH
36                                 VYKSPVVSGDTSPRHL
37                                 VYKSPVVSGDTSPRHLS
38                                 VYKSPVVSGDTSPRHLSN
39                                 VYKSPVVSGDTSPRHLSNV
40                                 VYKSPVVSGDTSPRHLSNVS
41                                 VYKSPVVSGDTSPRHLSNVSST
42                                 VYKSPVVSGDTSPRHLSNVSSTG
43                                 VYKSPVVSGDTSPRHLSNVSSTGSID
44                                 VYKSPVVSGDTSPRHLSNVSSTGSIDM
45                                 VYKSPVVSGDTSPRHLSNVSSTGSIDMV
46                                 VYKSPVVSGDTSPRHLSNVSSTGSIDMV
```

```
47                                 VYKSPVVSGDTSPRHLSNVSSTGSIDMV
48                                 VYKSPVVSGDTSPRHLSNVSSTGSIDMV
49                                 VYKSPVVSGDTSPRHLSNVSSTGSIDMV
50                                 VYKSPVVSGDTSPRHLSNVSSTGSIDMV
51                                                 SNVSSTGSIDM
52                                                 SNVSSTGSIDMV
53                                                 SNVSSTGSIDMV
54                                                 SNVSSTGSIDMV
55                                                 SNVSSTGSIDMV
56                                                 SNVSSTGSIDMV
57                                                 SNVSSTGSIDMV
58                                                 SNVSSTGSIDMV
59                                                 SNVSSTGSIDMV
60                                                 SNVSSTGSIDMV
61                                                 SNVSSTGSIDMV
62                                                 SNVSSTGSIDMV
63                                                    SSTGSIDMV
64                                                    SSTGSIDMV
65                                                    SSTGSIDMV

 1 DSPQLATLADEVSASLAKQGL
46 D
47 DSP
48 DSPQL
49 DSPQLA
50 DSPQLATL
53 D
54 DS
55 DSP
56 DSPQ
57 DSPQL
58 DSPQLA
59 DSPQLATL
60 DSPQLATLADEV
61 DSPQLATLADEVSA
62 DSPQLATLADEVSASLAKQGL
63 DSPQL
64 DSPQLATL
65 DSPQLATLADEV
66     LATLADEVSASLAKQGL
67      ATLADEVSASLAKQGL
68       TLADEVSASLAKQGL
69        LADEVSASLAKQGL
70         ADEVSASLAKQGL
71             SASLAKQGL
```

**Soluble Tau 2 min**

```
 1 MAEPRQEFEVMEDHAGTYGLGDRKDQGGYTMHQDQEGDTDAGLKESPLQTPTEDGSEEPG
 2  AEPRQEFEV

 1 SETSDAKSTPTAEDVTAPLVDEGAPGKQAAAQPHTEIPEGTTAEEAGIGDTPSLEDEAAG

 1 HVTQARMVSKSKDGTGSDDKKAKGADGKTKIATPRGAAPPGQKGQANATRIPAKTPPAPK

 1 TPPSSGEPPKSGDRSGYSSPGSPGTPGSRSRTPSLPTPPTREPKKVAVVRTPPKSPSSAK
 3                                               AVVRTPPKSPSSAK
 4                                               AVVRTPPKSPSSAK

 1 SRLQTAPVPMPDLKNVKSKIGSTENLKHQPGGGKVQIINKKLDLSNVQSKCGSKDNIKHV
 3 SRLQ
 4 SRLQTAPVPMPDLKNV
 5     TAPVPMPDLKNV
 6                 KSKIGSTENL
 7                                      INKKLDLSNV
 8                                      INKKLDLSNVQS
 9                                      INKKLDLSNVQSKC
10                                       NKKLDLSNV
11                                       NKKLDLSNVQSKC
12                                         KLDLSNVQS
13                                         KLDLSNVQSKC
14                                           DLSNVQSKC
15                                           DLSNVQSKCGSKDNIKHV
16                                                QSKCGSKDNIKHV
17                                                  KCGSKDNIKHV
18                                                    GSKDNIKHV
19                                                    GSKDNIKHV

 1 PGGGSVQIVYKPVDLSKVTSKCGSLGNIHHKPGGGQVEVKSEKLDFKDRVQSKIGSLDNI
15 PGGGSVQI
16 PGGGSVQI
17 PGGGSVQI
18 PGGGSVQI
19 PGGGSVQIVYKPVDLSKVTSKC
20   GGSVQIVYKPVDLSKVTSKC
21         VYKPVDLS
22         VYKPVDLSKV
23         VYKPVDLSKVT
24         VYKPVDLSKVTSK
25         VYKPVDLSKVTSKC
26         VYKPVDLSKVTSKCGSLGNI
27          YKPVDLSKV
28          YKPVDLSKVTSKC
29              DLSKVTSKC
30              DLSKVTSKCGSLGNI
31                       GSLGNIHHKPGGGQVEV
32                       GSLGNIHHKPGGGQVEVKSEKL
33                          GNIHHKPGGGQVEVKSEKL
34                             HHKPGGGQVEVKSEKL
35                                             DFKDRVQSKIGSLDNI

 1 THVPGGGNKKIETHKLTFRENAKAKTDHGAEIVYKSPVVSGDTSPRHLSNVSSTGSIDMV
36 THVPGGGNKKIETHKL
37      GGNKKIETHKL
38                 TFRENAKAKTD
39                 TFRENAKAKTDHGAEI
40                 TFRENAKAKTDHGAEIV
41                 TFRENAKAKTDHGAEIVY
42                                 VYKSPVVSG
43                                 VYKSPVVSGD
```

```
44                                 VYKSPVVSGDT
45                                 VYKSPVVSGDTS
46                                 VYKSPVVSGDTSP
47                                 VYKSPVVSGDTSPR
48                                 VYKSPVVSGDTSPRH
49                                 VYKSPVVSGDTSPRHL
50                                 VYKSPVVSGDTSPRHLS
51                                 VYKSPVVSGDTSPRHLSN
52                                 VYKSPVVSGDTSPRHLSNV
53                                 VYKSPVVSGDTSPRHLSNVS
54                                 VYKSPVVSGDTSPRHLSNVSS
55                                 VYKSPVVSGDTSPRHLSNVSST
56                                 VYKSPVVSGDTSPRHLSNVSSTG
57                                 VYKSPVVSGDTSPRHLSNVSSTGS
58                                 VYKSPVVSGDTSPRHLSNVSSTGSID
59                                 VYKSPVVSGDTSPRHLSNVSSTGSIDM
60                                 VYKSPVVSGDTSPRHLSNVSSTGSIDMV
61                                 VYKSPVVSGDTSPRHLSNVSSTGSIDMV
62                                 VYKSPVVSGDTSPRHLSNVSSTGSIDMV
63                                 VYKSPVVSGDTSPRHLSNVSSTGSIDMV
64                                 VYKSPVVSGDTSPRHLSNVSSTGSIDMV
65                                 VYKSPVVSGDTSPRHLSNVSSTGSIDMV
66                                  YKSPVVSGDTSPRHL
67                                       VSGDTSPRHL
68                                                 SNVSSTGSIDM
69                                                 SNVSSTGSIDMV
70                                                 SNVSSTGSIDMV
71                                                 SNVSSTGSIDMV
72                                                 SNVSSTGSIDMV
73                                                 SNVSSTGSIDMV
74                                                 SNVSSTGSIDMV
75                                                 SNVSSTGSIDMV
76                                                 SNVSSTGSIDMV
77                                                 SNVSSTGSIDMV
78                                                 SNVSSTGSIDMV
79                                                 SNVSSTGSIDMV
80                                                 SNVSSTGSIDMV
81                                                 SNVSSTGSIDMV
82                                                 SNVSSTGSIDMV
83                                                    SSTGSIDMV
84                                                    SSTGSIDMV
85                                                    SSTGSIDMV
86                                                    SSTGSIDMV
87                                                    SSTGSIDMV
88                                                       GSIDMV
89                                                       GSIDMV

 1 DSPQLATLADEVSASLAKQGL
60 D
61 DSP
62 DSPQ
63 DSPQL
64 DSPQLA
65 DSPQLATL
70 D
71 DS
72 DSP
73 DSPQ
74 DSPQL
75 DSPQLA
76 DSPQLAT
77 DSPQLATL
78 DSPQLATLAD
79 DSPQLATLADE
```

80 DSPQLATLADEV
81 DSPQLATLADEVSA
82 DSPQLATLADEVSASLAKQGL
83 DSPQL
84 DSPQLA
85 DSPQLATL
86 DSPQLATLADEV
87 DSPQLATLADEVSASLAKQGL
88 DSPQLA
89 DSPQLATL
90 DSPQLATLADEV
91 TLADEVSASLAKQGL
92 LADEVSASLAKQGL
93 ADEVSASLAKQ
94 ADEVSASLAKQG
95 ADEVSASLAKQGL
96 EVSASLAKQGL
97 SASLAKQGL

**Soluble Tau 5 min**

```
  1 MAEPRQEFEVMEDHAGTYGLGDRKDQGGYTMHQDQEGDTDAGLKESPLQTPTEDGSEEPG
  2  AEPRQEFEVM

  1 SETSDAKSTPTAEDVTAPLVDEGAPGKQAAAQPHTEIPEGTTAEEAGIGDTPSLEDEAAG

  1 HVTQARMVSKSKDGTGSDDKKAKGADGKTKIATPRGAAPPGQKGQANATRIPAKTPPAPK

  1 TPPSSGEPPKSGDRSGYSSPGSPGTPGSRSRTPSLPTPPTREPKKVAVVRTPPKSPSSAK
  3                                               AVVRTPPKSPSSAK

  1 SRLQTAPVPMPDLKNVKSKIGSTENLKHQPGGGKVQIINKKLDLSNVQSKCGSKDNIKHV
  3 SRLQTAPVPMPDLKNV
  4     TAPVPMPDLKNV
  5                 KSKIGSTENL
  6                 KSKIGSTENLKHQPGGGKVQII
  7                           KHQPGGGKVQI
  8                           KHQPGGGKVQII
  9                           KHQPGGGKVQIINKKLDLSNV
 10                           KHQPGGGKVQIINKKLDLSNVQSKC
 11                                    QIINKKLDLSNV
 12                                      INKKLDLSNV
 13                                      INKKLDLSNVQSKC
 14                                       NKKLDLSNV
 15                                       NKKLDLSNVQSKC
 16                                         KLDLSNVQSKC
 17                                           DLSNVQSKC
 18                                           DLSNVQSKCGSKDNIKHV
 19                                             SNVQSKCGSKDNIKHV
 20                                              NVQSKCGSKDNIKHV
 21                                                QSKCGSKDNIKHV
 22                                                    GSKDNIKHV

  1 PGGGSVQIVYKPVDLSKVTSKCGSLGNIHHKPGGGQVEVKSEKLDFKDRVQSKIGSLDNI
 18 PGGGSVQI
 19 PGGGSVQI
 20 PGGGSVQI
 21 PGGGSVQI
 22 PGGGSVQI
 23   GGSVQIVYKPVDLSKVTSKC
 24         VYKPVDLS
 25         VYKPVDLSKV
 26         VYKPVDLSKVT
 27         VYKPVDLSKVTSK
 28         VYKPVDLSKVTSKC
 29         VYKPVDLSKVTSKCGSL
 30         VYKPVDLSKVTSKCGSLGNIH
 31          YKPVDLSKV
 32          YKPVDLSKVTSKC
 33          YKPVDLSKVTSKCGSLGNI
 34             DLSKVTSKC
 35             DLSKVTSKCGSLGNI
 36                  TSKCGSLGNI
 37                      GSLGNIHHKPGGGQVEV
 38                      GSLGNIHHKPGGGQVEVKSEKL
 39                                             DFKDRVQSKIGSLDNI
 40                                                   QSKIGSLDNI

  1 THVPGGGNKKIETHKLTFRENAKAKTDHGAEIVYKSPVVSGDTSPRHLSNVSSTGSIDMV
 41 THVPGGGNKKIETHKL
 42                 TFRENAKAKTDHGAEI
 43                 TFRENAKAKTDHGAEIV
 44                 TFRENAKAKTDHGAEIVY
```

45 TFRENAKAKTDHGAEIVYK
46 TFRENAKAKTDHGAEIVYKSPVVSGDTSPRHL
47 VYKSPVVSG
48 VYKSPVVSGD
49 VYKSPVVSGDT
50 VYKSPVVSGDTS
51 VYKSPVVSGDTSP
52 VYKSPVVSGDTSPR
53 VYKSPVVSGDTSPRH
54 VYKSPVVSGDTSPRHL
55 VYKSPVVSGDTSPRHLS
56 VYKSPVVSGDTSPRHLSN
57 VYKSPVVSGDTSPRHLSNV
58 VYKSPVVSGDTSPRHLSNVS
59 VYKSPVVSGDTSPRHLSNVSS
60 VYKSPVVSGDTSPRHLSNVSST
61 VYKSPVVSGDTSPRHLSNVSSTG
62 VYKSPVVSGDTSPRHLSNVSSTGS
63 VYKSPVVSGDTSPRHLSNVSSTGSID
64 VYKSPVVSGDTSPRHLSNVSSTGSIDM
65 VYKSPVVSGDTSPRHLSNVSSTGSIDMV
66 VYKSPVVSGDTSPRHLSNVSSTGSIDMV
67 VYKSPVVSGDTSPRHLSNVSSTGSIDMV
68 VYKSPVVSGDTSPRHLSNVSSTGSIDMV
69 VYKSPVVSGDTSPRHLSNVSSTGSIDMV
70 VYKSPVVSGDTSPRHLSNVSSTGSIDMV
71 VYKSPVVSGDTSPRHLSNVSSTGSIDMV
72 YKSPVVSGDTSPRHL
73 VSGDTSPRHL
74 SGDTSPRHL
75 SNVSSTGSIDM
76 SNVSSTGSIDMV
77 SNVSSTGSIDMV
78 SNVSSTGSIDMV
79 SNVSSTGSIDMV
80 SNVSSTGSIDMV
81 SNVSSTGSIDMV
82 SNVSSTGSIDMV
83 SNVSSTGSIDMV
84 SNVSSTGSIDMV
85 SNVSSTGSIDMV
86 SNVSSTGSIDMV
87 SNVSSTGSIDMV
88 SSTGSIDMV
89 SSTGSIDMV
90 SSTGSIDMV
91 SSTGSIDMV
92 SSTGSIDMV

1 DSPQLATLADEVSASLAKQGL
66 D
67 DSP
68 DSPQ
69 DSPQL
70 DSPQLA
71 DSPQLATL
77 D
78 DS
79 DSP
80 DSPQ
81 DSPQL
82 DSPQLA
83 DSPQLAT
84 DSPQLATL

85 DSPQLATLADEV
86 DSPQLATLADEVSA
87 DSPQLATLADEVSASLAKQGL
88 DSPQL
89 DSPQLA
90 DSPQLATL
91 DSPQLATLADEV
92 DSPQLATLADEVSASLAKQGL
93 LATLADEVSASLAKQGL
94 ATLADEVSASLAKQGL
95 TLADEVSASLAKQGL
96 LADEVSASLAKQGL
97 ADEVSASLAKQGL
98 EVSASLAKQGL
99 VSASLAKQGL
100 SASLAKQGL

**Soluble Tau 10 min**

```
  1 MAEPRQEFEVMEDHAGTYGLGDRKDQGGYTMHQDQEGDTDAGLKESPLQTPTEDGSEEPG

  1 SETSDAKSTPTAEDVTAPLVDEGAPGKQAAAQPHTEIPEGTTAEEAGIGDTPSLEDEAAG

  1 HVTQARMVSKSKDGTGSDDKKAKGADGKTKIATPRGAAPPGQKGQANATRIPAKTPPAPK

  1 TPPSSGEPPKSGDRSGYSSPGSPGTPGSRSRTPSLPTPPTREPKKVAVVRTPPKSPSSAK
  2                                               AVVRTPPKSPSSAK
  3                                               AVVRTPPKSPSSAK
  4                                                  RTPPKSPSSAK

  1 SRLQTAPVPMPDLKNVKSKIGSTENLKHQPGGGKVQIINKKLDLSNVQSKCGSKDNIKHV
  2 SRLQ
  3 SRLQTAPVPMPDLKNV
  4 SRLQTAPVPMPDLKNV
  5     TAPVPMPDLKNV
  6                 KSKIGSTENL
  7                 KSKIGSTENLKHQPGGGKVQI
  8                           KHQPGGGKVQI
  9                           KHQPGGGKVQII
 10                           KHQPGGGKVQIINKKLDLSNV
 11                           KHQPGGGKVQIINKKLDLSNVQSKC
 12                                    QIINKKLDLSNV
 13                                    QIINKKLDLSNVQSKC
 14                                      INKKLDLSNV
 15                                      INKKLDLSNVQS
 16                                      INKKLDLSNVQSKC
 17                                       NKKLDLSNV
 18                                       NKKLDLSNVQS
 19                                       NKKLDLSNVQSKC
 20                                         KLDLSNVQS
 21                                         KLDLSNVQSKC
 22                                           DLSNVQSKC
 23                                           DLSNVQSKCGSKDNIKHV
 24                                             SNVQSKCGSKDNIKHV
 25                                              NVQSKCGSKDNIKHV
 26                                                QSKCGSKDNIKHV
 27                                                  KCGSKDNIKHV
 28                                                    GSKDNIKHV
 29                                                    GSKDNIKHV
 30                                                    GSKDNIKHV
 31                                                    GSKDNIKHV
 32                                                        NIKHV

  1 PGGGSVQIVYKPVDLSKVTSKCGSLGNIHHKPGGGQVEVKSEKLDFKDRVQSKIGSLDNI
 23 PGGGSVQI
 24 PGGGSVQI
 25 PGGGSVQI
 26 PGGGSVQI
 27 PGGGSVQI
 28 PGGGSVQI
 29 PGGGSVQIV
 30 PGGGSVQIVYKPV
 31 PGGGSVQIVYKPVDLSKVTSKC
 32 PGGGSVQI
 33   GGSVQIVYKPVDLSKVTSKC
 34         VYKPVDLS
 35         VYKPVDLSKV
 36         VYKPVDLSKVT
 37         VYKPVDLSKVTSK
 38         VYKPVDLSKVTSKC
 39         VYKPVDLSKVTSKCG
```

```
 40         VYKPVDLSKVTSKCGSL
 41         VYKPVDLSKVTSKCGSLGNI
 42         VYKPVDLSKVTSKCGSLGNIH
 43          YKPVDLSKV
 44          YKPVDLSKVTSKC
 45            PVDLSKVTSKC
 46              DLSKVTSKC
 47              DLSKVTSKCGSLGNI
 48                   TSKCGSLGNI
 49                       GSLGNIHHKPGGGQVEV
 50                       GSLGNIHHKPGGGQVEVKS
 51                       GSLGNIHHKPGGGQVEVKSEKL
 52                             HHKPGGGQVEVKSEKL
 53                                             DFKDRVQSKIGSL
 54                                             DFKDRVQSKIGSLDNI
 55                                                   QSKIGSLDNI

  1 THVPGGGNKKIETHKLTFRENAKAKTDHGAEIVYKSPVVSGDTSPRHLSNVSSTGSIDMV
 56 THVPGGGNKKIETHKL
 57    PGGGNKKIETHKL
 58      GGNKKIETHKL
 59                 TFRENAKAKTD
 60                 TFRENAKAKTDH
 61                 TFRENAKAKTDHGAEI
 62                 TFRENAKAKTDHGAEIV
 63                 TFRENAKAKTDHGAEIVY
 64                 TFRENAKAKTDHGAEIVYK
 65                 TFRENAKAKTDHGAEIVYKSPVVSGDTSPRHL
 66                  FRENAKAKTDHGAEI
 67                                 VYKSPVVSG
 68                                 VYKSPVVSGD
 69                                 VYKSPVVSGDT
 70                                 VYKSPVVSGDTS
 71                                 VYKSPVVSGDTSP
 72                                 VYKSPVVSGDTSPR
 73                                 VYKSPVVSGDTSPRH
 74                                 VYKSPVVSGDTSPRHL
 75                                 VYKSPVVSGDTSPRHLS
 76                                 VYKSPVVSGDTSPRHLSN
 77                                 VYKSPVVSGDTSPRHLSNV
 78                                 VYKSPVVSGDTSPRHLSNVS
 79                                 VYKSPVVSGDTSPRHLSNVSS
 80                                 VYKSPVVSGDTSPRHLSNVSST
 81                                 VYKSPVVSGDTSPRHLSNVSSTG
 82                                 VYKSPVVSGDTSPRHLSNVSSTGS
 83                                 VYKSPVVSGDTSPRHLSNVSSTGSID
 84                                 VYKSPVVSGDTSPRHLSNVSSTGSIDM
 85                                 VYKSPVVSGDTSPRHLSNVSSTGSIDMV
 86                                 VYKSPVVSGDTSPRHLSNVSSTGSIDMV
 87                                 VYKSPVVSGDTSPRHLSNVSSTGSIDMV
 88                                 VYKSPVVSGDTSPRHLSNVSSTGSIDMV
 89                                 VYKSPVVSGDTSPRHLSNVSSTGSIDMV
 90                                 VYKSPVVSGDTSPRHLSNVSSTGSIDMV
 91                                 VYKSPVVSGDTSPRHLSNVSSTGSIDMV
 92                                 VYKSPVVSGDTSPRHLSNVSSTGSIDMV
 93                                  YKSPVVSGDTSPRHL
 94                                       VSGDTSPRHL
 95                                       VSGDTSPRHLSNV
 96                                        SGDTSPRHL
 97                                                 SNVSSTGSIDM
 98                                                 SNVSSTGSIDMV
 99                                                 SNVSSTGSIDMV
100                                                 SNVSSTGSIDMV
101                                                 SNVSSTGSIDMV
```

```
102                                                 SNVSSTGSIDMV
103                                                 SNVSSTGSIDMV
104                                                 SNVSSTGSIDMV
105                                                 SNVSSTGSIDMV
106                                                 SNVSSTGSIDMV
107                                                 SNVSSTGSIDMV
108                                                 SNVSSTGSIDMV
109                                                 SNVSSTGSIDMV
110                                                 SNVSSTGSIDMV
111                                                 SNVSSTGSIDMV
112                                                    SSTGSIDMV
113                                                    SSTGSIDMV
114                                                    SSTGSIDMV
115                                                    SSTGSIDMV
116                                                    SSTGSIDMV
117                                                    SSTGSIDMV
118                                                       GSIDMV
119                                                       GSIDMV

  1 DSPQLATLADEVSASLAKQGL
 86 D
 87 DSP
 88 DSPQ
 89 DSPQL
 90 DSPQLA
 91 DSPQLAT
 92 DSPQLATL
 99 D
100 DS
101 DSP
102 DSPQ
103 DSPQL
104 DSPQLA
105 DSPQLAT
106 DSPQLATL
107 DSPQLATLAD
108 DSPQLATLADE
109 DSPQLATLADEV
110 DSPQLATLADEVSA
111 DSPQLATLADEVSASLAKQGL
112 DSPQL
113 DSPQLA
114 DSPQLATL
115 DSPQLATLADEV
116 DSPQLATLADEVSA
117 DSPQLATLADEVSASLAKQGL
118 DSPQLATL
119 DSPQLATLADEV
120 DSPQLATLADEV
121     LATLADEVSASLAKQGL
122      ATLADEVSASLAKQGL
123       TLADEVSASLAKQGL
124        LADEVSASLAKQGL
125         ADEVSASLAKQ
126         ADEVSASLAKQGL
127           EVSASLAKQGL
128            VSASLAKQGL
129             SASLAKQGL
```

**Soluble Tau 120 min**

```
 1 MAEPRQEFEVMEDHAGTYGLGDRKDQGGYTMHQDQEGDTDAGLKESPLQTPTEDGSEEPG
 2  AEPRQEFEV
 3  AEPRQEFEVM

 1 SETSDAKSTPTAEDVTAPLVDEGAPGKQAAAQPHTEIPEGTTAEEAGIGDTPSLEDEAAG
 4                                               GIGDTPSLEDEAAG

 1 HVTQARMVSKSKDGTGSDDKKAKGADGKTKIATPRGAAPPGQKGQANATRIPAKTPPAPK
 4 HVTQA

 1 TPPSSGEPPKSGDRSGYSSPGSPGTPGSRSRTPSLPTPPTREPKKVAVVRTPPKSPSSAK
 5                                               AVVRTPPKSPSSA
 6                                               AVVRTPPKSPSSAK
 7                                               AVVRTPPKSPSSAK
 8                                                 VRTPPKSPSSAK
 9                                                  RTPPKSPSSAK
10                                                  RTPPKSPSSAK
11                                                          SAK

 1 SRLQTAPVPMPDLKNVKSKIGSTENLKHQPGGGKVQIINKKLDLSNVQSKCGSKDNIKHV
 6 SRLQ
 7 SRLQTAPVPMPDLKNV
 8 SRLQTAPVPMPDLKNV
 9 SRLQ
10 SRLQTAPVPMPDLKNV
11 SRLQTAPVPMPDLKNV
12  RLQTAPVPMPDLKNV
13    QTAPVPMPDLKNV
14     TAPVPMPDLKNV
15                 KSKIGSTENL
16                 KSKIGSTENLKHQPGGGKV
17                 KSKIGSTENLKHQPGGGKVQI
18                 KSKIGSTENLKHQPGGGKVQII
19                 KSKIGSTENLKHQPGGGKVQIINK
20                 KSKIGSTENLKHQPGGGKVQIINKKL
21                 KSKIGSTENLKHQPGGGKVQIINKKLDLSNV
22                 KSKIGSTENLKHQPGGGKVQIINKKLDLSNVQS
23                        ENLKHQPGGGKVQIINKKLDLSNV
24                           KHQPGGGKVQI
25                           KHQPGGGKVQII
26                           KHQPGGGKVQIINK
27                           KHQPGGGKVQIINKKLDLSNV
28                           KHQPGGGKVQIINKKLDLSNVQS
29                           KHQPGGGKVQIINKKLDLSNVQSKC
30                                    QIINKKLDLSNV
31                                    QIINKKLDLSNVQSKC
32                                      INKKLDLSNV
33                                      INKKLDLSNVQS
34                                      INKKLDLSNVQSKC
35                                      INKKLDLSNVQSKCGSKDNIKHV
36                                       NKKLDLSNV
37                                       NKKLDLSNVQS
38                                       NKKLDLSNVQSKC
39                                       NKKLDLSNVQSKCGSKDNIKHV
40                                         KLDLSNVQS
41                                         KLDLSNVQSKC
42                                         KLDLSNVQSKCGSKDNIKHV
43                                           DLSNVQSKC
44                                           DLSNVQSKCGSKDNIKHV
45                                             SNVQSKCGSKDNIKHV
46                                              NVQSKCGSKDNIKHV
47                                                QSKCGSKDNIKHV
```

```
48                                                   KCGSKDNIKHV
49                                                     GSKDNIKHV
50                                                     GSKDNIKHV
51                                                     GSKDNIKHV
52                                                     GSKDNIKHV
53                                                     GSKDNIKHV
54                                                         NIKHV

 1 PGGGSVQIVYKPVDLSKVTSKCGSLGNIHHKPGGGQVEVKSEKLDFKDRVQSKIGSLDNI
35 PGGGSVQI
39 PGGGSVQI
42 PGGGSVQI
44 PGGGSVQI
45 PGGGSVQI
46 PGGGSVQI
47 PGGGSVQI
48 PGGGSVQI
49 PGGGSVQI
50 PGGGSVQIV
51 PGGGSVQIVYKPV
52 PGGGSVQIVYKPVDLSKV
53 PGGGSVQIVYKPVDLSKVTSKC
54 PGGGSVQI
55   GGSVQIVYKPVDLSKV
56   GGSVQIVYKPVDLSKVTSKC
57       QIVYKPVDLSKV
58       QIVYKPVDLSKVTSKC
59        IVYKPVDLSKVTSKC
60         VYKPVDLS
61         VYKPVDLSK
62         VYKPVDLSKV
63         VYKPVDLSKVT
64         VYKPVDLSKVTSK
65         VYKPVDLSKVTSKC
66         VYKPVDLSKVTSKCG
67         VYKPVDLSKVTSKCGS
68         VYKPVDLSKVTSKCGSL
69         VYKPVDLSKVTSKCGSLG
70         VYKPVDLSKVTSKCGSLGN
71         VYKPVDLSKVTSKCGSLGNI
72         VYKPVDLSKVTSKCGSLGNIH
73         VYKPVDLSKVTSKCGSLGNIHH
74         VYKPVDLSKVTSKCGSLGNIHHKPGGGQVEVKSEKL
75          YKPVDLSKV
76          YKPVDLSKVTSKC
77          YKPVDLSKVTSKCGSLGNI
78           KPVDLSKVTSKC
79            PVDLSKVTSKC
80              DLSKVTSKC
81              DLSKVTSKCGSLGNI
82               LSKVTSKC
83                SKVTSKCGSLGNI
84                   TSKCGSLGNI
85                       GSLGNIHHKPGG
86                       GSLGNIHHKPGGG
87                       GSLGNIHHKPGGGQVEV
88                       GSLGNIHHKPGGGQVEVKS
89                       GSLGNIHHKPGGGQVEVKSE
90                       GSLGNIHHKPGGGQVEVKSEK
91                       GSLGNIHHKPGGGQVEVKSEKL
92                       GSLGNIHHKPGGGQVEVKSEKLD
93                         LGNIHHKPGGGQVEVKSEKL
94                          GNIHHKPGGGQVEVKSEKL
95                             HHKPGGGQVEVKSEKL
```

```
 96                                      EVKSEKL
 97                                             DFKDRVQSKIGSL
 98                                             DFKDRVQSKIGSLDNI
 99                                             DFKDRVQSKIGSLDNI
100                                             DFKDRVQSKIGSLDNI
101                                                   QSKIGSLDNI
102                                                   QSKIGSLDNI
103                                                          DNI

  1 THVPGGGNKKIETHKLTFRENAKAKTDHGAEIVYKSPVVSGDTSPRHLSNVSSTGSIDMV
 99 T
100 THVPGGGNKKIETHKL
102 THVPGGGNKKIETHKL
103 THVPGGGNKKIETHKL
104 THVPGGGNKKIETHKL
105    PGGGNKKIETHKL
106     GGGNKKIETHKL
107      GGNKKIETHKL
108       GNKKIETHKL
109                 TFRENAKAKTD
110                 TFRENAKAKTDH
111                 TFRENAKAKTDHGA
112                 TFRENAKAKTDHGAEI
113                 TFRENAKAKTDHGAEIV
114                 TFRENAKAKTDHGAEIVY
115                 TFRENAKAKTDHGAEIVYK
116                 TFRENAKAKTDHGAEIVYKSPVVSG
117                 TFRENAKAKTDHGAEIVYKSPVVSGDTSPRHL
118                 TFRENAKAKTDHGAEIVYKSPVVSGDTSPRHLSNV
119                  FRENAKAKTDHGAEI
120                                 VYKSPVVSG
121                                 VYKSPVVSGD
122                                 VYKSPVVSGDT
123                                 VYKSPVVSGDTS
124                                 VYKSPVVSGDTSPR
125                                 VYKSPVVSGDTSPRH
126                                 VYKSPVVSGDTSPRHL
127                                 VYKSPVVSGDTSPRHLS
128                                 VYKSPVVSGDTSPRHLSN
129                                 VYKSPVVSGDTSPRHLSNV
130                                 VYKSPVVSGDTSPRHLSNVS
131                                 VYKSPVVSGDTSPRHLSNVSS
132                                 VYKSPVVSGDTSPRHLSNVSST
133                                 VYKSPVVSGDTSPRHLSNVSSTG
134                                 VYKSPVVSGDTSPRHLSNVSSTGS
135                                 VYKSPVVSGDTSPRHLSNVSSTGSID
136                                 VYKSPVVSGDTSPRHLSNVSSTGSIDM
137                                 VYKSPVVSGDTSPRHLSNVSSTGSIDMV
138                                 VYKSPVVSGDTSPRHLSNVSSTGSIDMV
139                                 VYKSPVVSGDTSPRHLSNVSSTGSIDMV
140                                 VYKSPVVSGDTSPRHLSNVSSTGSIDMV
141                                 VYKSPVVSGDTSPRHLSNVSSTGSIDMV
142                                 VYKSPVVSGDTSPRHLSNVSSTGSIDMV
143                                 VYKSPVVSGDTSPRHLSNVSSTGSIDMV
144                                 VYKSPVVSGDTSPRHLSNVSSTGSIDMV
145                                  YKSPVVSGDTSPRHL
146                                  YKSPVVSGDTSPRHLSNVSSTGSIDMV
147                                      VSGDTSPRHL
148                                      VSGDTSPRHLSNV
149                                      VSGDTSPRHLSNVSSTGSIDMV
150                                      VSGDTSPRHLSNVSSTGSIDMV
151                                       SGDTSPRHL
152                                                SNVSSTGSIDM
153                                                SNVSSTGSIDMV
```

```
154                                                 SNVSSTGSIDMV
155                                                 SNVSSTGSIDMV
156                                                 SNVSSTGSIDMV
157                                                 SNVSSTGSIDMV
158                                                 SNVSSTGSIDMV
159                                                 SNVSSTGSIDMV
160                                                 SNVSSTGSIDMV
161                                                 SNVSSTGSIDMV
162                                                 SNVSSTGSIDMV
163                                                 SNVSSTGSIDMV
164                                                 SNVSSTGSIDMV
165                                                 SNVSSTGSIDMV
166                                                    SSTGSIDMV
167                                                    SSTGSIDMV
168                                                    SSTGSIDMV
169                                                    SSTGSIDMV
170                                                    SSTGSIDMV
171                                                    SSTGSIDMV
172                                                       GSIDMV
173                                                       GSIDMV
174                                                       GSIDMV

  1 DSPQLATLADEVSASLAKQGL
138 D
139 DS
140 DSP
141 DSPQ
142 DSPQL
143 DSPQLA
144 DSPQLATL
146 DSPQLATL
149 DSPQLATL
150 DSPQLATLADEV
154 D
155 DS
156 DSP
157 DSPQ
158 DSPQL
159 DSPQLA
160 DSPQLATL
161 DSPQLATLAD
162 DSPQLATLADE
163 DSPQLATLADEV
164 DSPQLATLADEVSA
165 DSPQLATLADEVSASLAKQGL
166 DSPQL
167 DSPQLA
168 DSPQLATL
169 DSPQLATLADEV
170 DSPQLATLADEVSA
171 DSPQLATLADEVSASLAKQGL
172 DSPQLA
173 DSPQLATL
174 DSPQLATLADEV
175 DSPQLATLADEV
176  SPQLATLADEVSASLAKQGL
177     LATLADEVSASLAKQGL
178      ATLADEVSASLAKQGL
179       TLADEVSASLAKQGL
180        LADEVSASLAKQGL
181         ADEVSASLAKQ
182         ADEVSASLAKQG
183         ADEVSASLAKQGL
184           EVSASLAKQGL
```

185 VSASLAKQGL
186 SASLAKQGL

Supplemental data 2

```
CLUSTAL O(1.2.4) multiple sequence alignment

sp|P10637-2|MOUSE                   MADPRQEFDTMEDHAG--------------DYTLLQDQEGDMDHGLKESPPQPPADDGAE    46
sp|P19332|RAT                       MAEPRQEFDTMEDQAG--------------DYTMLQDQEGDMDHGLKESPPQPPADDGSE    46
tr|F6UXK3|CALJA                     MAEPRQEFNVMEDHTGTYGLED-------------QDQEGDTDTGLKESPLQTPAEDGSE    47
sp|O02828|CAPHI                     MAEPRQEFDVMEDHA-------------QGDY-TLQDHEGDMEPGLKESPLQTPADDGSE    46
sp|P29172|BOVIN                     MAEPRQEFDVMEDHA-------------QGDY-TLQDQEGDMDPGLKESPLQTPADDGSE    46
sp|P10636-8|HUMAN                   MAEPRQEFEVMEDHAGTYGLGDRKD---QGGYTMHQDQEGDTDAGLKESPLQTPTEDGSE    57
tr|A0A2K6S2L5|SAIBB                 MAEPRQEFDVMEDHAGTYGLEDQ-------------DQEGDTDTGLKESPLQTPAEDGSE    47
tr|A0A8P0SZT9|CANLF                 MAEPRQEFTVMEDHAGTYG----KDLPSQGGYTLLQDHEGDVDHGLKESPLQTPADDGSE    56
tr|A0A7N5KFP8|AILME                 MAEPRQEFTATEDHAGTYGMEERKDLPSQGGYTLLQDHEGDMDHGLKESPLQTPADDGSE    60
                                    **:***** . **::                     *:*** : ****** * *::**:*

sp|P10637-2|MOUSE                   EPGSETSDAKSTPTAEDVTAPLVDERAPDKQAAAQPHTEIPEGITAEEAGIGDTPNQEDQ    106
sp|P19332|RAT                       EPGSETSDAKSTPTAEDVTAPLVEERAPDKQATAQSHTEIPEGTTAEEAGIGDTPNMEDQ    106
tr|F6UXK3|CALJA                     EPGSESSDAKSTPTVEDVTAPLVDERAPGKQAAAQPHTEIPEGTTAEEAGIGDTPTPEDQ    107
sp|O02828|CAPHI                     EPGSETSDAKSTPTAE-----------------------------AEEAGIGDTSNLEDQ    77
sp|P29172|BOVIN                     EPGSETSDAKSTPTAEDATAPLVDEGAPGEQAAAQAPAEIPEGTAAEEAGIGDTSNLEDQ    106
sp|P10636-8|HUMAN                   EPGSETSDAKSTPTAEDVTAPLVDEGAPGKQAAAQPHTEIPEGTTAEEAGIGDTPSLEDE    117
tr|A0A2K6S2L5|SAIBB                 EPGSETSDAKSTPTAEDVTAPLVDERAPGEQAAAQPHTEIPEGTTAEEAGIGDTPTPEDQ    107
tr|A0A8P0SZT9|CANLF                 EPGSETSDAKSTPTAEDVTAPLVDEGTPGEQAAAQPPMEIPEGATAEEAGIGDTPNLEDQ    116
tr|A0A7N5KFP8|AILME                 EPGSETSDAKSTPTAEDVTAPLVDEGAPGEQAAAQPHTEIPEGATAEEAGIGDTPSLEDQ    120
                                    *****:********.*                             ********* . **:

sp|P10637-2|MOUSE                   AAGHVTQARVA--SKDRTGNDEKKAKGADGKTGAKIATPRGAASPAQKGTSNATRIPAKT    164
sp|P19332|RAT                       AAGHVTQARVAGVSKDRTGNDEKKAKGADGKTGAKIATPRGAATPGQKGTSNATRIPAKT    166
tr|F6UXK3|CALJA                     AAGHVTQARMVSKSKDGTGGDDKKAKGADGK--TKIATPRGTAPPGQKGQANATRIPAKT    165
sp|O02828|CAPHI                     AAGHVTQARMVSKGKDGTGPDDKKAKGADGKPGTKIATPRGAAPPGQKGQANATRIPAKT    137
sp|P29172|BOVIN                     AAGHVTQARMVSKGKDGTGPDDKKTKGADGKPGTKIATPRGAAPPGQKGQANATRIPAKT    166
sp|P10636-8|HUMAN                   AAGHVTQARMVSKSKDGTGSDDKKAKGADGKT--KIATPRGAAPPGQKGQANATRIPAKT    175
tr|A0A2K6S2L5|SAIBB                 AAGHVTQARMVSKSKDGTGGDDKKAKGADGKT--KIATPRGTAPPGQKGQANATRIPAKT    165
tr|A0A8P0SZT9|CANLF                 AAGHVTQARMVSKGKDGTGTDDKKAKGADGKTGTKIATPRGATPSGQKGQANATRIPAKT    176
tr|A0A7N5KFP8|AILME                 AAGHVTQARMVSKGKDGTGPEDKKAKGADGKTGTKIATPRGAAPPGQKGQANATRIPAKT    180
                                    *********:.  .** ** ::**:******   *******::  .*** :*********

sp|P10637-2|MOUSE                   TPSPKTPPG----------------SGEPPKSGERSGYSSPGSPGTPGSRSRTPSLPTPP    208
sp|P19332|RAT                       TPSPKTPPG----------------SGEPPKSGERSGYSSPGSPGTPGSRSRTPSLPTPP    210
tr|F6UXK3|CALJA                     PPAPKTPPS----------------SGETPKSGDRSGYSSPGSPGTPGSRSRTPSLPTPP    209
sp|O02828|CAPHI                     TPTPKTSPG----------------TGESGKSGDRSGYSSPGSPGTPGSRSRTPSLPTPP    181
sp|P29172|BOVIN                     TPTPKTSPATMQVQKKPPPAGAKSERGESGKSGDRSGYSSPGSPGTPGSRSRTPSLPTPP    226
sp|P10636-8|HUMAN                   PPAPKTPPS----------------SGEPPKSGDRSGYSSPGSPGTPGSRSRTPSLPTPP    219
```

```
tr|A0A2K6S2L5|SAIBB                 PPTPKTPPG----------------SGETPKSGDRSGYSSPGSPGTPGSRSRTPSLPTPP    209
tr|A0A8P0SZT9|CANLF                 TPSPKTPPG-----------------GESGKSGDRSGYSSPGSPGTPGSRSRTPSLPTPP    219
tr|A0A7N5KFP8|AILME                 TPSPKTPPG------------------ESGKSGDRSGYSSPGSPGTPGSRSRTPSLPTPP    222
                                     *:*** *.                  *  ***:**************************

sp|P10637-2|MOUSE                   TREPKKVAVVRTPPKSPSASKSRLQTAPVPMPDLKNVRSKIGSTENLKHQPGGGKVQIIN    268
sp|P19332|RAT                       TREPKKVAVVRTPPKSPSASKSRLQTAPVPMPDLKNVRSKIGSTENLKHQPGGGKVQIIN    270
tr|F6UXK3|CALJA                     TREPKKVAVVRTPPKSPSSTKSRLQTAPVPMPDLKNVKSKIGSTENLKHQPGGGKVQIIN    269
sp|O02828|CAPHI                     TREPKKVAVVRTPPKSPSAAKSRLQAAPGPMPDLKNVKSKIGSTENLKHQPGGGKVQIIN    241
sp|P29172|BOVIN                     TREPKKVAVVRTPPKSPSAAKSRLQAAPGPMPDLKNVKSKIGSTENLKHQPGGGKVQIIN    286
sp|P10636-8|HUMAN                   TREPKKVAVVRTPPKSPSSAKSRLQTAPVPMPDLKNVKSKIGSTENLKHQPGGGKVQIIN    279
tr|A0A2K6S2L5|SAIBB                 TREPKKVAVVRTPPKSPSSAKSRLQTAPVPMPDLKNVKSKIGSTENLKHQPGGGKVQIIN    269
tr|A0A8P0SZT9|CANLF                 TREPKKVAVVRTPPKSPSAAKSRLQTAPVPMPDLKNVRSKIGSTENLKHQPGGGKVQIIN    279
tr|A0A7N5KFP8|AILME                 TREPKKVAVVRTPPKSPSSAKSRLQTAPVPMPDLKNVRSKIGSTENLKHQPGGGKVQIIN    282
                                    ******************::*****:** ********:**********************

sp|P10637-2|MOUSE                   KKLDLSNVQSKCGSKDNIKHVPGGGSVQIVYKPVDLSKVTSKCGSLGNIHHKPGGGQVEV    328
sp|P19332|RAT                       KKLDLSNVQSKCGSKDNIKHVPGGGSVHIVYKPVDLSKVTSKCGSLGNIHHKPGGGQVEV    330
tr|F6UXK3|CALJA                     KKLDLSNVQSKCGSKDNIKHVPGGGSVQIVYKPVDLSKVTSKCGSLGNIHHKPGGGQVEV    329
sp|O02828|CAPHI                     KKLDLSNVQSKCGSKDNIKHVPGGGSVQIVYKPVDLSKVTSKCGSLGNIHHKPGGGQVEV    301
sp|P29172|BOVIN                     KKLDLSNVQSKCGSKDNIKHVPGGGSVQIVYKPVDLSKVTSKCGSLGNIHHKPGGGQVEV    346
sp|P10636-8|HUMAN                   KKLDLSNVQSKCGSKDNIKHVPGGGSVQIVYKPVDLSKVTSKCGSLGNIHHKPGGGQVEV    339
tr|A0A2K6S2L5|SAIBB                 KKLDLSNVQSKCGSKDNIKHVPGGGSVQIVYKPVDLSKVTSKCGSLGNIHHKPGGGQVEV    329
tr|A0A8P0SZT9|CANLF                 KKLDLSNVQSKCGSKDNIKHVPGGGSVQIVYKPVDLSKVTSKCGSLGNIHHKPGGGQVEV    339
tr|A0A7N5KFP8|AILME                 KKLDLSNVQSKCGSKDNIKHVPGGGSVQIVYKPVDLSKVTSKCG----------GGQVEV    332
                                    ***************************:****************          ******

sp|P10637-2|MOUSE                   KSEKLDFKDRVQSKIGSLDNITHVPGGGNKKIETHKLTFRENAKAKTDHGAEIVYKSPVV    388
sp|P19332|RAT                       KSEKLDFKDRVQSKIGSLDNITHVPGGGNKKIETHKLTFRENAKAKTDHGAEIVYKSPVV    390
tr|F6UXK3|CALJA                     KSEKLDFKDRVQSKIGSLDNITHVPGGGNKKIETHKLTFRENAKAKTDHGAEIVYKSPVV    389
sp|O02828|CAPHI                     KSEKLDFKDRVQSKIGSLDNITHVPGGGNKKIETHKLTFRENAKAKTDHGAEIVYKSPVV    361
sp|P29172|BOVIN                     KSEKLDFKDRVQSKIGSLDNITHVPGGGNKKIETHKLTFRENAKAKTDHGAEIVYKSPVV    406
sp|P10636-8|HUMAN                   KSEKLDFKDRVQSKIGSLDNITHVPGGGNKKIETHKLTFRENAKAKTDHGAEIVYKSPVV    399
tr|A0A2K6S2L5|SAIBB                 KSEKLDFKDRVQSKIGSLDNITHVPGGGNKKIETHKLTFRENAKAKTDHGAEIVYKSPVV    389
tr|A0A8P0SZT9|CANLF                 KSEKLDFKDRVQSKIGSLDNITHVPGGGNKKIETHKLTFRENAKAKTDHGAEIVYKSPVV    399
tr|A0A7N5KFP8|AILME                 KSEKLDFKDRVQSKIGSLDNITHVPGGGNKKIETHKLTFRENAKAKTDHGAEIVYKSPVV    392
                                    ************************************************************

sp|P10637-2|MOUSE                   SGDTSPRHLSNVSSTGSIDMVDSPQLATLADEVSASLAKQGL    430
sp|P19332|RAT                       SGDTSPRHLSNVSSTGSIDMVDSPQLATLADEVSASLAKQGL    432
tr|F6UXK3|CALJA                     SGDTSPRHLSNVSSTGSIDMVDSPQLATLADEVSASLAKQGL    431
sp|O02828|CAPHI                     SGDTSPRHLSNVSSTGSIDMVDSPQLATLADEVSASLAKQGL    403
```

```
sp|P29172|BOVIN            SGDTSPRHLSNVSSTGSIDMVDSPQLATLADEVSASLAKQGL     448
sp|P10636-8|HUMAN          SGDTSPRHLSNVSSTGSIDMVDSPQLATLADEVSASLAKQGL     441
tr|A0A2K6S2L5|SAIBB        SGDTSPRHLSNVSSTGSIDMADSAQLATLADEVSASLAKQGL     431
tr|A0A8P0SZT9|CANLF        SGDTSPRHLSNVSSTGSIDMVDSPQLATLADEVSASLAKQGL     441
tr|A0A7N5KFP8|AILME        SGDTSPRHLSNVSSTGSIDMVDSPQLATLADEVSASLAKQGL     434
                           ********************.** ******************
```

**Supplemental data 3**

**Tau fibrils (β-sheets highlighted in yellow) 15 sec**

```
 1 MAEPRQEFEVMEDHAGTYGLGDRKDQGGYTMHQDQEGDTDAGLKESPLQTPTEDGSEEPG

 1 SETSDAKSTPTAEDVTAPLVDEGAPGKQAAAQPHTEIPEGTTAEEAGIGDTPSLEDEAAG

 1 HVTQARMVSKSKDGTGSDDKKAKGADGKTKIATPRGAAPPGQKGQANATRIPAKTPPAPK

 1 TPPSSGEPPKSGDRSGYSSPGSPGTPGSRSRTPSLPTPPTREPKKVAVVRTPPKSPSSAK

 1 SRLQTAPVPMPDLKNVKSKIGSTENLKHQPGGGKVQIINKKLDLSNVQSKCGSKDNIKHV
 2     TAPVPMPDLKNV
 3                 KSKIGSTENL

 1 PGGGSVQIVYKPVDLSKVTSKCGSLGNIHHKPGGGQVEVKSEKLDFKDRVQSKIGSLDNI
 4         VYKPVDLSKV

 1 THVPGGGNKKIETHKLTFRENAKAKTDHGAEIVYKSPVVSGDTSPRHLSNVSSTGSIDMV
 5 THVPGGGNKKIETHKL
 6                 TFRENAKA
 7                 TFRENAKAKTDH
 8                 TFRENAKAKTDHGAEI
 9                 TFRENAKAKTDHGAEIV
10                                 VYKSPVVSG
11                                 VYKSPVVSGD
12                                 VYKSPVVSGDT
13                                 VYKSPVVSGDTS
14                                 VYKSPVVSGDTSP
15                                 VYKSPVVSGDTSPRH
16                                 VYKSPVVSGDTSPRHL
17                                 VYKSPVVSGDTSPRHLSN
18                                 VYKSPVVSGDTSPRHLSNV
19                                                 SNVSSTGSIDM
20                                                 SNVSSTGSIDMV
21                                                 SNVSSTGSIDMV
22                                                 SNVSSTGSIDMV
23                                                 SNVSSTGSIDMV
24                                                 SNVSSTGSIDMV
25                                                 SNVSSTGSIDMV
26                                                 SNVSSTGSIDMV
27                                                 SNVSSTGSIDMV
28                                                 SNVSSTGSIDMV
29                                                    SSTGSIDMV
30                                                    SSTGSIDMV
31                                                    SSTGSIDMV
32                                                    SSTGSIDMV

 1 DSPQLATLADEVSASLAKQGL
20 D
21 DSP
22 DSPQ
23 DSPQL
24 DSPQLA
25 DSPQLATL
26 DSPQLATLADEV
27 DSPQLATLADEVSA
28 DSPQLATLADEVSASLAKQGL
29 DSPQL
30 DSPQLA
31 DSPQLATL
32 DSPQLATLADEVSASLAKQGL
33         ADEVSASLAKQG
```

| | |
|---|---|
| 34 | SASLAKQGL |
| 35 | ASLAKQGL |

**Tau fibrils 1 min**

```
 1 MAEPRQEFEVMEDHAGTYGLGDRKDQGGYTMHQDQEGDTDAGLKESPLQTPTEDGSEEPG

 1 SETSDAKSTPTAEDVTAPLVDEGAPGKQAAAQPHTEIPEGTTAEEAGIGDTPSLEDEAAG

 1 HVTQARMVSKSKDGTGSDDKKAKGADGKTKIATPRGAAPPGQKGQANATRIPAKTPPAPK

 1 TPPSSGEPPKSGDRSGYSSPGSPGTPGSRSRTPSLPTPPTREPKKVAVVRTPPKSPSSAK
 2                                               AVVRTPPKSPSSA
 3                                               AVVRTPPKSPSSAK
 4                                                  RTPPKSPSSAK

 1 SRLQTAPVPMPDLKNVKSKIGSTENLKHQPGGGKVQIINKKLDLSNVQSKCGSKDNIKHV
 3 SRLQTAPVPMPDLKNV
 4 SRLQTAPVPMPDLKNV
 5     TAPVPMPDLKNV
 6      APVPMPDLKNV
 7                 KSKIGSTENL
 8                           KHQPGGGKVQI
 9                           KHQPGGGKVQII
10                                      INKKLDLSNV
11                                       NKKLDLSNV
12                                                    GSKDNIKHV
13                                                          KHV

 1 PGGGSVQIVYKPVDLSKVTSKCGSLGNIHHKPGGGQVEVKSEKLDFKDRVQSKIGSLDNI
12 PGGGSVQI
13 PGGGSVQI
14         VYKPVDLSK
15         VYKPVDLSKV
16         VYKPVDLSKVT
17         VYKPVDLSKVTSK
18         VYKPVDLSKVTSKC
19          YKPVDLSKV
20                                             DFKDRVQSKIGSLDNI
21                                                   QSKIGSLDNI
22                                                          DNI

 1 THVPGGGNKKIETHKLTFRENAKAKTDHGAEIVYKSPVVSGDTSPRHLSNVSSTGSIDMV
22 THVPGGGNKKIETHKL
23 THVPGGGNKKIETHKL
24 THVPGGGNKKIETHKLT
25    PGGGNKKIETHKL
26      GGNKKIETHKL
27       GNKKIETHKL
28                 TFRENAKA
29                 TFRENAKAKTD
30                 TFRENAKAKTDH
31                 TFRENAKAKTDHGAE
32                 TFRENAKAKTDHGAEI
33                 TFRENAKAKTDHGAEIV
34                 TFRENAKAKTDHGAEIVY
35                 TFRENAKAKTDHGAEIVYK
36                 TFRENAKAKTDHGAEIVYKSPV
37                 TFRENAKAKTDHGAEIVYKSPVV
38                 TFRENAKAKTDHGAEIVYKSPVVS
39                 TFRENAKAKTDHGAEIVYKSPVVSG
40                 TFRENAKAKTDHGAEIVYKSPVVSGD
41                 TFRENAKAKTDHGAEIVYKSPVVSGDTS
42                 TFRENAKAKTDHGAEIVYKSPVVSGDTSP
43                 TFRENAKAKTDHGAEIVYKSPVVSGDTSPRH
44                 TFRENAKAKTDHGAEIVYKSPVVSGDTSPRHL
45                 TFRENAKAKTDHGAEIVYKSPVVSGDTSPRHLSN
```

```
 46                 TFRENAKAKTDHGAEIVYKSPVVSGDTSPRHLSNV
 47                                 VYKSPVVS
 48                                 VYKSPVVSG
 49                                 VYKSPVVSGD
 50                                 VYKSPVVSGDT
 51                                 VYKSPVVSGDTS
 52                                 VYKSPVVSGDTSP
 53                                 VYKSPVVSGDTSPR
 54                                 VYKSPVVSGDTSPRH
 55                                 VYKSPVVSGDTSPRHL
 56                                 VYKSPVVSGDTSPRHLS
 57                                 VYKSPVVSGDTSPRHLSN
 58                                 VYKSPVVSGDTSPRHLSNV
 59                                 VYKSPVVSGDTSPRHLSNVS
 60                                 VYKSPVVSGDTSPRHLSNVSS
 61                                 VYKSPVVSGDTSPRHLSNVSST
 62                                 VYKSPVVSGDTSPRHLSNVSSTG
 63                                 VYKSPVVSGDTSPRHLSNVSSTGS
 64                                 VYKSPVVSGDTSPRHLSNVSSTGSID
 65                                 VYKSPVVSGDTSPRHLSNVSSTGSIDM
 66                                 VYKSPVVSGDTSPRHLSNVSSTGSIDMV
 67                                 VYKSPVVSGDTSPRHLSNVSSTGSIDMV
 68                                 VYKSPVVSGDTSPRHLSNVSSTGSIDMV
 69                                 VYKSPVVSGDTSPRHLSNVSSTGSIDMV
 70                                 VYKSPVVSGDTSPRHLSNVSSTGSIDMV
 71                                  YKSPVVSGDTSPRHL
 72                                  YKSPVVSGDTSPRHLSNV
 73                                      VSGDTSPRHL
 74                                      VSGDTSPRHLSNV
 75                                      VSGDTSPRHLSNVSSTGSIDMV
 76                                       SGDTSPRHL
 77                                        GDTSPRHLSNVSSTGSIDMV
 78                                                SNVSSTGSIDM
 79                                                SNVSSTGSIDMV
 80                                                SNVSSTGSIDMV
 81                                                SNVSSTGSIDMV
 82                                                SNVSSTGSIDMV
 83                                                SNVSSTGSIDMV
 84                                                SNVSSTGSIDMV
 85                                                SNVSSTGSIDMV
 86                                                SNVSSTGSIDMV
 87                                                SNVSSTGSIDMV
 88                                                SNVSSTGSIDMV
 89                                                SNVSSTGSIDMV
 90                                                SNVSSTGSIDMV
 91                                                SNVSSTGSIDMV
 92                                                SNVSSTGSIDMV
 93                                                   SSTGSIDMV
 94                                                   SSTGSIDMV
 95                                                   SSTGSIDMV
 96                                                   SSTGSIDMV
 97                                                   SSTGSIDMV
 98                                                   SSTGSIDMV
 99                                                      GSIDMV
100                                                      GSIDMV

  1 DSPQLATLADEVSASLAKQGL
 66 D
 67 DSPQ
 68 DSPQL
 69 DSPQLA
 70 DSPQLATL
 75 DSPQLATL
 77 DSPQLATL
```

```
 80 D
 81 DS
 82 DSP
 83 DSPQ
 84 DSPQL
 85 DSPQLA
 86 DSPQLAT
 87 DSPQLATL
 88 DSPQLATLAD
 89 DSPQLATLADEV
 90 DSPQLATLADEVSA
 91 DSPQLATLADEVSAS
 92 DSPQLATLADEVSASLAKQGL
 93 DSPQ
 94 DSPQL
 95 DSPQLA
 96 DSPQLATL
 97 DSPQLATLADEV
 98 DSPQLATLADEVSASLAKQGL
 99 DSPQLATL
100 DSPQLATLADEVSASLAKQGL
101 DSPQLATLADEVSASLAKQGL
102     LATLADEVSASLAKQGL
103      ATLADEVSASLAKQGL
104       TLADEVSASLAKQGL
105        LADEVSASLAKQGL
106         ADEVSASLAK
107         ADEVSASLAKQG
108         ADEVSASLAKQGL
109             SASLAKQGL
110              ASLAKQGL
```

**Tau fibrils 2 min**

```
 1 MAEPRQEFEVMEDHAGTYGLGDRKDQGGYTMHQDQEGDTDAGLKESPLQTPTEDGSEEPG

 1 SETSDAKSTPTAEDVTAPLVDEGAPGKQAAAQPHTEIPEGTTAEEAGIGDTPSLEDEAAG

 1 HVTQARMVSKSKDGTGSDDKKAKGADGKTKIATPRGAAPPGQKGQANATRIPAKTPPAPK

 1 TPPSSGEPPKSGDRSGYSSPGSPGTPGSRSRTPSLPTPPTREPKKVAVVRTPPKSPSSAK
 2                                               AVVRTPPKSPSSAK
 3                                               AVVRTPPKSPSSAK
 4                                                  RTPPKSPSSAK
 5                                                          SAK
 6                                                            K

 1 SRLQTAPVPMPDLKNVKSKIGSTENLKHQPGGGKVQIINKKLDLSNVQSKCGSKDNIKHV
 2 SRLQ
 3 SRLQTAPVPMPDLKNV
 4 SRLQTAPVPMPDLKNV
 5 SRLQTAPVPMPDLKNV
 6 SRLQTAPVPMPDLKNV
 7  RLQTAPVPMPDLKNV
 8   LQTAPVPMPDLKNV
 9    QTAPVPMPDLKNV
10     TAPVPMPDLKNV
11      APVPMPDLKNV
12                 KSKIGSTENL
13                           KHQPGGGKVQI
14                           KHQPGGGKVQII
15                                      INKKLDLSNV
16                                       NKKLDLSNV
17                                                    GSKDNIKHV

 1 PGGGSVQIVYKPVDLSKVTSKCGSLGNIHHKPGGGQVEVKSEKLDFKDRVQSKIGSLDNI
17 PGGGSVQI
18         VYKPVDLSK
19         VYKPVDLSKV
20         VYKPVDLSKVT
21         VYKPVDLSKVTSK
22         VYKPVDLSKVTSKC
23          YKPVDLSKV
24                                             DFKDRVQSKIGSLDNI
25                                                   QSKIGSLDNI
26                                                    SKIGSLDNI
27                                                          DNI

 1 THVPGGGNKKIETHKLTFRENAKAKTDHGAEIVYKSPVVSGDTSPRHLSNVSSTGSIDMV
27 THVPGGGNKKIETHKL
28 THVPGGGN
29 THVPGGGNKKIETHKL
30 THVPGGGNKKIETHKLT
31    PGGGNKKIETHKL
32      GGNKKIETHKL
33       GNKKIETHKL
34                 TFRENAKA
35                 TFRENAKAKTD
36                 TFRENAKAKTDH
37                 TFRENAKAKTDHGAE
38                 TFRENAKAKTDHGAEI
39                 TFRENAKAKTDHGAEIV
40                 TFRENAKAKTDHGAEIVY
41                 TFRENAKAKTDHGAEIVYK
42                 TFRENAKAKTDHGAEIVYKSPV
43                 TFRENAKAKTDHGAEIVYKSPVVS
```

| | |
|---|---|
| 44 | TFRENAKAKTDHGAEIVYKSPVVSGDTSPRHL |
| 45 | TFRENAKAKTDHGAEIVYKSPVVSGDTSPRHLSNV |
| 46 | VYKSPVVS |
| 47 | VYKSPVVSG |
| 48 | VYKSPVVSGD |
| 49 | VYKSPVVSGDT |
| 50 | VYKSPVVSGDTS |
| 51 | VYKSPVVSGDTSP |
| 52 | VYKSPVVSGDTSPR |
| 53 | VYKSPVVSGDTSPRH |
| 54 | VYKSPVVSGDTSPRHL |
| 55 | VYKSPVVSGDTSPRHLS |
| 56 | VYKSPVVSGDTSPRHLSN |
| 57 | VYKSPVVSGDTSPRHLSNV |
| 58 | VYKSPVVSGDTSPRHLSNVS |
| 59 | VYKSPVVSGDTSPRHLSNVSS |
| 60 | VYKSPVVSGDTSPRHLSNVSST |
| 61 | VYKSPVVSGDTSPRHLSNVSSTG |
| 62 | VYKSPVVSGDTSPRHLSNVSSTGSID |
| 63 | VYKSPVVSGDTSPRHLSNVSSTGSIDM |
| 64 | VYKSPVVSGDTSPRHLSNVSSTGSIDMV |
| 65 | VYKSPVVSGDTSPRHLSNVSSTGSIDMV |
| 66 | VYKSPVVSGDTSPRHLSNVSSTGSIDMV |
| 67 | VYKSPVVSGDTSPRHLSNVSSTGSIDMV |
| 68 | VYKSPVVSGDTSPRHLSNVSSTGSIDMV |
| 69 | VYKSPVVSGDTSPRHLSNVSSTGSIDMV |
| 70 | VYKSPVVSGDTSPRHLSNVSSTGSIDMV |
| 71 | YKSPVVSGDTSPRHL |
| 72 | YKSPVVSGDTSPRHLSNV |
| 73 | VSGDTSPRHL |
| 74 | VSGDTSPRHLSNV |
| 75 | VSGDTSPRHLSNVSSTGSIDMV |
| 76 | VSGDTSPRHLSNVSSTGSIDMV |
| 77 | SGDTSPRHL |
| 78 | GDTSPRHLSNVSSTGSIDMV |
| 79 | SNVSSTGSIDM |
| 80 | SNVSSTGSIDMV |
| 81 | SNVSSTGSIDMV |
| 82 | SNVSSTGSIDMV |
| 83 | SNVSSTGSIDMV |
| 84 | SNVSSTGSIDMV |
| 85 | SNVSSTGSIDMV |
| 86 | SNVSSTGSIDMV |
| 87 | SNVSSTGSIDMV |
| 88 | SNVSSTGSIDMV |
| 89 | SNVSSTGSIDMV |
| 90 | SNVSSTGSIDMV |
| 91 | SNVSSTGSIDMV |
| 92 | SNVSSTGSIDMV |
| 93 | SNVSSTGSIDMV |
| 94 | SNVSSTGSIDMV |
| 95 | SNVSSTGSIDMV |
| 96 | SSTGSIDMV |
| 97 | SSTGSIDMV |
| 98 | SSTGSIDMV |
| 99 | SSTGSIDMV |
| 100 | SSTGSIDMV |
| 101 | SSTGSIDMV |
| 102 | SSTGSIDMV |
| 103 | SSTGSIDMV |
| 104 | SSTGSIDMV |
| 105 | GSIDMV |
| 106 | GSIDMV |
| 107 | GSIDMV |

```
108                                                       GSIDMV
109                                                           MV
110                                                            V

  1 DSPQLATLADEVSASLAKQGL
 65 D
 66 DSP
 67 DSPQ
 68 DSPQL
 69 DSPQLA
 70 DSPQLATL
 75 DSPQL
 76 DSPQLATL
 78 DSPQLATL
 81 D
 82 DS
 83 DSP
 84 DSPQ
 85 DSPQL
 86 DSPQLA
 87 DSPQLAT
 88 DSPQLATL
 89 DSPQLATLA
 90 DSPQLATLAD
 91 DSPQLATLADE
 92 DSPQLATLADEV
 93 DSPQLATLADEVSA
 94 DSPQLATLADEVSAS
 95 DSPQLATLADEVSASLAKQGL
 96 DSPQ
 97 DSPQL
 98 DSPQLA
 99 DSPQLAT
100 DSPQLATL
101 DSPQLATLA
102 DSPQLATLADEV
103 DSPQLATLADEVSA
104 DSPQLATLADEVSASLAKQGL
105 DSPQLA
106 DSPQLATL
107 DSPQLATLADEV
108 DSPQLATLADEVSASLAKQGL
109 DSPQLATL
110 DSPQLATLADEV
111 DSPQLATLADEV
112 DSPQLATLADEVSASLAKQGL
113     LATLADEVSASLAKQGL
114      ATLADEVSASLAKQGL
115       TLADEVSASLAKQGL
116        LADEVSASLAKQGL
117         ADEVSASLAK
118         ADEVSASLAKQG
119         ADEVSASLAKQGL
120             SASLAKQGL
121              ASLAKQGL
```

**Tau fibrils 5 min**

```
 1 MAEPRQEFEVMEDHAGTYGLGDRKDQGGYTMHQDQEGDTDAGLKESPLQTPTEDGSEEPG
 2  AEPRQEFEVMEDHAGTYGLGDRKDQGGYTMH

 1 SETSDAKSTPTAEDVTAPLVDEGAPGKQAAAQPHTEIPEGTTAEEAGIGDTPSLEDEAAG
 3                                               GIGDTPSLEDEAAG

 1 HVTQARMVSKSKDGTGSDDKKAKGADGKTKIATPRGAAPPGQKGQANATRIPAKTPPAPK
 3 HVTQA

 1 TPPSSGEPPKSGDRSGYSSPGSPGTPGSRSRTPSLPTPPTREPKKVAVVRTPPKSPSSAK
 4                                               AVVRTPPKSPS
 5                                               AVVRTPPKSPSSAK
 6                                               AVVRTPPKSPSSAK
 7                                               AVVRTPPKSPSSAK
 8                                                 VRTPPKSPSSAK
 9                                                  RTPPKSPSSAK
10                                                       SPSSAK
11                                                         SSAK
12                                                          SAK
13                                                            K

 1 SRLQTAPVPMPDLKNVKSKIGSTENLKHQPGGGKVQIINKKLDLSNVQSKCGSKDNIKHV
 5 S
 6 SRLQ
 7 SRLQTAPVPMPDLKNV
 8 SRLQTAPVPMPDLKNV
 9 SRLQTAPVPMPDLKNV
10 SRLQTAPVPMPDLKNV
11 SRLQTAPVPMPDLKNV
12 SRLQTAPVPMPDLKNV
13 SRLQTAPVPMPDLKNV
14  RLQTAPVPMPDLKNV
15  RLQTAPVPMPDLKNVKSKIGSTENL
16   LQTAPVPMPDLKNV
17    QTAPVPMPDLKNV
18     TAPVPMPDLKNV
19     TAPVPMPDLKNVKSKIGSTENL
20      APVPMPDLKNV
21                 KSKIGSTENL
22                 KSKIGSTENLKHQPGGGKVQI
23                 KSKIGSTENLKHQPGGGKVQII
24                           KHQPGGGKVQI
25                           KHQPGGGKVQII
26                           KHQPGGGKVQIINKKLDLS
27                           KHQPGGGKVQIINKKLDLSNV
28                                    QIINKKLDLSNV
29                                     IINKKLDLSNV
30                                      INKKLDLSNV
31                                       NKKLDLSNV
32                                                    GSKDNIKHV
33                                                          KHV
34                                                          KHV

 1 PGGGSVQIVYKPVDLSKVTSKCGSLGNIHHKPGGGQVEVKSEKLDFKDRVQSKIGSLDNI
32 PGGGSVQI
33 PGGGSVQI
34 PGGGSVQIVYKPVDLSKV
35         VYKPVDL
36         VYKPVDLS
37         VYKPVDLSK
38         VYKPVDLSKV
39         VYKPVDLSKVT
```

```
40         VYKPVDLSKVTS
41         VYKPVDLSKVTSK
42         VYKPVDLSKVTSKC
43          YKPVDLSKV
44                       GSLGNIHHKPGGGQVEV
45                       GSLGNIHHKPGGGQVEVKSEKL
46                             HHKPGGGQVEVKSEKL
47                                           KLDFKDRVQSKIGSLDNI
48                                             DFKDRVQSK
49                                             DFKDRVQSKIGSL
50                                             DFKDRVQSKIGSLDN
51                                             DFKDRVQSKIGSLDNI
52                                             DFKDRVQSKIGSLDNI
53                                             DFKDRVQSKIGSLDNI
54                                             DFKDRVQSKIGSLDNI
55                                             DFKDRVQSKIGSLDNI
56                                             DFKDRVQSKIGSLDNI
57                                                   QSKIGSLDNI
58                                                   QSKIGSLDNI
59                                                   QSKIGSLDNI
60                                                    SKIGSLDNI
61                                                       GSLDNI
62                                                        SLDNI
63                                                          DNI
64                                                          DNI

 1 THVPGGGNKKIETHKLTFRENAKAKTDHGAEIVYKSPVVSGDTSPRHLSNVSSTGSIDMV
52 T
53 TH
54 THVPG
55 THVPGGGN
56 THVPGGGNKKIETHKL
58 T
59 THVPGGGNKKIETHKL
61 THVPGGGNKKIETHKL
62 THVPGGGNKKIETHKL
63 THVPGGGNKKIETHKL
64 THVPGGGNKKIETHKLT
65 THVPGGGN
66 THVPGGGNKKIETH
67 THVPGGGNKKIETHK
68 THVPGGGNKKIETHKL
69 THVPGGGNKKIETHKLT
70    PGGGNKKIETHKL
71     GGGNKKIETHKL
72      GGNKKIETHKL
73                 TFRENAKAKTD
74                 TFRENAKAKTDH
75                 TFRENAKAKTDHGA
76                 TFRENAKAKTDHGAE
77                 TFRENAKAKTDHGAEI
78                 TFRENAKAKTDHGAEIV
79                 TFRENAKAKTDHGAEIVY
80                 TFRENAKAKTDHGAEIVYK
81                 TFRENAKAKTDHGAEIVYKS
82                 TFRENAKAKTDHGAEIVYKSPV
83                 TFRENAKAKTDHGAEIVYKSPVV
84                 TFRENAKAKTDHGAEIVYKSPVVS
85                 TFRENAKAKTDHGAEIVYKSPVVSG
86                 TFRENAKAKTDHGAEIVYKSPVVSGD
87                 TFRENAKAKTDHGAEIVYKSPVVSGDT
88                 TFRENAKAKTDHGAEIVYKSPVVSGDTS
89                 TFRENAKAKTDHGAEIVYKSPVVSGDTSP
90                 TFRENAKAKTDHGAEIVYKSPVVSGDTSPR
```

91 TFRENAKAKTDHGAEIVYKSPVVSGDTSPRH
92 TFRENAKAKTDHGAEIVYKSPVVSGDTSPRHL
93 TFRENAKAKTDHGAEIVYKSPVVSGDTSPRHLS
94 TFRENAKAKTDHGAEIVYKSPVVSGDTSPRHLSN
95 TFRENAKAKTDHGAEIVYKSPVVSGDTSPRHLSNV
96 TFRENAKAKTDHGAEIVYKSPVVSGDTSPRHLSNVS
97 KTDHGAEIVYKSPVVSGDTSPRHL
98 VYKSPVVSG
99 VYKSPVVSGD
100 VYKSPVVSGDT
101 VYKSPVVSGDTS
102 VYKSPVVSGDTSP
103 VYKSPVVSGDTSPR
104 VYKSPVVSGDTSPRH
105 VYKSPVVSGDTSPRHL
106 VYKSPVVSGDTSPRHLS
107 VYKSPVVSGDTSPRHLSN
108 VYKSPVVSGDTSPRHLSNV
109 VYKSPVVSGDTSPRHLSNVS
110 VYKSPVVSGDTSPRHLSNVSS
111 VYKSPVVSGDTSPRHLSNVSST
112 VYKSPVVSGDTSPRHLSNVSSTG
113 VYKSPVVSGDTSPRHLSNVSSTGS
114 VYKSPVVSGDTSPRHLSNVSSTGSID
115 VYKSPVVSGDTSPRHLSNVSSTGSIDM
116 VYKSPVVSGDTSPRHLSNVSSTGSIDMV
117 VYKSPVVSGDTSPRHLSNVSSTGSIDMV
118 VYKSPVVSGDTSPRHLSNVSSTGSIDMV
119 VYKSPVVSGDTSPRHLSNVSSTGSIDMV
120 VYKSPVVSGDTSPRHLSNVSSTGSIDMV
121 VYKSPVVSGDTSPRHLSNVSSTGSIDMV
122 VYKSPVVSGDTSPRHLSNVSSTGSIDMV
123 VYKSPVVSGDTSPRHLSNVSSTGSIDMV
124 VYKSPVVSGDTSPRHLSNVSSTGSIDMV
125 YKSPVVSGDTSPRHL
126 YKSPVVSGDTSPRHLSNV
127 YKSPVVSGDTSPRHLSNVSSTGSIDMV
128 KSPVVSGDTSPRHL
129 VSGDTSPRHL
130 VSGDTSPRHLSNV
131 VSGDTSPRHLSNVSSTGSIDMV
132 VSGDTSPRHLSNVSSTGSIDMV
133 VSGDTSPRHLSNVSSTGSIDMV
134 SGDTSPRHL
135 SGDTSPRHLSNV
136 GDTSPRHLSNVSSTGSIDMV
137 GDTSPRHLSNVSSTGSIDMV
138 SNVSSTGSIDM
139 SNVSSTGSIDMV
140 SNVSSTGSIDMV
141 SNVSSTGSIDMV
142 SNVSSTGSIDMV
143 SNVSSTGSIDMV
144 SNVSSTGSIDMV
145 SNVSSTGSIDMV
146 SNVSSTGSIDMV
147 SNVSSTGSIDMV
148 SNVSSTGSIDMV
149 SNVSSTGSIDMV
150 SNVSSTGSIDMV
151 SNVSSTGSIDMV
152 SNVSSTGSIDMV
153 SNVSSTGSIDMV
154 SNVSSTGSIDMV

155 SNVSSTGSIDMV
156 VSSTGSIDMV
157 SSTGSIDMV
158 SSTGSIDMV
159 SSTGSIDMV
160 SSTGSIDMV
161 SSTGSIDMV
162 SSTGSIDMV
163 SSTGSIDMV
164 SSTGSIDMV
165 SSTGSIDMV
166 SSTGSIDMV
167 GSIDMV
168 GSIDMV
169 GSIDMV
170 GSIDMV
171 MV
172 MV
173 V

1 DSPQLATLADEVSASLAKQGL
117 D
118 DS
119 DSP
120 DSPQ
121 DSPQL
122 DSPQLA
123 DSPQLAT
124 DSPQLATL
127 DSPQLATL
131 DSPQL
132 DSPQLATL
133 DSPQLATLADEV
136 DSPQLATL
137 DSPQLATLADEV
140 D
141 DS
142 DSP
143 DSPQ
144 DSPQL
145 DSPQLA
146 DSPQLAT
147 DSPQLATL
148 DSPQLATLA
149 DSPQLATLAD
150 DSPQLATLADE
151 DSPQLATLADEV
152 DSPQLATLADEVS
153 DSPQLATLADEVSA
154 DSPQLATLADEVSAS
155 DSPQLATLADEVSASLAKQGL
156 DSPQLATLADEVSASLAKQGL
157 DSPQ
158 DSPQL
159 DSPQLA
160 DSPQLAT
161 DSPQLATL
162 DSPQLATLA
163 DSPQLATLADEV
164 DSPQLATLADEVSA
165 DSPQLATLADEVSAS
166 DSPQLATLADEVSASLAKQGL
167 DSPQLA
168 DSPQLATL

169 DSPQLATLADEV
170 DSPQLATLADEVSASLAKQGL
171 DSPQLATL
172 DSPQLATLADEV
173 DSPQLATLADEV
174 DSPQLATLADEV
175 DSPQLATLADEVSASLAKQGL
176 LATLADEVSASLAKQGL
177 ATLADEVSASLAKQGL
178 TLADEVSASLAKQGL
179 LADEVSASLAKQGL
180 ADEVSASLAK
181 ADEVSASLAKQG
182 ADEVSASLAKQGL
183 SASLAKQGL
184 ASLAKQGL

**Tau fibrils 10 min**

```
 1 MAEPRQEFEVMEDHAGTYGLGDRKDQGGYTMHQDQEGDTDAGLKESPLQTPTEDGSEEPG
 2  AEPRQEFEVMEDHAGTYGL
 3  AEPRQEFEVMEDHAGTYGLGDRKDQGGYTMH

 1 SETSDAKSTPTAEDVTAPLVDEGAPGKQAAAQPHTEIPEGTTAEEAGIGDTPSLEDEAAG
 4                                               GIGDTPSLEDEAAG

 1 HVTQARMVSKSKDGTGSDDKKAKGADGKTKIATPRGAAPPGQKGQANATRIPAKTPPAPK
 4 HVTQA

 1 TPPSSGEPPKSGDRSGYSSPGSPGTPGSRSRTPSLPTPPTREPKKVAVVRTPPKSPSSAK
 5                                TPSLPTPPTREPKKV
 6                                               AVVRTPPKSPS
 7                                               AVVRTPPKSPSSAK
 8                                               AVVRTPPKSPSSAK
 9                                               AVVRTPPKSPSSAK
10                                                 VRTPPKSPSSAK
11                                                  RTPPKSPSSAK
12                                                       SPSSAK
13                                                         SSAK
14                                                          SAK
15                                                            K

 1 SRLQTAPVPMPDLKNVKSKIGSTENLKHQPGGGKVQIINKKLDLSNVQSKCGSKDNIKHV
 7 S
 8 SRLQ
 9 SRLQTAPVPMPDLKNV
10 SRLQTAPVPMPDLKNV
11 SRLQTAPVPMPDLKNV
12 SRLQTAPVPMPDLKNV
13 SRLQTAPVPMPDLKNV
14 SRLQTAPVPMPDLKNV
15 SRLQTAPVPMPDLKNV
16  RLQTAPVPMPDLKNV
17  RLQTAPVPMPDLKNVKSKIGSTENL
18   LQTAPVPMPDLKNV
19    QTAPVPMPDLKNV
20     TAPVPMPDLKNV
21     TAPVPMPDLKNVKSKIGSTENL
22      APVPMPDLKNV
23        VPMPDLKNV
24                 KSKIGSTENL
25                 KSKIGSTENLKHQPGGGKVQI
26                 KSKIGSTENLKHQPGGGKVQII
27                           KHQPGGGKVQI
28                           KHQPGGGKVQII
29                           KHQPGGGKVQIINKKLDLS
30                           KHQPGGGKVQIINKKLDLSNV
31                           KHQPGGGKVQIINKKLDLSNVQS
32                                    QIINKKLDLSNV
33                                     IINKKLDLSNV
34                                      INKKLDLSNV
35                                      INKKLDLSNVQS
36                                       NKKLDLSNV
37                                       NKKLDLSNVQSKC
38                                                   GSKDNIKHV
39                                                         KHV
40                                                         KHV

 1 PGGGSVQIVYKPVDLSKVTSKCGSLGNIHHKPGGGQVEVKSEKLDFKDRVQSKIGSLDNI
38 PGGGSVQI
39 PGGGSVQI
```

```
40 PGGGSVQIVYKPVDLSKV
41         VYKPVDL
42         VYKPVDLS
43         VYKPVDLSK
44         VYKPVDLSKV
45         VYKPVDLSKVT
46         VYKPVDLSKVTS
47         VYKPVDLSKVTSK
48         VYKPVDLSKVTSKC
49          YKPVDLSKV
50                       GSLGNIHHKPGGGQVEV
51                       GSLGNIHHKPGGGQVEVKSEKL
52                          GNIHHKPGGGQVEVKSEKL
53                             HHKPGGGQVEVKSEKL
54                                        KSEKLDFKDRVQSKIGSLDNI
55                                           KLDFKDRVQSKIGSLDNI
56                                             DFKDRVQSK
57                                             DFKDRVQSKIGS
58                                             DFKDRVQSKIGSL
59                                             DFKDRVQSKIGSLDN
60                                             DFKDRVQSKIGSLDNI
61                                             DFKDRVQSKIGSLDNI
62                                             DFKDRVQSKIGSLDNI
63                                             DFKDRVQSKIGSLDNI
64                                             DFKDRVQSKIGSLDNI
65                                             DFKDRVQSKIGSLDNI
66                                             DFKDRVQSKIGSLDNI
67                                             DFKDRVQSKIGSLDNI
68                                             DFKDRVQSKIGSLDNI
69                                                  VQSKIGSLDNI
70                                                   QSKIGSLDNI
71                                                   QSKIGSLDNI
72                                                   QSKIGSLDNI
73                                                    SKIGSLDNI
74                                                    SKIGSLDNI
75                                                     KIGSLDNI
76                                                       GSLDNI
77                                                        SLDNI
78                                                          DNI
79                                                          DNI

 1 THVPGGGNKKIETHKLTFRENAKAKTDHGAEIVYKSPVVSGDTSPRHLSNVSSTGSIDMV
61 T
62 TH
63 THVPG
64 THVPGG
65 THVPGGG
66 THVPGGGN
67 THVPGGGNKKIETHKL
68 THVPGGGNKKIETHKLT
71 T
72 THVPGGGNKKIETHKL
74 THVPGGGNKKIETHKL
75 THVPGGGNKKIETHKL
76 THVPGGGNKKIETHKL
77 THVPGGGNKKIETHKL
78 THVPGGGNKKIETHKL
79 THVPGGGNKKIETHKLT
80 THVPGGGNKKIETHK
81 THVPGGGNKKIETHKL
82 THVPGGGNKKIETHKLT
83 THVPGGGNKKIETHKLTFRENA
84    PGGGNKKIETHKL
85     GGGNKKIETHKL
```

| | |
|---|---|
| 86 | GGNKKIETHKL |
| 87 | GNKKIETHKL |
| 88 | TFRENAKAKTD |
| 89 | TFRENAKAKTDH |
| 90 | TFRENAKAKTDHGAE |
| 91 | TFRENAKAKTDHGAEI |
| 92 | TFRENAKAKTDHGAEIV |
| 93 | TFRENAKAKTDHGAEIVY |
| 94 | TFRENAKAKTDHGAEIVYK |
| 95 | TFRENAKAKTDHGAEIVYKS |
| 96 | TFRENAKAKTDHGAEIVYKSPV |
| 97 | TFRENAKAKTDHGAEIVYKSPVV |
| 98 | TFRENAKAKTDHGAEIVYKSPVVS |
| 99 | TFRENAKAKTDHGAEIVYKSPVVSG |
| 100 | TFRENAKAKTDHGAEIVYKSPVVSGD |
| 101 | TFRENAKAKTDHGAEIVYKSPVVSGDT |
| 102 | TFRENAKAKTDHGAEIVYKSPVVSGDTS |
| 103 | TFRENAKAKTDHGAEIVYKSPVVSGDTSP |
| 104 | TFRENAKAKTDHGAEIVYKSPVVSGDTSPR |
| 105 | TFRENAKAKTDHGAEIVYKSPVVSGDTSPRH |
| 106 | TFRENAKAKTDHGAEIVYKSPVVSGDTSPRHL |
| 107 | TFRENAKAKTDHGAEIVYKSPVVSGDTSPRHLS |
| 108 | TFRENAKAKTDHGAEIVYKSPVVSGDTSPRHLSN |
| 109 | TFRENAKAKTDHGAEIVYKSPVVSGDTSPRHLSNV |
| 110 | TFRENAKAKTDHGAEIVYKSPVVSGDTSPRHLSNVS |
| 111 | KTDHGAEIVYKSPVVSGDTSPRHL |
| 112 | VYKSPVVSG |
| 113 | VYKSPVVSGD |
| 114 | VYKSPVVSGDT |
| 115 | VYKSPVVSGDTS |
| 116 | VYKSPVVSGDTSP |
| 117 | VYKSPVVSGDTSPR |
| 118 | VYKSPVVSGDTSPRH |
| 119 | VYKSPVVSGDTSPRHL |
| 120 | VYKSPVVSGDTSPRHLS |
| 121 | VYKSPVVSGDTSPRHLSN |
| 122 | VYKSPVVSGDTSPRHLSNV |
| 123 | VYKSPVVSGDTSPRHLSNVS |
| 124 | VYKSPVVSGDTSPRHLSNVSST |
| 125 | VYKSPVVSGDTSPRHLSNVSSTG |
| 126 | VYKSPVVSGDTSPRHLSNVSSTGS |
| 127 | VYKSPVVSGDTSPRHLSNVSSTGSID |
| 128 | VYKSPVVSGDTSPRHLSNVSSTGSIDM |
| 129 | VYKSPVVSGDTSPRHLSNVSSTGSIDMV |
| 130 | VYKSPVVSGDTSPRHLSNVSSTGSIDMV |
| 131 | VYKSPVVSGDTSPRHLSNVSSTGSIDMV |
| 132 | VYKSPVVSGDTSPRHLSNVSSTGSIDMV |
| 133 | VYKSPVVSGDTSPRHLSNVSSTGSIDMV |
| 134 | VYKSPVVSGDTSPRHLSNVSSTGSIDMV |
| 135 | VYKSPVVSGDTSPRHLSNVSSTGSIDMV |
| 136 | VYKSPVVSGDTSPRHLSNVSSTGSIDMV |
| 137 | VYKSPVVSGDTSPRHLSNVSSTGSIDMV |
| 138 | YKSPVVSGDTSPRHL |
| 139 | YKSPVVSGDTSPRHLSNV |
| 140 | YKSPVVSGDTSPRHLSNVSSTGSIDMV |
| 141 | KSPVVSGDTSPRHL |
| 142 | VSGDTSPRHL |
| 143 | VSGDTSPRHLSNV |
| 144 | VSGDTSPRHLSNVSSTGSIDMV |
| 145 | VSGDTSPRHLSNVSSTGSIDMV |
| 146 | VSGDTSPRHLSNVSSTGSIDMV |
| 147 | VSGDTSPRHLSNVSSTGSIDMV |
| 148 | SGDTSPRHL |
| 149 | SGDTSPRHLSNV |

```
150                                         GDTSPRHLSNVSSTGSIDMV
151                                         GDTSPRHLSNVSSTGSIDMV
152                                         GDTSPRHLSNVSSTGSIDMV
153                                                 SNVSSTGSIDM
154                                                 SNVSSTGSIDMV
155                                                 SNVSSTGSIDMV
156                                                 SNVSSTGSIDMV
157                                                 SNVSSTGSIDMV
158                                                 SNVSSTGSIDMV
159                                                 SNVSSTGSIDMV
160                                                 SNVSSTGSIDMV
161                                                 SNVSSTGSIDMV
162                                                 SNVSSTGSIDMV
163                                                 SNVSSTGSIDMV
164                                                 SNVSSTGSIDMV
165                                                 SNVSSTGSIDMV
166                                                 SNVSSTGSIDMV
167                                                 SNVSSTGSIDMV
168                                                 SNVSSTGSIDMV
169                                                 SNVSSTGSIDMV
170                                                 SNVSSTGSIDMV
171                                                   VSSTGSIDMV
172                                                    SSTGSIDMV
173                                                    SSTGSIDMV
174                                                    SSTGSIDMV
175                                                    SSTGSIDMV
176                                                    SSTGSIDMV
177                                                    SSTGSIDMV
178                                                    SSTGSIDMV
179                                                    SSTGSIDMV
180                                                    SSTGSIDMV
181                                                       GSIDMV
182                                                       GSIDMV
183                                                       GSIDMV
184                                                       GSIDMV
185                                                           MV
186                                                           MV
187                                                            V

  1 DSPQLATLADEVSASLAKQGL
130 D
131 DS
132 DSP
133 DSPQ
134 DSPQL
135 DSPQLA
136 DSPQLAT
137 DSPQLATL
140 DSPQLATL
144 DSPQL
145 DSPQLA
146 DSPQLATL
147 DSPQLATLADEV
150 DSPQLA
151 DSPQLATL
152 DSPQLATLADEV
155 D
156 DS
157 DSP
158 DSPQ
159 DSPQL
160 DSPQLA
161 DSPQLAT
162 DSPQLATL
```

163 DSPQLATLA
164 DSPQLATLAD
165 DSPQLATLADE
166 DSPQLATLADEV
167 DSPQLATLADEVS
168 DSPQLATLADEVSA
169 DSPQLATLADEVSAS
170 DSPQLATLADEVSASLAKQGL
171 DSPQLATLADEVSASLAKQGL
172 DSPQ
173 DSPQL
174 DSPQLA
175 DSPQLAT
176 DSPQLATL
177 DSPQLATLADEV
178 DSPQLATLADEVSA
179 DSPQLATLADEVSAS
180 DSPQLATLADEVSASLAKQGL
181 DSPQLA
182 DSPQLATL
183 DSPQLATLADEV
184 DSPQLATLADEVSASLAKQGL
185 DSPQLATL
186 DSPQLATLADEV
187 DSPQLATLADEV
188 DSPQLATLADEV
189 DSPQLATLADEVSASLAKQGL
190     LATLADEVSASLAKQGL
191      ATLADEVSASLAKQGL
192       TLADEVSASLAKQGL
193        LADEVSASLAKQGL
194         ADEVSASLAK
195         ADEVSASLAKQG
196         ADEVSASLAKQGL
197             SASLAKQGL
198              ASLAKQGL

**Tau fibrils 120 min**

```
  1 MAEPRQEFEVMEDHAGTYGLGDRKDQGGYTMHQDQEGDTDAGLKESPLQTPTEDGSEEPG

  1 SETSDAKSTPTAEDVTAPLVDEGAPGKQAAAQPHTEIPEGTTAEEAGIGDTPSLEDEAAG
  2                                               GIGDTPSLEDEAAG
  3                                                     SLEDEAAG

  1 HVTQARMVSKSKDGTGSDDKKAKGADGKTKIATPRGAAPPGQKGQANATRIPAKTPPAPK
  2 HVTQA
  3 HVTQA

  1 TPPSSGEPPKSGDRSGYSSPGSPGTPGSRSRTPSLPTPPTREPKKVAVVRTPPKSPSSAK
  4                            SRSRTPSLPTPPTREPKKV
  5                                TPSLPTPPTREPKKV
  6                                               AVVRTPPKSPS
  7                                               AVVRTPPKSPSSA
  8                                               AVVRTPPKSPSSAK
  9                                               AVVRTPPKSPSSAK
 10                                               AVVRTPPKSPSSAK
 11                                               AVVRTPPKSPSSAK
 12                                                VVRTPPKSPSSAK
 13                                                 VRTPPKSPSSAK
 14                                                  RTPPKSPSSAK
 15                                                       SPSSAK
 16                                                         SSAK
 17                                                          SAK
 18                                                          SAK
 19                                                            K

  1 SRLQTAPVPMPDLKNVKSKIGSTENLKHQPGGGKVQIINKKLDLSNVQSKCGSKDNIKHV
  8 S
  9 SRLQ
 10 SRLQT
 11 SRLQTAPVPMPDLKNV
 12 SRLQTAPVPMPDLKNV
 13 SRLQTAPVPMPDLKNV
 14 SRLQTAPVPMPDLKNV
 15 SRLQTAPVPMPDLKNV
 16 SRLQTAPVPMPDLKNV
 17 SRLQTAPVPMPDLKNV
 18 SRLQTAPVPMPDLKNVKSKIGST
 19 SRLQTAPVPMPDLKNV
 20 SRLQTAPVPMPDLKNV
 21  RLQTAPVPMPDLKNV
 22  RLQTAPVPMPDLKNVKSKIGSTENL
 23   LQTAPVPMPDLKNV
 24    QTAPVPMPDLKNV
 25     TAPVPMPDLKNV
 26     TAPVPMPDLKNVKSKIGST
 27     TAPVPMPDLKNVKSKIGSTENL
 28      APVPMPDLKNV
 29       PVPMPDLKNV
 30        VPMPDLKNV
 31                 KSKIGSTENL
 32                 KSKIGSTENLKHQPGGGKVQI
 33                 KSKIGSTENLKHQPGGGKVQII
 34                        ENLKHQPGGGKVQI
 35                        ENLKHQPGGGKVQII
 36                           KHQPGGGKVQI
 37                           KHQPGGGKVQII
 38                           KHQPGGGKVQIINKKLDLS
 39                           KHQPGGGKVQIINKKLDLSNV
 40                           KHQPGGGKVQIINKKLDLSNVQS
```

```
41                                    QIINKKLDLSNV
42                                     IINKKLDLSNV
43                                      INKKLDLSNV
44                                      INKKLDLSNVQS
45                                       NKKLDLSNV
46                                       NKKLDLSNVQS
47                                       NKKLDLSNVQSKC
48                                                    GSKDNIKHV
49                                                          KHV
50                                                          KHV

 1 PGGGSVQIVYKPVDLSKVTSKCGSLGNIHHKPGGGQVEVKSEKLDFKDRVQSKIGSLDNI
48 PGGGSVQI
49 PGGGSVQI
50 PGGGSVQIVYKPVDLSKV
51         VYKPVDL
52         VYKPVDLS
53         VYKPVDLSK
54         VYKPVDLSKV
55         VYKPVDLSKVT
56         VYKPVDLSKVTS
57         VYKPVDLSKVTSK
58         VYKPVDLSKVTSKC
59          YKPVDLSKV
60                       GSLGNIHHKPGGGQVEV
61                       GSLGNIHHKPGGGQVEVKSEKL
62                          GNIHHKPGGGQVEVKSEKL
63                             HHKPGGGQVEVKSEKL
64                                        KSEKLDFKDRVQSKIGSLDNI
65                                           KLDFKDRVQSKIGSLDNI
66                                             DFKDRVQSKIGS
67                                             DFKDRVQSKIGSL
68                                             DFKDRVQSKIGSLD
69                                             DFKDRVQSKIGSLDN
70                                             DFKDRVQSKIGSLDNI
71                                             DFKDRVQSKIGSLDNI
72                                             DFKDRVQSKIGSLDNI
73                                             DFKDRVQSKIGSLDNI
74                                             DFKDRVQSKIGSLDNI
75                                             DFKDRVQSKIGSLDNI
76                                             DFKDRVQSKIGSLDNI
77                                                   QSKIGSLDNI
78                                                   QSKIGSLDNI
79                                                   QSKIGSLDNI
80                                                   QSKIGSLDNI
81                                                    SKIGSLDNI
82                                                    SKIGSLDNI
83                                                     KIGSLDNI
84                                                       GSLDNI
85                                                        SLDNI
86                                                          DNI
87                                                          DNI

 1 THVPGGGNKKIETHKLTFRENAKAKTDHGAEIVYKSPVVSGDTSPRHLSNVSSTGSIDMV
71 T
72 TH
73 THV
74 THVPG
75 THVPGGGN
76 THVPGGGNKKIETHKL
78 T
79 TH
80 THVPGGGNKKIETHKL
82 THVPGGGNKKIETHKL
```

```
 83 THVPGGGNKKIETHKL
 84 THVPGGGNKKIETHKL
 85 THVPGGGNKKIETHKL
 86 THVPGGGNKKIETHKL
 87 THVPGGGNKKIETHKLT
 88 THVPGGGN
 89 THVPGGGNKKIETH
 90 THVPGGGNKKIETHK
 91 THVPGGGNKKIETHKL
 92 THVPGGGNKKIETHKLT
 93    PGGGNKKIETHKL
 94     GGGNKKIETHKL
 95      GGNKKIETHKL
 96       GNKKIETHKL
 97                 TFRENAKAKTD
 98                 TFRENAKAKTDH
 99                 TFRENAKAKTDHGA
100                 TFRENAKAKTDHGAE
101                 TFRENAKAKTDHGAEI
102                 TFRENAKAKTDHGAEIV
103                 TFRENAKAKTDHGAEIVY
104                 TFRENAKAKTDHGAEIVYK
105                 TFRENAKAKTDHGAEIVYKS
106                 TFRENAKAKTDHGAEIVYKSPV
107                 TFRENAKAKTDHGAEIVYKSPVV
108                 TFRENAKAKTDHGAEIVYKSPVVS
109                 TFRENAKAKTDHGAEIVYKSPVVSG
110                 TFRENAKAKTDHGAEIVYKSPVVSGD
111                 TFRENAKAKTDHGAEIVYKSPVVSGDT
112                 TFRENAKAKTDHGAEIVYKSPVVSGDTS
113                 TFRENAKAKTDHGAEIVYKSPVVSGDTSP
114                 TFRENAKAKTDHGAEIVYKSPVVSGDTSPR
115                 TFRENAKAKTDHGAEIVYKSPVVSGDTSPRH
116                 TFRENAKAKTDHGAEIVYKSPVVSGDTSPRHL
117                 TFRENAKAKTDHGAEIVYKSPVVSGDTSPRHLS
118                 TFRENAKAKTDHGAEIVYKSPVVSGDTSPRHLSN
119                 TFRENAKAKTDHGAEIVYKSPVVSGDTSPRHLSNV
120                 TFRENAKAKTDHGAEIVYKSPVVSGDTSPRHLSNVS
121                         KTDHGAEIVYKSPVVSGDTSPRHL
122                                 VYKSPVVS
123                                 VYKSPVVSG
124                                 VYKSPVVSGD
125                                 VYKSPVVSGDT
126                                 VYKSPVVSGDTS
127                                 VYKSPVVSGDTSP
128                                 VYKSPVVSGDTSPR
129                                 VYKSPVVSGDTSPRH
130                                 VYKSPVVSGDTSPRHL
131                                 VYKSPVVSGDTSPRHLS
132                                 VYKSPVVSGDTSPRHLSN
133                                 VYKSPVVSGDTSPRHLSNV
134                                 VYKSPVVSGDTSPRHLSNVS
135                                 VYKSPVVSGDTSPRHLSNVSS
136                                 VYKSPVVSGDTSPRHLSNVSST
137                                 VYKSPVVSGDTSPRHLSNVSSTG
138                                 VYKSPVVSGDTSPRHLSNVSSTGSID
139                                 VYKSPVVSGDTSPRHLSNVSSTGSIDM
140                                 VYKSPVVSGDTSPRHLSNVSSTGSIDMV
141                                 VYKSPVVSGDTSPRHLSNVSSTGSIDMV
142                                 VYKSPVVSGDTSPRHLSNVSSTGSIDMV
143                                 VYKSPVVSGDTSPRHLSNVSSTGSIDMV
144                                 VYKSPVVSGDTSPRHLSNVSSTGSIDMV
145                                 VYKSPVVSGDTSPRHLSNVSSTGSIDMV
146                                 VYKSPVVSGDTSPRHLSNVSSTGSIDMV
```

```
147                                 VYKSPVVSGDTSPRHLSNVSSTGSIDMV
148                                 VYKSPVVSGDTSPRHLSNVSSTGSIDMV
149                                  YKSPVVSGDTSPRHL
150                                  YKSPVVSGDTSPRHLSNV
151                                  YKSPVVSGDTSPRHLSNVSSTGSIDMV
152                                   KSPVVSGDTSPRHL
153                                       VSGDTSPRHL
154                                       VSGDTSPRHLSNV
155                                       VSGDTSPRHLSNVSSTGSIDMV
156                                       VSGDTSPRHLSNVSSTGSIDMV
157                                       VSGDTSPRHLSNVSSTGSIDMV
158                                       VSGDTSPRHLSNVSSTGSIDMV
159                                        SGDTSPRHL
160                                        SGDTSPRHLSNV
161                                         GDTSPRHLSNVSSTGSIDMV
162                                         GDTSPRHLSNVSSTGSIDMV
163                                         GDTSPRHLSNVSSTGSIDMV
164                                                 SNVSSTGSIDM
165                                                 SNVSSTGSIDMV
166                                                 SNVSSTGSIDMV
167                                                 SNVSSTGSIDMV
168                                                 SNVSSTGSIDMV
169                                                 SNVSSTGSIDMV
170                                                 SNVSSTGSIDMV
171                                                 SNVSSTGSIDMV
172                                                 SNVSSTGSIDMV
173                                                 SNVSSTGSIDMV
174                                                 SNVSSTGSIDMV
175                                                 SNVSSTGSIDMV
176                                                 SNVSSTGSIDMV
177                                                 SNVSSTGSIDMV
178                                                 SNVSSTGSIDMV
179                                                 SNVSSTGSIDMV
180                                                 SNVSSTGSIDMV
181                                                 SNVSSTGSIDMV
182                                                    SSTGSIDMV
183                                                    SSTGSIDMV
184                                                    SSTGSIDMV
185                                                    SSTGSIDMV
186                                                    SSTGSIDMV
187                                                    SSTGSIDMV
188                                                    SSTGSIDMV
189                                                    SSTGSIDMV
190                                                       GSIDMV
191                                                       GSIDMV
192                                                       GSIDMV
193                                                       GSIDMV
194                                                           MV
195                                                            V

  1 DSPQLATLADEVSASLAKQGL
141 D
142 DS
143 DSP
144 DSPQ
145 DSPQL
146 DSPQLA
147 DSPQLAT
148 DSPQLATL
151 DSPQLATL
155 DSPQL
156 DSPQLA
157 DSPQLATL
158 DSPQLATLADEV
```

```
161 DSPQLA
162 DSPQLATL
163 DSPQLATLADEV
166 D
167 DS
168 DSP
169 DSPQ
170 DSPQL
171 DSPQLA
172 DSPQLAT
173 DSPQLATL
174 DSPQLATLA
175 DSPQLATLAD
176 DSPQLATLADE
177 DSPQLATLADEV
178 DSPQLATLADEVS
179 DSPQLATLADEVSA
180 DSPQLATLADEVSAS
181 DSPQLATLADEVSASLAKQGL
183 DSPQL
184 DSPQLA
185 DSPQLATL
186 DSPQLATLADEV
187 DSPQLATLADEVSA
188 DSPQLATLADEVSAS
189 DSPQLATLADEVSASLAKQGL
190 DSPQLA
191 DSPQLATL
192 DSPQLATLADEV
193 DSPQLATLADEVSASLAKQGL
194 DSPQLATLADEV
195 DSPQLATLADEV
196 DSPQLATLADEV
197 DSPQLATLADEVSASLAKQGL
198     LATLADEVSASLAKQGL
199      ATLADEVSASLAKQGL
200       TLADEVSASLAKQGL
201        LADEVSASLAKQGL
202         ADEVSASLAK
203         ADEVSASLAKQG
204         ADEVSASLAKQGL
205             SASLAKQGL
206              ASLAKQGL
```

**Supplemental data 4**

**Tau CN (β-sheets highlighted in yellow, N-and C-terminal swaps highlighted in grey)**

MAEPRQEFEVMEDHAGTYGLGDRKDQGGYTMHQDQEGDTDAEIVYKSPVVSGDTSPRHLSNVSST
GSIDMVDSPQLATLADEVSASLAKQGLEGTTAEEAGIGDTPSLEDEAAGHVTQARMVSKSKDGTG
SDDKKAKGADGKTKIATPRGAAPPGQKGQANATRIPAKTPPAPKTPPSSGEPPKSGDRSGYSSPG
SPGTPGSRSRTPSLPTPPTREPKKVAVVRTPPKSPSSAKSRLQTAPVPMPDLKNVKSKIGSTENL
KHQPGGGKVQIINKKLDLSNVQSKCGSKDNIKHVPGGGSVQIVYKPVDLSKVTSKCGSLGNIHHK
PGGGQVEVKSEKLDFKDRVQSKIGSLDNITHVPGGGNKKIETHKLTFRENAKAKTDHGAGLKESP
LQTPTEDGSEEPGSETSDAKSTPTAEDVTAPLVDEGAPGKQAAAQPHTEIP

**Tau CN fibrils 0 sec**

```
  1 MAEPRQEFEVMEDHAGTYGLGDRKDQGGYTMHQDQEGDTDAEIVYKSPVVSGDTSPRHLS

  1 NVSSTGSIDMVDSPQLATLADEVSASLAKQGLEGTTAEEAGIGDTPSLEDEAAGHVTQAR
  2                        SASLAKQGL
  3                        SASLAKQGLEGTTAEEA
  4                        SASLAKQGLEGTTAEEAGIGDTPSLEDEAAGHVTQA

  1 MVSKSKDGTGSDDKKAKGADGKTKIATPRGAAPPGQKGQANATRIPAKTPPAPKTPPSSG

  1 EPPKSGDRSGYSSPGSPGTPGSRSRTPSLPTPPTREPKKVAVVRTPPKSPSSAKSRLQTA
  5                                                         LQTA

  1 PVPMPDLKNVKSKIGSTENLKHQPGGGKVQIINKKLDLSNVQSKCGSKDNIKHVPGGGSV
  5 PVPMPDLKNVKSKIG

  1 QIVYKPVDLSKVTSKCGSLGNIHHKPGGGQVEVKSEKLDFKDRVQSKIGSLDNITHVPGG

  1 GNKKIETHKLTFRENAKAKTDHGAGLKESPLQTPTEDGSEEPGSETSDAKSTPTAEDVTA

  1 PLVDEGAPGKQAAAQPHTEIP
```

**Tau CN fibrils 15 sec**

```
 1 MAEPRQEFEVMEDHAGTYGLGDRKDQGGYTMHQDQEGDTDAEIVYKSPVVSGDTSPRHLS
 2  AEPRQEFEV
 3                                            VYKSPVVSGDTSPRH
 4                                            VYKSPVVSGDTSPRHL
 5                                            VYKSPVVSGDTSPRHLS
 6                                            VYKSPVVSGDTSPRHLS
 7                                                              S
 8                                                              S
 9                                                              S

 1 NVSSTGSIDMVDSPQLATLADEVSASLAKQGLEGTTAEEAGIGDTPSLEDEAAGHVTQAR
 6 NV
 7 NVSSTGSIDMVDSPQLATL
 8 NVSSTGSIDMVDSPQLATLADEV
 9 NVSSTGSIDMVDSPQLATLADEVSASLAKQGL
10   SSTGSIDMVDSPQLATL
11   SSTGSIDMVDSPQLATLADEV
12                    ADEVSASLAKQGL
13                        SASLAKQGL
14                        SASLAKQGLEGTTAEEA
15                        SASLAKQGLEGTTAEEAGIGDTPSLEDEAAGHVTQA

 1 MVSKSKDGTGSDDKKAKGADGKTKIATPRGAAPPGQKGQANATRIPAKTPPAPKTPPSSG

 1 EPPKSGDRSGYSSPGSPGTPGSRSRTPSLPTPPTREPKKVAVVRTPPKSPSSAKSRLQTA
16                                         AVVRTPPKSPSSAKSRLQTA
17                                                    SAKSRLQTA
18                                                         LQTA

 1 PVPMPDLKNVKSKIGSTENLKHQPGGGKVQIINKKLDLSNVQSKCGSKDNIKHVPGGGSV
16 PVPMPDLKNV
17 PVPMPDLKNV
18 PVPMPDLKNVKSKIG
19                               IINKKLDLSNV
20                                              GSKDNIKHVPGGGSV

 1 QIVYKPVDLSKVTSKCGSLGNIHHKPGGGQVEVKSEKLDFKDRVQSKIGSLDNITHVPGG
20 QI
21  IVYKPVDLSKV
22   VYKPVDLSKV
23   VYKPVDLSKVTS
24                                        DFKDRVQSKIGSLDNI
25                                                       THVPGG

 1 GNKKIETHKLTFRENAKAKTDHGAGLKESPLQTPTEDGSEEPGSETSDAKSTPTAEDVTA
25 GNKKIETH
26           TFRENAKAKTD

 1 PLVDEGAPGKQAAAQPHTEIP
```

**Tau CN fibrils 1 min**

```
 1 MAEPRQEFEVMEDHAGTYGLGDRKDQGGYTMHQDQEGDTDAEIVYKSPVVSGDTSPRHLS
 2  AEPRQEFEV
 3                                            VYKSPVVSGDTSPR
 4                                            VYKSPVVSGDTSPRH
 5                                            VYKSPVVSGDTSPRHL
 6                                            VYKSPVVSGDTSPRHLS
 7                                            VYKSPVVSGDTSPRHLS
 8                                            VYKSPVVSGDTSPRHLS
 9                                            VYKSPVVSGDTSPRHLS
10                                             YKSPVVSGDTSPRHL
11                                                  VSGDTSPRHL
12                                                   SGDTSPRHL
13                                                            S
14                                                            S
15                                                            S

 1 NVSSTGSIDMVDSPQLATLADEVSASLAKQGLEGTTAEEAGIGDTPSLEDEAAGHVTQAR
 7 N
 8 NV
 9 NVS
13 NVSSTGSIDMVDSPQL
14 NVSSTGSIDMVDSPQLATL
15 NVSSTGSIDMVDSPQLATLADEVSASLAKQGL
16   SSTGSIDMVDSPQLATL
17   SSTGSIDMVDSPQLATLADEVSASLAKQGL
18            DSPQLATLADEVSASLAKQGL
19                  TLADEVSASLAKQGL
20                   LADEVSASLAKQGL
21                    ADEVSASLAKQ
22                    ADEVSASLAKQGL
23                        SASLAKQGL
24                        SASLAKQGLEGTTAEEAGIGDTPSLEDEAAGHVTQA

 1 MVSKSKDGTGSDDKKAKGADGKTKIATPRGAAPPGQKGQANATRIPAKTPPAPKTPPSSG

 1 EPPKSGDRSGYSSPGSPGTPGSRSRTPSLPTPPTREPKKVAVVRTPPKSPSSAKSRLQTA
25                                         AVVRTPPKSPSSAKSRLQTA
26                                          VVRTPPKSPSSAKSRLQTA
27                                            RTPPKSPSSAKSRLQTA
28                                                    SAKSRLQTA
29                                                       SRLQTA
30                                                         LQTA
31                                                         LQTA
32                                                           TA

 1 PVPMPDLKNVKSKIGSTENLKHQPGGGKVQIINKKLDLSNVQSKCGSKDNIKHVPGGGSV
25 PVPMPDLKNV
26 PVPMPDLKNV
27 PVPMPDLKNV
28 PVPMPDLKNV
29 PVPMPDLKNV
30 PVPMPDLKNV
31 PVPMPDLKNVKSKIG
32 PVPMPDLKNVKSKIGSTENL
33           KSKIGSTENL
34            SKIGSTENL
35                     KHQPGGGKVQI
36                               IINKKLDLSNV
37                                INKKLDLSNV
38                                 NKKLDLSNV
39                                              GSKDNIKHVPGGGSV
40                                               SKDNIKHVPGGGSV
```

```
 1 QIVYKPVDLSKVTSKCGSLGNIHHKPGGGQVEVKSEKLDFKDRVQSKIGSLDNITHVPGG
39 QI
40 QI
41  IVYKPVDLSKV
42   VYKPVDL
43   VYKPVDLSKV
44   VYKPVDLSKVT
45   VYKPVDLSKVTS
46   VYKPVDLSKVTSK
47                 GSLGNIHHKPGGGQ
48                 GSLGNIHHKPGGGQVEV
49                                       DFKDRVQSKIGSL
50                                       DFKDRVQSKIGSLDNI
51                                             QSKIGSLDNI
52                                                    DNITHVPGG
53                                                       THVPGG
54                                                       THVPGG
55                                                       THVPGG
56                                                       THVPGG
57                                                       THVPGG
58                                                         VPGG
59                                                          PGG
60                                                           GG
61                                                            G

 1 GNKKIETHKLTFRENAKAKTDHGAGLKESPLQTPTEDGSEEPGSETSDAKSTPTAEDVTA
52 GNKKIE
53 GNKKIE
54 GNKKIETH
55 GNKKIETHK
56 GNKKIETHKL
57 GNKKIETHKLT
58 GNKKIETHKL
59 GNKKIETHKL
60 GNKKIETHKL
61 GNKKIETHKL
62 GNKKIETHKL
63           TFRENAKAKTD
64           TFRENAKAKTDHGA

 1 PLVDEGAPGKQAAAQPHTEIP
```

**Tau CN fibrils 2 min**

```
  1 MAEPRQEFEVMEDHAGTYGLGDRKDQGGYTMHQDQEGDTDAEIVYKSPVVSGDTSPRHLS
  2  AEPRQEFEV
  3                                            VYKSPVVS
  4                                            VYKSPVVSGDT
  5                                            VYKSPVVSGDTSPR
  6                                            VYKSPVVSGDTSPRH
  7                                            VYKSPVVSGDTSPRHL
  8                                            VYKSPVVSGDTSPRHLS
  9                                                  VSGDTSPRHL
 10                                                  VSGDTSPRHLS
 11                                                  VSGDTSPRHLS
 12                                                   SGDTSPRHL
 13                                                           S
 14                                                           S
 15                                                           S
 16                                                           S
 17                                                           S
 18                                                           S
 19                                                           S
 20                                                           S
 21                                                           S
 22                                                           S

  1 NVSSTGSIDMVDSPQLATLADEVSASLAKQGLEGTTAEEAGIGDTPSLEDEAAGHVTQAR
  8 NV
 10 NV
 11 NVSSTGSIDMVDSPQLATL
 13 NVSSTGSIDMV
 14 NVSSTGSIDMVDSPQL
 15 NVSSTGSIDMVDSPQLA
 16 NVSSTGSIDMVDSPQLAT
 17 NVSSTGSIDMVDSPQLATL
 18 NVSSTGSIDMVDSPQLATLA
 19 NVSSTGSIDMVDSPQLATLADEV
 20 NVSSTGSIDMVDSPQLATLADEVSAS
 21 NVSSTGSIDMVDSPQLATLADEVSASLAKQ
 22 NVSSTGSIDMVDSPQLATLADEVSASLAKQGL
 23   SSTGSIDMVDSPQLA
 24   SSTGSIDMVDSPQLATL
 25   SSTGSIDMVDSPQLATLADEV
 26   SSTGSIDMVDSPQLATLADEVSAS
 27   SSTGSIDMVDSPQLATLADEVSASLAKQGL
 28      GSIDMVDSPQLA
 29      GSIDMVDSPQLATL
 30      GSIDMVDSPQLATLADEV
 31            DSPQLATLADEVSASLAKQGL
 32                LATLADEVSASLAKQGL
 33                 ATLADEVSASLAKQGL
 34                  TLADEVSASLAKQ
 35                  TLADEVSASLAKQGL
 36                   LADEVSASLAKQGL
 37                    ADEVSASLA
 38                    ADEVSASLAKQ
 39                    ADEVSASLAKQGL
 40                     DEVSASLAKQGL
 41                      EVSASLAKQGL
 42                        SASLAKQGL
 43                        SASLAKQGLEGT
 44                        SASLAKQGLEGTTAEEA
 45                          SLAKQGL

  1 MVSKSKDGTGSDDKKAKGADGKTKIATPRGAAPPGQKGQANATRIPAKTPPAPKTPPSSG
```

```
 1 EPPKSGDRSGYSSPGSPGTPGSRSRTPSLPTPPTREPKKVAVVRTPPKSPSSAKSRLQTA
46                                         AVVRTPPKSPSSAKSRLQTA
47                                                    SAKSRLQTA
48                                                       SRLQTA
49                                                        RLQTA
50                                                         LQTA
51                                                           TA
52                                                           TA
53                                                            A

 1 PVPMPDLKNVKSKIGSTENLKHQPGGGKVQIINKKLDLSNVQSKCGSKDNIKHVPGGGSV
46 PVPMPDLKNV
47 PVPMPDLKNV
48 PVPMPDLKNV
49 PVPMPDLKNV
50 PVPMPDLKNV
51 PVPMPDLKNV
52 PVPMPDLKNVKSKIGSTENL
53 PVPMPDLKNV
54 PVPMPDLKNV
55  VPMPDLKNV
56   PMPDLKNV
57           KSKIGSTENL
58            SKIGSTENL
59                     KHQPGGGKVQI
60                               IINKKLDLSNV
61                                INKKLDLSNV
62                                   KLDLSNV
63                                                    KHVPGGGSV

 1 QIVYKPVDLSKVTSKCGSLGNIHHKPGGGQVEVKSEKLDFKDRVQSKIGSLDNITHVPGG
63 QI
64  IVYKPVDLSKV
65   VYKPVDL
66   VYKPVDLS
67   VYKPVDLSKV
68   VYKPVDLSKVTS
69                                       DFKDRVQSKIGSLDNI
70                                             QSKIGSLDNI
71                                              SKIGSLDNI
72                                                          PGG
73                                                           GG
74                                                            G

 1 GNKKIETHKLTFRENAKAKTDHGAGLKESPLQTPTEDGSEEPGSETSDAKSTPTAEDVTA
72 GNKKIETHKL
73 GNKKIETHKL
74 GNKKIETHKL

 1 PLVDEGAPGKQAAAQPHTEIP
```

**Tau CN fibrils 5 min**

```
 1 MAEPRQEFEVMEDHAGTYGLGDRKDQGGYTMHQDQEGDTDAEIVYKSPVVSGDTSPRHLS
 2                                            VYKSPVVSGDT
 3                                            VYKSPVVSGDTSPR
 4                                            VYKSPVVSGDTSPRH
 5                                            VYKSPVVSGDTSPRHL
 6                                            VYKSPVVSGDTSPRHLS
 7                                            VYKSPVVSGDTSPRHLS
 8                                            VYKSPVVSGDTSPRHLS
 9                                            VYKSPVVSGDTSPRHLS
10                                            VYKSPVVSGDTSPRHLS
11                                             YKSPVVSGDTSPRHL
12                                             YKSPVVSGDTSPRHLS
13                                                  VSGDTSPRHL
14                                                   SGDTSPRHL
15                                                            S
16                                                            S
17                                                            S
18                                                            S
19                                                            S
20                                                            S
21                                                            S

 1 NVSSTGSIDMVDSPQLATLADEVSASLAKQGLEGTTAEEAGIGDTPSLEDEAAGHVTQAR
 7 N
 8 NV
 9 NVS
10 NVSST
12 NV
15 NVSSTGSIDMV
16 NVSSTGSIDMVDSPQL
17 NVSSTGSIDMVDSPQLA
18 NVSSTGSIDMVDSPQLATL
19 NVSSTGSIDMVDSPQLATLA
20 NVSSTGSIDMVDSPQLATLADEV
21 NVSSTGSIDMVDSPQLATLADEVSASLAKQGL
22   SSTGSIDMVDSPQLA
23   SSTGSIDMVDSPQLATL
24   SSTGSIDMVDSPQLATLADEV
25   SSTGSIDMVDSPQLATLADEVSASLAKQGL
26            DSPQLATLADEVSASLAKQGL
27                LATLADEVSASLAKQGL
28                 ATLADEVSASLAKQGL
29                  TLADEVSASLAKQGL
30                   LADEVSASLAKQGL
31                    ADEVSASLA
32                    ADEVSASLAKQ
33                    ADEVSASLAKQGL
34                        SASLAKQGL
35                        SASLAKQGLEGT
36                        SASLAKQGLEGTTAEEAGIGDTPSLEDEAAGHVTQA

 1 MVSKSKDGTGSDDKKAKGADGKTKIATPRGAAPPGQKGQANATRIPAKTPPAPKTPPSSG

 1 EPPKSGDRSGYSSPGSPGTPGSRSRTPSLPTPPTREPKKVAVVRTPPKSPSSAKSRLQTA
37                                         AVVRTPPKSPSSAKS
38                                         AVVRTPPKSPSSAKSRLQTA
39                                          VVRTPPKSPSSAKSRLQTA
40                                                    SAKSRLQTA
41                                                       SRLQTA
42                                                         LQTA
43                                                         LQTA
44                                                           TA
```

```
45                                                            TA
46                                                             A

 1 PVPMPDLKNVKSKIGSTENLKHQPGGGKVQIINKKLDLSNVQSKCGSKDNIKHVPGGGSV
38 PVPMPDLKNV
39 PVPMPDLKNV
40 PVPMPDLKNV
41 PVPMPDLKNV
42 PVPMPDLKNV
43 PVPMPDLKNVKSKIG
44 PVPMPDLKNVKSKIGST
45 PVPMPDLKNVKSKIGSTENL
46 PVPMPDLKNV
47           KSKIGSTENL
48            SKIGSTENL
49                     KHQPGGGKVQI
50                               IINKKLDLSNV
51                                INKKLDLSNV
52                                 NKKLDLSNV
53                                              GSKDNIKHVPGGGSV
54                                               SKDNIKHVPGGGSV
55                                                    KHVPGGGSV

 1 QIVYKPVDLSKVTSKCGSLGNIHHKPGGGQVEVKSEKLDFKDRVQSKIGSLDNITHVPGG
53 QI
54 QI
55 QI
56  IVYKPVDLSKV
57   VYKPVDL
58   VYKPVDLSKV
59   VYKPVDLSKVT
60   VYKPVDLSKVTS
61   VYKPVDLSKVTSK
62    YKPVDLSKV
63                 GSLGNIHHKPGGGQVEV
64                 GSLGNIHHKPGGGQVEVKSEKL
65                                        DFKDRVQS
66                                        DFKDRVQSKIGSL
67                                        DFKDRVQSKIGSLDNI
68                                              QSKIGSLDNI
69                                                     DNITHVPGG
70                                                        THVPGG
71                                                        THVPGG
72                                                        THVPGG
73                                                        THVPGG
74                                                        THVPGG
75                                                        THVPGG
76                                                          VPGG
77                                                           PGG
78                                                            GG
79                                                             G

 1 GNKKIETHKLTFRENAKAKTDHGAGLKESPLQTPTEDGSEEPGSETSDAKSTPTAEDVTA
69 GNKKIETHKL
70 GNKKIE
71 GNKKIET
72 GNKKIETH
73 GNKKIETHK
74 GNKKIETHKL
75 GNKKIETHKLT
76 GNKKIETHKL
77 GNKKIETHKL
78 GNKKIETHKL
79 GNKKIETHKL
```

```
80 GNKKIETHKL
81           TFRENAKAKTD
82           TFRENAKAKTDHGA

 1 PLVDEGAPGKQAAAQPHTEIP
```

**Tau CN fibrils 10 min**

```
 1 MAEPRQEFEVMEDHAGTYGLGDRKDQGGYTMHQDQEGDTDAEIVYKSPVVSGDTSPRHLS
 2                                            VYKSPVVS
 3                                            VYKSPVVSGDTSPR
 4                                            VYKSPVVSGDTSPRH
 5                                            VYKSPVVSGDTSPRHL
 6                                            VYKSPVVSGDTSPRHLS
 7                                            VYKSPVVSGDTSPRHLS
 8                                            VYKSPVVSGDTSPRHLS
 9                                            VYKSPVVSGDTSPRHLS
10                                            VYKSPVVSGDTSPRHLS
11                                            VYKSPVVSGDTSPRHLS
12                                            VYKSPVVSGDTSPRHLS
13                                             YKSPVVSGDTSPRHL
14                                             YKSPVVSGDTSPRHLS
15                                               SPVVSGDTSPRHL
16                                                  VSGDTSPRHL
17                                                  VSGDTSPRHLS
18                                                  VSGDTSPRHLS
19                                                   SGDTSPRHL
20                                                   SGDTSPRHLS
21                                                            S
22                                                            S
23                                                            S
24                                                            S
25                                                            S
26                                                            S
27                                                            S
28                                                            S
29                                                            S

 1 NVSSTGSIDMVDSPQLATLADEVSASLAKQGLEGTTAEEAGIGDTPSLEDEAAGHVTQAR
 7 N
 8 NV
 9 NVS
10 NVSST
11 NVSSTG
12 NVSSTGSIDMVDSPQLA
14 NV
17 NV
18 NVSSTGSIDMVDSPQLATL
20 NV
21 NVSSTGSIDMV
22 NVSSTGSIDMVDSPQL
23 NVSSTGSIDMVDSPQLA
24 NVSSTGSIDMVDSPQLAT
25 NVSSTGSIDMVDSPQLATL
26 NVSSTGSIDMVDSPQLATLA
27 NVSSTGSIDMVDSPQLATLADEV
28 NVSSTGSIDMVDSPQLATLADEVSAS
29 NVSSTGSIDMVDSPQLATLADEVSASLAKQGL
30   SSTGSIDMVDSPQLA
31   SSTGSIDMVDSPQLATL
32   SSTGSIDMVDSPQLATLADEV
33   SSTGSIDMVDSPQLATLADEVSAS
34   SSTGSIDMVDSPQLATLADEVSASLAKQGL
35      GSIDMVDSPQLATL
36            DSPQLATLADEVSASLAKQGL
37                LATLADEVSASLAKQGL
38                 ATLADEVSASLAKQGL
39                  TLADEVSASLAKQ
40                  TLADEVSASLAKQGL
41                   LADEVSASLAKQGL
```

```
42                    ADEVSASLA
43                    ADEVSASLAKQ
44                    ADEVSASLAKQGL
45                     DEVSASLAKQGL
46                      EVSASLAKQGL
47                        SASLAKQGL
48                        SASLAKQGLEGT
49                        SASLAKQGLEGTTAEEA
50                          SLAKQGL

 1 MVSKSKDGTGSDDKKAKGADGKTKIATPRGAAPPGQKGQANATRIPAKTPPAPKTPPSSG

 1 EPPKSGDRSGYSSPGSPGTPGSRSRTPSLPTPPTREPKKVAVVRTPPKSPSSAKSRLQTA
51                      SRSRTPSLPTPPTREPKKV
52                                         AVVRTPPKSPSSAKS
53                                         AVVRTPPKSPSSAKSRLQ
54                                         AVVRTPPKSPSSAKSRLQTA
55                                          VVRTPPKSPSSAKSRLQTA
56                                           VRTPPKSPSSAKSRLQTA
57                                            RTPPKSPSSAKSRLQTA
58                                                    SAKSRLQTA
59                                                    SAKSRLQTA
60                                                       SRLQTA
61                                                        RLQTA
62                                                         LQTA
63                                                         LQTA
64                                                           TA
65                                                           TA
66                                                            A

 1 PVPMPDLKNVKSKIGSTENLKHQPGGGKVQIINKKLDLSNVQSKCGSKDNIKHVPGGGSV
54 PVPMPDLKNV
55 PVPMPDLKNV
56 PVPMPDLKNV
57 PVPMPDLKNV
58 PVPMPDLKNV
59 PVPMPDLKNVKSKIGSTENL
60 PVPMPDLKNV
61 PVPMPDLKNV
62 PVPMPDLKNV
63 PVPMPDLKNVKSKIG
64 PVPMPDLKNV
65 PVPMPDLKNVKSKIGSTENL
66 PVPMPDLKNV
67 PVPMPDLKNV
68           KSKIGSTENL
69           KSKIGSTENLKHQPGGGKVQ
70            SKIGSTENL
71                     KHQPGGGKVQI
72                     KHQPGGGKVQIINKKLDLSNV
73                               IINKKLDLSNV
74                               IINKKLDLSNVQS
75                                INKKLDLSNV
76                                 NKKLDLSNV
77                                  KKLDLSNV
78                                   KLDLSNV
79                                              GSKDNIKHVPGGGSV
80                                               SKDNIKHVPGGGSV
81                                                    KHVPGGGSV

 1 QIVYKPVDLSKVTSKCGSLGNIHHKPGGGQVEVKSEKLDFKDRVQSKIGSLDNITHVPGG
79 QI
80 QI
81 QI
```

```
 82  IVYKPVDLSKV
 83  IVYKPVDLSKVTS
 84   VYKPVDL
 85   VYKPVDLS
 86   VYKPVDLSKV
 87   VYKPVDLSKVT
 88   VYKPVDLSKVTS
 89   VYKPVDLSKVTSK
 90   VYKPVDLSKVTSKC
 91    YKPVDLSKV
 92                 GSLGNIHHKPGGGQ
 93                 GSLGNIHHKPGGGQVEV
 94                 GSLGNIHHKPGGGQVEVKSEKL
 95                  SLGNIHHKPGGGQVE
 96                                       DFKDRVQ
 97                                       DFKDRVQS
 98                                       DFKDRVQSK
 99                                       DFKDRVQSKIG
100                                       DFKDRVQSKIGSL
101                                       DFKDRVQSKIGSLDNI
102                                       DFKDRVQSKIGSLDNIT
103                                             QSKIGSLDNI
104                                             QSKIGSLDNITHVPGG
105                                                    DNITHVPGG
106                                                    DNITHVPGG
107                                                       THVPGG
108                                                       THVPGG
109                                                       THVPGG
110                                                       THVPGG
111                                                       THVPGG
112                                                       THVPGG
113                                                         VPGG
114                                                          PGG
115                                                          PGG
116                                                           GG
117                                                           GG
118                                                            G

  1 GNKKIETHKLTFRENAKAKTDHGAGLKESPLQTPTEDGSEEPGSETSDAKSTPTAEDVTA
104 GNKKIETHKL
105 GNKKIE
106 GNKKIETHKL
107 GNKKIE
108 GNKKIET
109 GNKKIETH
110 GNKKIETHK
111 GNKKIETHKL
112 GNKKIETHKLT
113 GNKKIETHKL
114 GNKKIE
115 GNKKIETHKL
116 GNKKIE
117 GNKKIETHKL
118 GNKKIETHKL
119 GNKKIETHKL
120           TFRENAKAKTD
121           TFRENAKAKTDHGA
122           TFRENAKAKTDHGAGLK
123           TFRENAKAKTDHGAGLKESPLQTPTEDGSEEPGSE

  1 PLVDEGAPGKQAAAQPHTEIP
```

**Tau CN fibrils 120 min**

```
 1 MAEPRQEFEVMEDHAGTYGLGDRKDQGGYTMHQDQEGDTDAEIVYKSPVVSGDTSPRHLS
 2  AEPRQEFEVM
 3                                            VYKSPVVS
 4                                            VYKSPVVSGDT
 5                                            VYKSPVVSGDTSPR
 6                                            VYKSPVVSGDTSPRH
 7                                            VYKSPVVSGDTSPRHL
 8                                            VYKSPVVSGDTSPRHLS
 9                                            VYKSPVVSGDTSPRHLS
10                                            VYKSPVVSGDTSPRHLS
11                                            VYKSPVVSGDTSPRHLS
12                                            VYKSPVVSGDTSPRHLS
13                                            VYKSPVVSGDTSPRHLS
14                                            VYKSPVVSGDTSPRHLS
15                                             YKSPVVSGDTSPRHL
16                                             YKSPVVSGDTSPRHLS
17                                               SPVVSGDTSPRHL
18                                                  VSGDTSPRHL
19                                                  VSGDTSPRHLS
20                                                  VSGDTSPRHLS
21                                                   SGDTSPRHL
22                                                   SGDTSPRHLS
23                                                          HLS
24                                                          HLS
25                                                            S
26                                                            S
27                                                            S
28                                                            S
29                                                            S
30                                                            S
31                                                            S
32                                                            S
33                                                            S

 1 NVSSTGSIDMVDSPQLATLADEVSASLAKQGLEGTTAEEAGIGDTPSLEDEAAGHVTQAR
 9 N
10 NV
11 NVS
12 NVSST
13 NVSSTG
14 NVSSTGSIDMVDSPQLA
16 NV
19 NV
20 NVSSTGSIDMVDSPQLATL
22 NV
23 NVSSTGSIDMVDSPQLATL
24 NVSSTGSIDMVDSPQLATLADEV
25 NVSSTGSIDMV
26 NVSSTGSIDMVDSPQL
27 NVSSTGSIDMVDSPQLA
28 NVSSTGSIDMVDSPQLAT
29 NVSSTGSIDMVDSPQLATL
30 NVSSTGSIDMVDSPQLATLA
31 NVSSTGSIDMVDSPQLATLADEV
32 NVSSTGSIDMVDSPQLATLADEVSAS
33 NVSSTGSIDMVDSPQLATLADEVSASLAKQGL
34   SSTGSIDMVDSPQLA
35   SSTGSIDMVDSPQLATL
36   SSTGSIDMVDSPQLATLADEV
37   SSTGSIDMVDSPQLATLADEVSAS
38   SSTGSIDMVDSPQLATLADEVSASLAKQGL
39      GSIDMVDSPQLA
```

```
40      GSIDMVDSPQLATL
41       SIDMVDSPQLATL
42            DSPQLATLADEV
43                  TLADEVSASLAKQGL
44                   LADEVSASLAKQGL
45                    ADEVSASLA
46                    ADEVSASLAKQ
47                    ADEVSASLAKQGL
48                        SASLAKQGL
49                        SASLAKQGLEGT
50                        SASLAKQGLEGTTAEEA
51                                         GIGDTPSLEDEAAGHVTQA
52                                         GIGDTPSLEDEAAGHVTQAR
53                                             TPSLEDEAAGHVTQA

 1 MVSKSKDGTGSDDKKAKGADGKTKIATPRGAAPPGQKGQANATRIPAKTPPAPKTPPSSG
52 MV

 1 EPPKSGDRSGYSSPGSPGTPGSRSRTPSLPTPPTREPKKVAVVRTPPKSPSSAKSRLQTA
54                      SRSRTPSLPTPPTREPKKV
55                                         AVVRTPPKSPS
56                                         AVVRTPPKSPSSA
57                                         AVVRTPPKSPSSAKS
58                                         AVVRTPPKSPSSAKSRL
59                                         AVVRTPPKSPSSAKSRLQ
60                                         AVVRTPPKSPSSAKSRLQT
61                                         AVVRTPPKSPSSAKSRLQTA
62                                         AVVRTPPKSPSSAKSRLQTA
63                                          VVRTPPKSPS
64                                          VVRTPPKSPSSAKSRLQTA
65                                           VRTPPKSPS
66                                           VRTPPKSPSSAKSRLQTA
67                                            RTPPKSPSSAKS
68                                            RTPPKSPSSAKSRLQ
69                                            RTPPKSPSSAKSRLQT
70                                            RTPPKSPSSAKSRLQTA
71                                             TPPKSPSSAKSRLQTA
72                                               PKSPSSAKSRLQ
73                                                 SPSSAKSRLQTA
74                                                   SSAKSRLQTA
75                                                    SAKSRLQTA
76                                                    SAKSRLQTA
77                                                      KSRLQTA
78                                                       SRLQTA
79                                                        RLQTA
80                                                         LQTA
81                                                         LQTA
82                                                          QTA
83                                                           TA
84                                                           TA
85                                                           TA
86                                                            A

 1 PVPMPDLKNVKSKIGSTENLKHQPGGGKVQIINKKLDLSNVQSKCGSKDNIKHVPGGGSV
61 PVPMPDL
62 PVPMPDLKNV
64 PVPMPDLKNV
66 PVPMPDLKNV
70 PVPMPDLKNV
71 PVPMPDLKNV
73 PVPMPDLKNV
74 PVPMPDLKNV
75 PVPMPDLKNV
76 PVPMPDLKNVKSKIGSTENL
```

```
 77 PVPMPDLKNV
 78 PVPMPDLKNV
 79 PVPMPDLKNV
 80 PVPMPDLKNV
 81 PVPMPDLKNVKSKIGSTENL
 82 PVPMPDLKNV
 83 PVPMPDLKNV
 84 PVPMPDLKNVKSKIGST
 85 PVPMPDLKNVKSKIGSTENL
 86 PVPMPDLKNV
 87 PVPMPDLKNV
 88  VPMPDLKNV
 89   PMPDLKNV
 90           KSKIGSTENL
 91           KSKIGSTENLKHQPGGGKVQ
 92           KSKIGSTENLKHQPGGGKVQI
 93            SKIGSTENL
 94            SKIGSTENLK
 95                     KHQPGGGKVQI
 96                     KHQPGGGKVQII
 97                     KHQPGGGKVQIINK
 98                     KHQPGGGKVQIINKKLDL
 99                     KHQPGGGKVQIINKKLDLSNV
100                              QIINKKLDLSNV
101                               IINKKLDLSNV
102                               IINKKLDLSNVQS
103                                INKKLDLSNV
104                                INKKLDLSNVQS
105                                 NKKLDLSNV
106                                  KKLDLSNV
107                                   KLDLSNV
108                                              GSKDNIKHVPGGGSV
109                                               SKDNIKHVPGGGSV
110                                                    KHVPGGGSV

  1 QIVYKPVDLSKVTSKCGSLGNIHHKPGGGQVEVKSEKLDFKDRVQSKIGSLDNITHVPGG
108 QI
109 QI
110 QI
111  IVYKPVDLSKV
112   VYKPVDL
113   VYKPVDLSKV
114   VYKPVDLSKVT
115   VYKPVDLSKVTS
116   VYKPVDLSKVTSK
117   VYKPVDLSKVTSKC
118    YKPVDLSKV
119     KPVDLSKV
120                 GSLGNIHHKPGGGQ
121                 GSLGNIHHKPGGGQVEV
122                 GSLGNIHHKPGGGQVEVKSEKL
123                   LGNIHHKPGGGQVEVKSEKL
124                    GNIHHKPGGGQVEVKSEKL
125                                      LDFKDRVQSK
126                                       DFKDRVQ
127                                       DFKDRVQS
128                                       DFKDRVQSK
129                                       DFKDRVQSKIGSL
130                                       DFKDRVQSKIGSLD
131                                       DFKDRVQSKIGSLDNI
132                                       DFKDRVQSKIGSLDNIT
133                                       DFKDRVQSKIGSLDNITHV
134                                            VQSKIGSLDNI
135                                             QSKIGSLDNI
```

```
136                                             QSKIGSLDNIT
137                                             QSKIGSLDNITHVPGG
138                                              SKIGSLDNI
139                                                    DNITHVPGG
140                                                    DNITHVPGG
141                                                       THVPGG
142                                                       THVPGG
143                                                       THVPGG
144                                                       THVPGG
145                                                       THVPGG
146                                                       THVPGG
147                                                         VPGG
148                                                          PGG
149                                                          PGG
150                                                           GG
151                                                           GG
152                                                            G

  1 GNKKIETHKLTFRENAKAKTDHGAGLKESPLQTPTEDGSEEPGSETSDAKSTPTAEDVTA
137 GNKKIETHKL
139 GNKKIE
140 GNKKIETHKL
141 GNKKIE
142 GNKKIET
143 GNKKIETH
144 GNKKIETHK
145 GNKKIETHKL
146 GNKKIETHKLT
147 GNKKIETHKL
148 GNKKIE
149 GNKKIETHKL
150 GNKKIE
151 GNKKIETHKL
152 GNKKIETHKL
153 GNKKIETHKL
154           TFRENAKAKTD
155           TFRENAKAKTDHGA
156           TFRENAKAKTDHGAGLK

  1 PLVDEGAPGKQAAAQPHTEIP
157            AAAQPHTEIP
```

#### Supplemental data 5

### Overview

| ACE project | Title |
|---|---|
| ACE_0711-02 | Time-resolved analysis of proteolytic cleavage sites in Tau (10µM) by HTRA1 (2 µM) (0s, 15s, 1m, 2m, 5m, 10m, 120m) in NaPi |
| ACE_0711-04 | Time-resolved analysis of proteolytic cleavage sites in fibrillar Tau (10µM) by HTRA1 (2 µM) (0s, 15s, 1m, 2m, 5m, 10m, 120m) in HEPES |
| ACE_0809-03 | Time-resolved analysis of proteolytic cleavage sites in fibrillar Tau_CN by HTRA1 (t= 0; 15 sec; 1 min; 2 min; 5 min; 10 min; 120 min) |

#### ACE_0711-02

#### File legend

| ACE ID | Alternate ID | Organ/ cell line | Treatment/ experimental setup |
|---|---|---|---|
| ACE_0711-02_BH01 | MS8_H_0 | Human (overexpressed in *E. coli*) | 2 µM HTRA1 Control, 0 sec, supernatant after acetone precipitation |
| ACE_0711-02_BH02 | MS8_H_0 | dito | 2 µM HTRA1 Control, 0 sec, supernatant after acetone precipitation |
| ACE_0711-02_BH03 | MS8_H_0 | dito | 2 µM HTRA1 Control, 0 sec, supernatant after acetone precipitation |
| ACE_0711-02_BH04 | MS8_H_0 | dito | 2 µM HTRA1 Control, 0 sec, supernatant after acetone precipitation |
| ACE_0711-02_BH05 | MS8_H_120 | dito | 2 µM HTRA1 Control, 120 min, supernatant after acetone precipitation |
| ACE_0711-02_BH06 | MS8_H_120 | dito | 2 µM HTRA1 Control, 120 min, supernatant after acetone precipitation |
| ACE_0711-02_BH07 | MS8_H_120 | dito | 2 µM HTRA1 Control, 120 min, supernatant after acetone precipitation |
| ACE_0711-02_BH08 | MS8_H_120 | dito | 2 µM HTRA1 Control, 120 min, supernatant after acetone precipitation |
| ACE_0711-02_BH09 | MS8_T_0 | dito | 10 µM Tau Control, 0 sec, supernatant after acetone precipitation |
| ACE_0711-02_BH10 | MS8_T_0 | dito | 10 µM Tau Control, 0 sec, supernatant after acetone precipitation |
| ACE_0711-02_BH11 | MS8_T_0 | dito | 10 µM Tau Control, 0 sec, supernatant after acetone precipitation |
| ACE_0711-02_BH12 | MS8_T_0 | dito | 10 µM Tau Control, 0 sec, supernatant after acetone precipitation |
| ACE_0711-02_BH13 | MS8_T_120 | dito | 10 µM Tau Control, 120 min, supernatant after acetone precipitation |
| ACE_0711-02_BH14 | MS8_T_120 | dito | 10 µM Tau Control, 120 min, supernatant after acetone precipitation |
| ACE_0711-02_BH15 | MS8_T_120 | dito | 10 µM Tau Control, 120 min, supernatant after acetone precipitation |
| ACE_0711-02_BH16 | MS8_T_120 | dito | 10 µM Tau Control, 120 min, supernatant after acetone precipitation |
| ACE_0711-02_BH17 | MS8_TH_0 | dito | 10 µM Tau + 2 µM HTRA1, 0 sec, supernatant after acetone precipitation |
| ACE_0711-02_BH18 | MS8_TH_0 | dito | 10 µM Tau + 2 µM HTRA1, 0 sec, supernatant after acetone precipitation |
| ACE_0711-02_BH19 | MS8_TH_0 | dito | 10 µM Tau + 2 µM HTRA1, 0 sec, supernatant after acetone precipitation |
| ACE_0711-02_BH20 | MS8_TH_0 | dito | 10 µM Tau + 2 µM HTRA1, 0 sec, supernatant after acetone precipitation |
| ACE_0711-02_BH21 | MS8_TH_15 | dito | 10 µM Tau + 2 µM HTRA1, 15 sec, supernatant after acetone precipitation |
| ACE_0711-02_BH22 | MS8_TH_15 | dito | 10 µM Tau + 2 µM HTRA1, 15 sec, supernatant after acetone precipitation |
| ACE_0711-02_BH23 | MS8_TH_15 | dito | 10 µM Tau + 2 µM HTRA1, 15 sec, supernatant after acetone precipitation |
| ACE_0711-02_BH24 | MS8_TH_15 | dito | 10 µM Tau + 2 µM HTRA1, 15 sec, supernatant after acetone precipitation |
| ACE_0711-02_BH25 | MS8_TH_1 | dito | 10 µM Tau + 2 µM HTRA1, 1 min, supernatant after acetone precipitation |
| ACE_0711-02_BH26 | MS8_TH_1 | dito | 10 µM Tau + 2 µM HTRA1, 1 min, supernatant after acetone precipitation |
| ACE_0711-02_BH27 | MS8_TH_1 | dito | 10 µM Tau + 2 µM HTRA1, 1 min, supernatant after acetone precipitation |
| ACE_0711-02_BH28 | MS8_TH_1 | dito | 10 µM Tau + 2 µM HTRA1, 1 min, supernatant after acetone precipitation |
| ACE_0711-02_BH29 | MS8_TH_5 | dito | 10 µM Tau + 2 µM HTRA1, 5 min, supernatant after acetone precipitation |
| ACE_0711-02_BH30 | MS8_TH_5 | dito | 10 µM Tau + 2 µM HTRA1, 5 min, supernatant after acetone precipitation |
| ACE_0711-02_BH31 | MS8_TH_5 | dito | 10 µM Tau + 2 µM HTRA1, 5 min, supernatant after acetone precipitation |
| ACE_0711-02_BH32 | MS8_TH_5 | dito | 10 µM Tau + 2 µM HTRA1, 5 min, supernatant after acetone precipitation |
| ACE_0711-02_BH33 | MS8_TH_10 | dito | 10 µM Tau + 2 µM HTRA1, 10 min, supernatant after acetone precipitation |
| ACE_0711-02_BH34 | MS8_TH_10 | dito | 10 µM Tau + 2 µM HTRA1, 10 min, supernatant after acetone precipitation |
| ACE_0711-02_BH35 | MS8_TH_10 | dito | 10 µM Tau + 2 µM HTRA1, 10 min, supernatant after acetone precipitation |
| ACE_0711-02_BH36 | MS8_TH_10 | dito | 10 µM Tau + 2 µM HTRA1, 10 min, supernatant after acetone precipitation |
| ACE_0711-02_BH37 | MS8_TH_2 | dito | 10 µM Tau + 2 µM HTRA1, 2 min, supernatant after acetone precipitation |
| ACE_0711-02_BH38 | MS8_TH_2 | dito | 10 µM Tau + 2 µM HTRA1, 2 min, supernatant after acetone precipitation |

| | | | |
|---|---|---|---|
| ACE_0711-02_BH39 | MS8_TH_2 | dito | 10 µM Tau + 2 µM HTRA1, 2 min, supernatant after acetone precipitation |
| ACE_0711-02_BH40 | MS8_TH_2 | dito | 10 µM Tau + 2 µM HTRA1, 2 min, supernatant after acetone precipitation |
| ACE_0711-02_BH41 | MS8_TH_120 | dito | 10 µM Tau + 2 µM HTRA1, 120 min, supernatant after acetone precipitation |
| ACE_0711-02_BH42 | MS8_TH_120 | dito | 10 µM Tau + 2 µM HTRA1, 120 min, supernatant after acetone precipitation |
| ACE_0711-02_BH43 | MS8_TH_120 | dito | 10 µM Tau + 2 µM HTRA1, 120 min, supernatant after acetone precipitation |
| ACE_0711-02_BH44 | MS8_TH_120 | dito | 10 µM Tau + 2 µM HTRA1, 120 min, supernatant after acetone precipitation |

#### LC_Settings

| | |
|---|---|
| MS device | Orbitrap Elite |
| LC device | Evosep One |
| ion source | Thermo Nanospray Flex |
| **Analytical column** | EV1109 Analytical Column – 60 & 100 samples/day |
| column diameter | Length ($L_C$) = 8 cm; ID = 150; OD = PEEK; emitter EV-1086 Stainless steel emitter |
| stationary phase | Dr Maisch C18 AQ, 1.5 µm beads |
| particle diameter ($d_P$) | 1.5 µm |
| Pore size | 120 Å |
| Column ID | AC_EVO-006 |
| Column oven | na |
| Column oven temp. | na |
| **solvents** | A: 0.1% FA in UPLC water<br>B: 0.1% FA in UPLC ACN |
| gradient | 21 min gradient (60 SPD) |

#### MS_Settings

| Project | MS | general | MS1 | MS2 | MS2 | MS3 | Comments; special settings |
|---|---|---|---|---|---|---|---|
| ACE_0644 | Elite | Tune v2.7.0.1112 SP2<br>Gradient: 21 min | Analyzer: FT<br>Res.: 60000<br>SR: 300 - 1500<br>AGC: $3\times 10^6$<br>AcT: 50<br>RF: 30<br>SF: --<br>DDM: NS/15 | Analyzer: IT<br>Res./ScR: -/rapid<br>SR: Auto<br>AGC: $1\times 10^4$<br>AcT: 60<br>CS: +2 and higher<br>IsM: IT<br>IsW: 2.0<br>Frag.: CID<br>NCE: 35 | | | classic orbitrap experiment: MS1 in Orbitrap at high resolution and data dependent MS2 in Iontrap at rapid scan rate. Dynamic exclusion enabled (exclude after n times=1; Exclusion duration (s)= 30; mass tolerance= ± 100 ppm), exclusion list size: 500 |

Note: **FT**= Fourier Transform (Orbitrap); **IT**= Iontrap; **Q**= Quadrupol; **Res.**= max. Resolution at 200 m/z (Lumos) or 400 m/z (Elite) [FWHM (full width at half maximum)]; **ScR**= scan rate for measurements in the IT; **SR**= scan range [m/z]; **AGC**= automatic gain control, max number of acquired ions per measurement; **AcT**= max. Ion acquisition time [ms]; **CS**= charge states used for fragmentation; **IsM**= Isolation mode (Q or IT), MS2 isolation and further is only done in IT; **IsW**= Isolation window [m/z], value followed by scan mode the isolation is based on (MS1, MS2 …) **Frag.**= Fragmentation method; **HCD**= Higher-energy collisional dissociation; **CID**= Collision-induced dissociation; **ETD**= Electron-transfer dissociation**; EThcD=** Electron-Transfer/Higher-Energy Collision Dissociation; **sHCD**= stepped HCD**; NCE**= normalized collision energy; **cycles**: number of MSn recorded or max cycle time; RF= RF Lens [%]; **SF**= Source Fragmentation [V]; **DDM**: Data dependent Mode (cycle time in seconds, CT/[s] or number of scans, NS); **NS**= Number of data dependent scans

#### Search Settings

| | |
|---|---|
| Program & version | MaxQuant v2.0.2.0 |
| Search engine | Andromeda |
| settings | Basically default; LFQ and MBR were turned on |
| Static modification | none |
| Digestion mode | unspecific |
| | Min. peptide length for unspecific search 6<br>Max. peptide length for unspecific search 36 |
| Dynamic modification | Acetyl (N-term); Oxidation (M) |
| Modification included in quantification | Oxidation (M) |
| Databases | 1. Contaminants<br>2. ACE_0699_UP000000625_83333.fasta |
| Annotation | |

### ACE_0711-04

#### File legend

| ACE ID | Alternate ID | Organ/ cell line | Treatment/ experimental setup |
|---|---|---|---|
| ACE_0711-02_BH01 | MS5_H_0 | Human (overexpressed in *E. coli*) | 2 µM HTRA1 Control, 0 sec, supernatant after acetone precipitation |
| ACE_0711-02_BH02 | MS5_H_0 | dito | 2 µM HTRA1 Control, 0 sec, supernatant after acetone precipitation |
| ACE_0711-02_BH03 | MS5_H_0 | dito | 2 µM HTRA1 Control, 0 sec, supernatant after acetone precipitation |
| ACE_0711-02_BH04 | MS5_H_0 | dito | 2 µM HTRA1 Control, 0 sec, supernatant after acetone precipitation |
| ACE_0711-02_BH05 | MS5_H_120 | dito | 2 µM HTRA1 Control, 120 min, supernatant after acetone precipitation |
| ACE_0711-02_BH06 | MS5_H_120 | dito | 2 µM HTRA1 Control, 120 min, supernatant after acetone precipitation |
| ACE_0711-02_BH07 | MS5_H_120 | dito | 2 µM HTRA1 Control, 120 min, supernatant after acetone precipitation |
| ACE_0711-02_BH08 | MS5_H_120 | dito | 2 µM HTRA1 Control, 120 min, supernatant after acetone precipitation |
| ACE_0711-02_BH09 | MS5_T_0 | dito | 10 µM Tau Control, 0 sec, supernatant after acetone precipitation |
| ACE_0711-02_BH10 | MS5_T_0 | dito | 10 µM Tau Control, 0 sec, supernatant after acetone precipitation |
| ACE_0711-02_BH11 | MS5_T_0 | dito | 10 µM Tau Control, 0 sec, supernatant after acetone precipitation |
| ACE_0711-02_BH12 | MS5_T_0 | dito | 10 µM Tau Control, 0 sec, supernatant after acetone precipitation |
| ACE_0711-02_BH13 | MS5_T_120 | dito | 10 µM Tau Control, 120 min, supernatant after acetone precipitation |
| ACE_0711-02_BH14 | MS5_T_120 | dito | 10 µM Tau Control, 120 min, supernatant after acetone precipitation |
| ACE_0711-02_BH15 | MS5_T_120 | dito | 10 µM Tau Control, 120 min, supernatant after acetone precipitation |
| ACE_0711-02_BH16 | MS5_T_120 | dito | 10 µM Tau Control, 120 min, supernatant after acetone precipitation |
| ACE_0711-02_BH17 | MS5_TH_0 | dito | 10 µM Tau + 2 µM HTRA1, 0 sec, supernatant after acetone precipitation |
| ACE_0711-02_BH18 | MS5_TH_0 | dito | 10 µM Tau + 2 µM HTRA1, 0 sec, supernatant after acetone precipitation |
| ACE_0711-02_BH19 | MS5_TH_0 | dito | 10 µM Tau + 2 µM HTRA1, 0 sec, supernatant after acetone precipitation |
| ACE_0711-02_BH20 | MS5_TH_0 | dito | 10 µM Tau + 2 µM HTRA1, 0 sec, supernatant after acetone precipitation |
| ACE_0711-02_BH21 | MS5_TH_15 | dito | 10 µM Tau + 2 µM HTRA1, 15 sec, supernatant after acetone precipitation |
| ACE_0711-02_BH22 | MS5_TH_15 | dito | 10 µM Tau + 2 µM HTRA1, 15 sec, supernatant after acetone precipitation |
| ACE_0711-02_BH23 | MS5_TH_15 | dito | 10 µM Tau + 2 µM HTRA1, 15 sec, supernatant after acetone precipitation |
| ACE_0711-02_BH24 | MS5_TH_15 | dito | 10 µM Tau + 2 µM HTRA1, 15 sec, supernatant after acetone precipitation |
| ACE_0711-02_BH25 | MS5_TH_1 | dito | 10 µM Tau + 2 µM HTRA1, 1 min, supernatant after acetone precipitation |
| ACE_0711-02_BH26 | MS5_TH_1 | dito | 10 µM Tau + 2 µM HTRA1, 1 min, supernatant after acetone precipitation |
| ACE_0711-02_BH27 | MS5_TH_1 | dito | 10 µM Tau + 2 µM HTRA1, 1 min, supernatant after acetone precipitation |
| ACE_0711-02_BH28 | MS5_TH_1 | dito | 10 µM Tau + 2 µM HTRA1, 1 min, supernatant after acetone precipitation |
| ACE_0711-02_BH29 | MS5_TH_2 | dito | 10 µM Tau + 2 µM HTRA1, 2 min, supernatant after acetone precipitation |
| ACE_0711-02_BH30 | MS5_TH_2 | dito | 10 µM Tau + 2 µM HTRA1, 2 min, supernatant after acetone precipitation |
| ACE_0711-02_BH31 | MS5_TH_2 | dito | 10 µM Tau + 2 µM HTRA1, 2 min, supernatant after acetone precipitation |
| ACE_0711-02_BH32 | MS5_TH_2 | dito | 10 µM Tau + 2 µM HTRA1, 2 min, supernatant after acetone precipitation |
| ACE_0711-02_BH33 | MS5_TH_5 | dito | 10 µM Tau + 2 µM HTRA1, 5 min, supernatant after acetone precipitation |
| ACE_0711-02_BH34 | MS5_TH_5 | dito | 10 µM Tau + 2 µM HTRA1, 5 min, supernatant after acetone precipitation |
| ACE_0711-02_BH35 | MS5_TH_5 | dito | 10 µM Tau + 2 µM HTRA1, 5 min, supernatant after acetone precipitation |
| ACE_0711-02_BH36 | MS5_TH_5 | dito | 10 µM Tau + 2 µM HTRA1, 5 min, supernatant after acetone precipitation |
| ACE_0711-02_BH37 | MS5_TH_10 | dito | 10 µM Tau + 2 µM HTRA1, 10 min, supernatant after acetone precipitation |
| ACE_0711-02_BH38 | MS5_TH_10 | dito | 10 µM Tau + 2 µM HTRA1, 10 min, supernatant after acetone precipitation |

| | | | |
|---|---|---|---|
| ACE_0711-02_BH39 | MS5_TH_10 | dito | 10 µM Tau + 2 µM HTRA1, 10 min, supernatant after acetone precipitation |
| ACE_0711-02_BH40 | MS5_TH_10 | dito | 10 µM Tau + 2 µM HTRA1, 10 min, supernatant after acetone precipitation |
| ACE_0711-02_BH41 | MS5_TH_120 | dito | 10 µM Tau + 2 µM HTRA1, 120 min, supernatant after acetone precipitation |
| ACE_0711-02_BH42 | MS5_TH_120 | dito | 10 µM Tau + 2 µM HTRA1, 120 min, supernatant after acetone precipitation |
| ACE_0711-02_BH43 | MS5_TH_120 | dito | 10 µM Tau + 2 µM HTRA1, 120 min, supernatant after acetone precipitation |
| ACE_0711-02_BH44 | MS5_TH_120 | dito | 10 µM Tau + 2 µM HTRA1, 120 min, supernatant after acetone precipitation |

### LC_Settings

| | |
|---|---|
| MS device | Orbitrap Elite |
| LC device | Evosep One |
| ion source | Thermo Nanospray Flex |
| **Analytical column** | EV1109 Analytical Column – 60 & 100 samples/day |
| column diameter | Length ($L_C$) = 8 cm; ID = 150; OD = PEEK; emitter EV-1086 Stainless steel emitter |
| stationary phase | Dr Maisch C18 AQ, 1.5 µm beads |
| particle diameter ($d_p$) | 1.5 µm |
| Pore size | 120 Å |
| Column ID | AC_EVO-006 |
| Column oven | na |
| Column oven temp. | na |
| **solvents** | A: 0.1% FA in UPLC water<br>B: 0.1% FA in UPLC ACN |
| gradient | 21 min gradient (60 SPD) |

### MS_Settings

| Project | MS | general | MS1 | MS2 | MS2 | MS3 | Comments; special settings |
|---|---|---|---|---|---|---|---|
| ACE_0644 | Elite | Tune v2.7.0.1112 SP2<br>Gradient: 21 min | Analyzer: FT<br>Res.: 60000<br>SR: 300 - 1500<br>AGC: $3\times 10^6$<br>AcT: 50<br>RF: 30<br>SF: --<br>DDM: NS/15 | Analyzer: IT<br>Res./ScR: -/rapid<br>SR: Auto<br>AGC: $1\times 10^4$<br>AcT: 60<br>CS: +2 and higher<br>IsM: IT<br>IsW: 2.0<br>Frag.: CID<br>NCE: 35 | | | classic orbitrap experiment: MS1 in Orbitrap at high resolution and data dependent MS2 in Iontrap at rapid scan rate. Dynamic exclusion enabled (exclude after n times=1; Exclusion duration (s)= 30; mass tolerance= ± 100 ppm), exclusion list size: 500 |

Note: **FT**= Fourier Transform (Orbitrap); **IT**= Iontrap; **Q**= Quadrupol; **Res.**= max. Resolution at 200 m/z (Lumos) or 400 m/z (Elite) [FWHM (full width at half maximum)]; **ScR**= scan rate for measurements in the IT; **SR**= scan range [m/z]; **AGC**= automatic gain control, max number of acquired ions per measurement; **AcT**= max. Ion acquisition time [ms]; **CS**= charge states used for fragmentation; **IsM**= Isolation mode (Q or IT), MS2 isolation and further is only done in IT; **IsW**= Isolation window [m/z], value followed by scan mode the isolation is based on (MS1, MS2 …) **Frag.**= Fragmentation method; **HCD**= Higher-energy collisional dissociation; **CID**= Collision-induced dissociation; **ETD**= Electron-transfer dissociation**; EThcD=** Electron-Transfer/Higher-Energy Collision Dissociation; **sHCD**= stepped HCD**; NCE**= normalized collision energy; **cycles**: number of MSn recorded or max cycle time; RF= RF Lens [%]; **SF**= Source Fragmentation [V]; **DDM**: Data dependent Mode (cycle time in seconds, CT/[s] or number of scans, NS); **NS**= Number of data dependent scans

#### Search Settings

| | |
|---|---|
| Program & version | MaxQuant v2.0.2.0 |
| Search engine | Andromeda |
| settings | Basically default; LFQ and MBR were turned on |
| Static modification | none |
| Digestion mode | unspecific |
| | Min. peptide length for unspecific search 6<br>Max. peptide length for unspecific search 36 |
| Dynamic modification | Acetyl (N-term); Oxidation (M) |
| Modification included in quantification | Oxidation (M) |
| Databases | 1. Contaminants<br>2. ACE_0699_UP000000625_83333.fasta |
| Annotation | |

### ACE_0809-03

#### File legend

| ACE ID | Alternate ID | Organ/ cell line | Treatment/ experimental setup |
|---|---|---|---|
| ACE_0809-03_BH01 | MS15_H_0 | Human (overexpressed in *E. coli*) | 2 µM HTRA1 Control, 0 sec, supernatant after acetone precipitation |
| ACE_0809-03_BH02 | MS15_H_0 | dito | 2 µM HTRA1 Control, 0 sec, supernatant after acetone precipitation |
| ACE_0809-03_BH03 | MS15_H_0 | dito | 2 µM HTRA1 Control, 0 sec, supernatant after acetone precipitation |
| ACE_0809-03_BH04 | MS15_H_0 | dito | 2 µM HTRA1 Control, 0 sec, supernatant after acetone precipitation |
| ACE_0809-03_BH05 | MS15_H_120 | dito | 2 µM HTRA1 Control, 120 min, supernatant after acetone precipitation |
| ACE_0809-03_BH06 | MS15_H_120 | dito | 2 µM HTRA1 Control, 120 min, supernatant after acetone precipitation |
| ACE_0809-03_BH07 | MS15_H_120 | dito | 2 µM HTRA1 Control, 120 min, supernatant after acetone precipitation |
| ACE_0809-03_BH08 | MS15_H_120 | dito | 2 µM HTRA1 Control, 120 min, supernatant after acetone precipitation |
| ACE_0809-03_BH09 | MS15_T_0 | dito | 10 µM Tau Control, 0 sec, supernatant after acetone precipitation |
| ACE_0809-03_BH10 | MS15_T_0 | dito | 10 µM Tau Control, 0 sec, supernatant after acetone precipitation |
| ACE_0809-03_BH11 | MS15_T_0 | dito | 10 µM Tau Control, 0 sec, supernatant after acetone precipitation |
| ACE_0809-03_BH12 | MS15_T_0 | dito | 10 µM Tau Control, 0 sec, supernatant after acetone precipitation |
| ACE_0809-03_BH13 | MS15_T_120 | dito | 10 µM Tau Control, 120 min, supernatant after acetone precipitation |
| ACE_0809-03_BH14 | MS15_T_120 | dito | 10 µM Tau Control, 120 min, supernatant after acetone precipitation |
| ACE_0809-03_BH15 | MS15_T_120 | dito | 10 µM Tau Control, 120 min, supernatant after acetone precipitation |
| ACE_0809-03_BH16 | MS15_T_120 | dito | 10 µM Tau Control, 120 min, supernatant after acetone precipitation |
| ACE_0809-03_BH17 | MS15_TH_0 | dito | 10 µM Tau + 2 µM HTRA1, 0 sec, supernatant after acetone precipitation |
| ACE_0809-03_BH18 | MS15_TH_0 | dito | 10 µM Tau + 2 µM HTRA1, 0 sec, supernatant after acetone precipitation |
| ACE_0809-03_BH19 | MS15_TH_0 | dito | 10 µM Tau + 2 µM HTRA1, 0 sec, supernatant after acetone precipitation |
| ACE_0809-03_BH20 | MS15_TH_0 | dito | 10 µM Tau + 2 µM HTRA1, 0 sec, supernatant after acetone precipitation |
| ACE_0809-03_BH21 | MS15_TH_15 | dito | 10 µM Tau + 2 µM HTRA1, 15 sec, supernatant after acetone precipitation |
| ACE_0809-03_BH22 | MS15_TH_15 | dito | 10 µM Tau + 2 µM HTRA1, 15 sec, supernatant after acetone precipitation |
| ACE_0809-03_BH23 | MS15_TH_15 | dito | 10 µM Tau + 2 µM HTRA1, 15 sec, supernatant after acetone precipitation |
| ACE_0809-03_BH24 | MS15_TH_15 | dito | 10 µM Tau + 2 µM HTRA1, 15 sec, supernatant after acetone precipitation |
| ACE_0809-03_BH25 | MS15_TH_1 | dito | 10 µM Tau + 2 µM HTRA1, 1 min, supernatant after acetone precipitation |
| ACE_0809-03_BH26 | MS15_TH_1 | dito | 10 µM Tau + 2 µM HTRA1, 1 min, supernatant after acetone precipitation |
| ACE_0809-03_BH27 | MS15_TH_1 | dito | 10 µM Tau + 2 µM HTRA1, 1 min, supernatant after acetone precipitation |
| ACE_0809-03_BH28 | MS15_TH_1 | dito | 10 µM Tau + 2 µM HTRA1, 1 min, supernatant after acetone precipitation |
| ACE_0809-03_BH29 | MS15_TH_5 | dito | 10 µM Tau + 2 µM HTRA1, 5 min, supernatant after acetone precipitation |
| ACE_0809-03_BH30 | MS15_TH_5 | dito | 10 µM Tau + 2 µM HTRA1, 5 min, supernatant after acetone precipitation |
| ACE_0809-03_BH31 | MS15_TH_5 | dito | 10 µM Tau + 2 µM HTRA1, 5 min, supernatant after acetone precipitation |
| ACE_0809-03_BH32 | MS15_TH_5 | dito | 10 µM Tau + 2 µM HTRA1, 5 min, supernatant after acetone precipitation |
| ACE_0809-03_BH33 | MS15_TH_10 | dito | 10 µM Tau + 2 µM HTRA1, 10 min, supernatant after acetone precipitation |
| ACE_0809-03_BH34 | MS15_TH_10 | dito | 10 µM Tau + 2 µM HTRA1, 10 min, supernatant after acetone precipitation |
| ACE_0809-03_BH35 | MS15_TH_10 | dito | 10 µM Tau + 2 µM HTRA1, 10 min, supernatant after acetone precipitation |
| ACE_0809-03_BH36 | MS15_TH_10 | dito | 10 µM Tau + 2 µM HTRA1, 10 min, supernatant after acetone precipitation |
| ACE_0809-03_BH37 | MS15_TH_2 | dito | 10 µM Tau + 2 µM HTRA1, 2 min, supernatant after acetone precipitation |
| ACE_0809-03_BH38 | MS15_TH_2 | dito | 10 µM Tau + 2 µM HTRA1, 2 min, supernatant after acetone precipitation |

| | | | |
|---|---|---|---|
| ACE_0809-03_BH39 | MS15_TH_2 | dito | 10 µM Tau + 2 µM HTRA1, 2 min, supernatant after acetone precipitation |
| ACE_0809-03_BH40 | MS15_TH_2 | dito | 10 µM Tau + 2 µM HTRA1, 2 min, supernatant after acetone precipitation |
| ACE_0809-03_BH41 | MS15_TH_120 | dito | 10 µM Tau + 2 µM HTRA1, 120 min, supernatant after acetone precipitation |
| ACE_0809-03_BH42 | MS15_TH_120 | dito | 10 µM Tau + 2 µM HTRA1, 120 min, supernatant after acetone precipitation |
| ACE_0809-03_BH43 | MS15_TH_120 | dito | 10 µM Tau + 2 µM HTRA1, 120 min, supernatant after acetone precipitation |
| ACE_0809-03_BH44 | MS15_TH_120 | dito | 10 µM Tau + 2 µM HTRA1, 120 min, supernatant after acetone precipitation |

#### LC_Settings

| | |
|---|---|
| MS device | Orbitrap Fusion Lumos |
| LC device | Evosep One |
| ion source | Thermo Nanospray Flex |
| **Analytical column** | EV1064 Analytical Column – 60 & 100 samples/day |
| column diameter | Length ($L_C$) = 8 cm; ID = 100; OD = PEEK; emitter EV-1086 Stainless steel emitter |
| stationary phase | Dr Maisch C18 AQ, 3 µm beads |
| particle diameter ($d_p$) | 3 µm |
| Pore size | 120 Å |
| Column ID | AC_EVO-008 |
| Column oven | na |
| Column oven temp. | na |
| **solvents** | A: 0.1% FA in UPLC water<br>B: 0.1% FA in UPLC ACN |
| gradient | 21 min gradien (60 SPD) |

#### MS_Settings

| Project | MS | general | MS1 | MS2 | MS2 | MS3 | Comments; special settings |
|---|---|---|---|---|---|---|---|
| ACE_0809 | Lumos | Tune v3.5.3881.18<br>Xcalibur v4.5.445.18<br>Gradient: 21 min | Analyzer: FT<br>Res.: 120000<br>SR: 375 - 1600<br>AGC: Standard<br>AcT: Auto<br>RF: 30<br>SF: --<br>DDM: CT/3sec | Analyzer: IT<br>Res./ScR: -/rapid<br>SR: Auto<br>AGC: Standard<br>AcT: Auto<br>CS: +2 to +6<br>IsM: Q<br>IsW: 1.6<br>Frag.: HCD<br>NCE: 32 | | | classic orbitrap experiment: MS1 in Orbitrap at high resolution and data dependent MS2 in Iontrap at turbo scan rate. Dynamic exclusion enabled (exclude after n times=1; Exclusion duration (s)= 20; mass tolerance= ± 10ppm)<br>intensity threshold: $5 \times 10^3$ |

Note: **FT**= Fourier Transform (Orbitrap); **IT**= Iontrap; **Q**= Quadrupol; **Res.**= max. Resolution at 200 m/z (Lumos) or 400 m/z (Elite) [FWHM (full width at half maximum)]; **ScR**= scan rate for measurements in the IT; **SR**= scan range [m/z]; **AGC**= automatic gain control, max number of acquired ions per measurement; **AcT**= max. Ion acquisition time [ms]; **CS**= charge states used for fragmentation; **IsM**= Isolation mode (Q or IT), MS2 isolation and further is only done in IT; **IsW**= Isolation window [m/z], value followed by scan mode the isolation is based on (MS1, MS2 …) **Frag.**= Fragmentation method; **HCD**= Higher-energy collisional dissociation; **CID**= Collision-induced dissociation; **ETD**= Electron-transfer dissociation**; EThcD=** Electron-Transfer/Higher-Energy Collision Dissociation; **sHCD**= stepped HCD**; NCE**= normalized collision energy; **cycles**: number of MSn recorded or max cycle time; RF= RF Lens [%]; **SF**= Source Fragmentation [V]; **DDM**: Data dependent Mode (cycle time in seconds, CT/[s] or number of scans, NS); **NS**= Number of data dependent scans

#### Search Settings

| | |
|---|---|
| Program & version | MaxQuant v2.0.3.0 |
| Search engine | Andromeda |
| settings | Basically default; LFQ and MBR were turned on |
| Static modification | none |
| Digestion mode | unspecific |
| | Min. peptide length for unspecific search 6<br>Max. peptide length for unspecific search 36 |
| Dynamic modification | Acetyl (N-term); Oxidation (M); Phospho (STY) |
| Modification included in quantification | Oxidation (M);Acetyl (Protein N-term) |
| Databases | 3. Contaminants<br>1. ACE_0809_SOI_v01.fasta |
| Annotation | |

#### Supplemental data 6

#### Example for how UMSAP calculates the relative frequency of cleavages

In a MS experiment the following peptides were identified. They all share the same P1 site at V231

```
A -  RGHYV
B -   GHYV
C -       DFRTGH
D -       DFRTGHKL
E -       DFRTGHKLM
F -       DFRTGHKLMT
```

**Average intensities.** 0 values mean that the peptide was not detected in the given time point or the intensity values are not significantly different to the control experiments.

| Time Point | A | B | C | D | E | F |
|---|---|---|---|---|---|---|
| 5 min | 2 | 0 | 0 | 3 | 3 | 10 |
| 15 min | 4 | 1 | 0 | 0 | 6 | 5 |
| 30 min | 8 | 3 | 5 | 0 | 0 | 1 |

The first non-zero average intensity along the time points is taken as reference for the calculation of the relative intensity.

| | Relative Intensities | | | | | | Relative cleavage frequency |
|---|---|---|---|---|---|---|---|
| Time-point | A | B | C | D | E | F | P1 (V231) |
| 5 min | 2/2 = 1 | 0/1 = 0 | 0/5 = 0 | 3/3 = 1 | 3/3 = 1 | 10/10 = 1 | 1+0+0+1+1+1 = 4 |
| 15 min | 4/2 = 2 | 1/1 = 1 | 0/5 = 0 | 0/3 = 0 | 6/3 = 2 | 5/10 = 0.5 | 2+1+0+0+2+0.5 = 5.5 |
| 30 min | 8/2 = 4 | 3/1 = 1 | 5/5 = 1 | 0/3 = 0 | 0/3 = 0 | 1/10 = 0.1 | 4+1+1+0+0+0.1 = 6.1 |
